# Both L-lactyl and D-lactyl enantiomers modify histones in mouse testis

**DOI:** 10.1101/2025.10.09.681385

**Authors:** Julie Manessier, Hassan Hijazi, Sabine Brugière, Marie Courçon, Christophe Masselon, Alberto de la Iglesia, Julie Cocquet, Delphine Pflieger

## Abstract

Dynamic histone post-translational modifications are crucial to precisely orchestrate gene expression programs. The recently discovered histone lysine lactylation has already been explored in various pathological contexts, but less in normal tissues. This modification exists as two enantiomers, L- and D-lactylation; the first one may more likely modify histones due to abundant L-lactate produced by glycolysis. Here we report the identification by proteomics of L- and D-lactylation on lysines of histones H3 and H4 in mouse testis. We developed a targeted proteomic analysis of histone peptides using synthetic sequences modified by L- or D-lactyl, to acquire reliable identification and quantification data. Some histone peptides bearing either enantiomer are separated by reversed-phase chromatography. Interestingly, while several lysines of H3 and H4 exhibit a balanced amount of both enantiomers, residues 18 and 79 from histone H3 (H3K18 and H3K79) appear to be significantly more D-lactylated, while H3K23 is more L-lactylated. The stoichiometry of lactylation is low over the whole sequence of H3 and H4, representing 0.01 to 0.5%, which contrasts with acetylation whose abundance substantially varies between lysines. Interestingly, lactylation is more abundant than acetylation on the C-terminal half of H3 and H4. These results suggest a mechanism producing a mixture of the two enantiomers of lactate, or of a more direct substrate for lactylation, and supporting a site-selective addition of the enantiomers.

## Introduction

Gene expression is subtly regulated by the dynamics of histone post-translational modifications (PTMs), the diversity of which has dramatically expanded with the successive discovery of various structures modifying lysine residues, that resemble acetylation but vary in length, hydrophobicity and charge. These new acylations are progressively described to be endowed with specific functions compared to the canonical mark acetylation. These PTMs are likely linked to the corresponding acyl-CoA metabolites, which establishes an intricate link between metabolism and epigenetics regulatory mechanisms. In 2019, lactylation was discovered to regulate gene expression in macrophages and upon hypoxia^1^. It was described to be induced by L-lactate, a metabolite produced by glycolysis via the enzyme Lactate DeHydrogenase A (LDHA), which can be very abundant in cell types heavily resorting to this energy producing cycle, such as cancer cells. Yet, further publications supported two scenarios to explain the addition of lactylation to histones, and beyond, to non-histone proteins. The first one is enzymatic and uses L-lactate as indirect substrate^1^, whereas the second one hypothesizes a non-enzymatic addition of lactylation from D-lactoyl-glutathione^2^.

Indeed, on the one hand, several studies described the transfer of a lactyl moiety to protein lysines by enzymes of the Histone AcetylTransferase (HAT) families, or, more astonishingly, via other metabolic enzymes. In the original description of histone lactylation, Zhang *et al*. demonstrated that lactyl-CoA could serve as the substrate for histone lactylation in chromatin template-based histone modification and transcription assays *in vitro* involving p300 as the lactyltransferase^1^. Whether this mechanism holds true *in vivo* could remain questionable, due to the very low abundance of lactyl-CoA in mammalian tissues^*3*^. Recently, Alanyl-tRNA synthetase, AARS1, was shown to couple lactate with AMP and carry out the addition of a lactyl group to lysines of target proteins, including the YAP-TEAD complex in gastric cancer cells^4^ and the tumor-suppressor p53^5^. Shortly after, both AARS1 and AARS2 were described to constitute lactyltransferases capable of modifying numerous peritoneal macrophage proteins, and in particular the cyclic GMP-AMP synthase (cGAS), thus reducing its activity and disrupting its binding to DNA^6^. The latter study also described lactate-AMP as the reaction intermediate, and that depleting both AARS1 and AARS2 led to the near complete clearing of the cellular lactylome in macrophages. Besides, HBO1, a member of the MYST family of HATs, was demonstrated to catalyze lactylation of histones, particularly at lysine 9 from histone H3 (H3K9), and of non-histone proteins in HeLa cells^7^. Yet another mechanism was described wherein ACSS2 translocates to the nucleus upon EGFR activation in glioblastoma, and associates with KAT2A to catalyze lactylation of H3K14 and H3K18^8^. Finally, guanosine triphosphate-specific succinyl-CoA synthetase (GTPSCS) was shown to relocalize to the nucleus and associate to p300 to form a lactyltransferase complex, targeting mainly H3K18 in various cultured and glioma cells^9^. In all, the mechanisms leading to enzymatic histone lactylation and in particular the enzymes responsible for producing the substrate for lactylation (lactyl-CoA or lactate-AMP) and its transfer to histone and non-histone lysine residues may depend on the considered cell type, which determines the cellular abundance of these enzymes^9^.

On the other hand, Gaffney *et al*. described in 2020 the non-enzymatic addition of D-lactyl, mostly on non-histone proteins, with D-lactoylglutathione (LGSH) being the substrate for this modification^2^. The two major enzymes involved in regulating this PTM are GLO1 and GLO2, with GLO1 converting the glycolytic by-product methylglyoxal (MGO) into lactoylglutathione, and GLO2 regenerating glutathione and D-lactate from the latter metabolite. GLO1 and 2 play an important role in detoxifying toxic MGO which reacts with Cys, Arg and secondarily with Lys residues to produce glycated products. The same laboratory then pursued their inquiry by demonstrating *in vitro* that lactyl-CoA led more readily to histone lactylation than LGSH, but that LGSH efficiently reacted with CoA to generate lactyl-CoA, thus constituting an important indirect source of non-enzymatic protein lactylation. In addition, these authors established that macrophages depleted of GLO2 exhibited increased intracellular levels of LGSH and lactyl-CoA, but not of lactate^10^. Another group showed that D-lactate induced H3K18 lactylation and NF-kB signaling pathway in bovine mammary epithelial cells subjected to lipopolysaccharides ^11^.

Many of the above studies describing L-lactylation were performed on cancer cells and tumors, characterized by remarkably high intracellular levels of L-lactate derived from exacerbated aerobic glycolysis, a phenomenon called the Warburg effect^9^. In other cellular contexts, whether L- or D-lactate actually contributes to histone lysine lactylation was usually not assessed. This question is worth asking in physiological conditions, namely in non-diseased tissues. Another important aspect to investigate is the relative abundance of lactylation compared to acetylation. To the best of our knowledge, this information has never been assessed in reports on histone lactylation.

Here, we carefully dissected the lactylation pattern of histones H3 and H4 extracted from mouse testis. This organ and more specifically spermatogenesis constitute a valuable model to study the functional roles of histone PTMs, because the differentiation program of male gametes into spermatozoa is characterized by dramatic chromatin rearrangements and waves of histone PTM changes, including acetylation^12^, crotonylation^13^ and butyrylation^14^. Lactylation has started being studied in this organ, considering that differentiating male germ cells rely on lactate as a major energy source^15,16^. These studies exclusively relied on antibody-based experiments targeting selected N-terminal lysines of H3 and H4^17^, and H4K8lac was more specifically demonstrated to mark the promoters of genes involved in meiosis and to be associated with recombination hotspots^18^. By quantitative proteomics using heavy labelled synthetic peptides modified with L- or D-lactylation, we demonstrate on several lysine residues from H3 and H4 that the two enantiomers co-exist. We also estimate that the relative stoichiometry of lactylation is constantly low, ranging between 0.01 and 0.5%, over the whole sequence of both histones. This is in stark contrast with the stoichiometries of acetylation, which can be very abundant on the N-terminal tails (e.g. at H3K14, H3K23 and H4K16), and appear to be of lower abundance than lactylation at H3K56, H3K64, H3K79, H3K122 and H4K77. In all, our findings prompt us to reconsider the mechanism by which histone lysines become lactylated, to account for the addition of both enantiomers.

## Material and Methods

### Materials

All mouse testes used came from mice of the C57BL/6 genotype and aged between two and three months. Procedures were subjected to local ethical review (Comité d’Ethique pour l’Experimentation Animale, Université Paris Descartes, registration number CEEA34.JC.114.12, APAFIS 14214-2017072510448522v26). Synthetic peptides were purchased from JPT Peptide Technologies, their sequences and modifications are listed in **Suppl. Table 1**. TEAB (ref. T7408), sulfuric acid (ref. 339741), tricholoroacetic acid (TCA, ref. T0699), propionic anhydride (ref. 240311), hydroxilamine (ref. 438227) and NH_4_OH (ref. 221228) were purchased from Sigma.

### Acid extraction of histones from mouse testes

Histones were extracted from mouse testis nuclei using the following protocol, derived from ^19^. Briefly, each testis was homogenized in 1 mL of Nuclei Isolation Buffer (NIB; 15 mM Tris-HCl (pH 7.5), 60 mM KCl, 11 mM CaCl2, 5 mM MgCl2, 250 mM sucrose, 1 mM DTT, 10 mM sodium butyrate, 3 µM trichostatin A, 50 mM nicotinamide, complete mini protease inhibitor cocktail from Roche, and 0.3% final (v/v) NP-40) using a Dounce homogenizer. Lysates containing nuclei were centrifuged at 1000g, 4°C for 5 min. The pelleted nuclei were then resuspended in NIB without NP-40 and centrifuged at 1000g, 4°C for 5 min. This wash was repeated a second time. The nuclei were resuspended in 450 µL of 0.2 M H_2_SO_4_ and rotated at 4°C for 1h, then centrifuged at 4°C at maximum speed for 5 min. Histone-containing supernatants were collected in new tubes and precipitated by adding pure TCA (final concentration 20%) and incubating for 2h in ice, then centrifuged at 20000g, 4°C for 15 min. The supernatant was aspirated and the precipitated histones were coated with 0.1% HCl in freezer-cold acetone, then centrifuged at 4°C for 5 min at maximum speed. This step was repeated once with non-acidified acetone. After removing the acetone, samples were dried under the hood. Finally, histone samples were resuspended in 100 µL of Milli-Q water and stored at -20°C. An aliquot of histones was separated by SDS-PAGE (4-12%) to verify their purity and estimate the total amount (approximately 200 µg of histones were extracted per testis). For the gel-based analysis, the bands corresponding to histones H3 and H4 were cut with a scalpel and stored at -20°C until being processed.

### In vitro propionylation and tryptic digestion of histones

For each step of primary amine derivatization by propionyl, two propionic anhydride solutions were extemporaneously prepared as follows: a first dilution of pure propionic anhydride diluted at 1:100 in water, followed by a second dilution of this solution in 1 mL of 100 mM TEAB (pH 8.5). In-gel and in-solution derivatization procedures share most steps. Histone-containing gel pieces were first destained. For derivatization, 100 µL of propionic anhydride solution diluted in TEAB were added to each gel band for 30 min. This step was repeated a second time, after removing the first solution and drying the gel pieces in pure acetonitrile (ACN). Histones in solution were propionylated in two rounds as above, separated by a drying of samples in a speedvac. In both approaches, histones were then digested with trypsin diluted in 50 mM TEAB overnight at 37°C, at an enzyme:substrate mass ratio of 1:20. After digestion, peptides were extracted from the gel pieces. Another propionylation step was performed to label the N-terminal ends of peptides, as previously performed on intact histones. To remove non-specific propionylated sites (typically at Ser/Thr residues), a reverse propionylation step was carried out by applying 20 µL of 500 mM hydroxylamine and 6 µL of ammonium hydroxide (pH 12) for 20 min at room temperature with stirring (600 rpm)^20–22^. The reaction was stopped by adding a few drops of pure trifluoroacetic acid (TFA). Samples were dried in a SpeedVac and desalted using Affinisep SPE cartridges (BioSPE PurePrep).

### Preparation of synthetic peptides

The synthetic heavy peptides modified by either L-lactylation, D-lactylation or acetylation were verified by LC-MS/MS analysis in terms of correct sequence, modification site and absence of light counterparts. To test the chromatographic behavior of D- and L-lactylated peptides, three different mixtures of these synthetic peptides were prepared, in molar ratios 1:1, 1:2 and 2:1, respectively. These samples were propionylated as described above, followed by a reverse propionylation step. The peptides were then dried and desalted, before being analyzed by LC-MS/MS on a C18 Aurora column. To assess the presence of L- and then also D-lactylation in mouse testis histones, endogenous samples were successively spiked with two mixtures of synthetic peptides. A first mixture was made up of all L-lactylated and acetylated peptides and a second mixture of L-lactylated, acetylated and D-lactylated peptides. About 0.5 µg of endogenous histone peptides and 50 fmol of each synthetic peptide were injected each time.

### Liquid chromatography-mass spectrometry analysis

Peptides were loaded onto a PepMap C18 pre-column (300 μm × 5 mm, Thermo Fisher Scientific) with 0.1% formic acid. The peptides were then separated on a reverse-phase capillary column (Aurora C18, 75 µm x 25 cm, 1.7 µm beads, 120 Å particles) on a Nano Ultimate 3000 system (Thermo Fisher Scientific) coupled to an Exactive HF mass spectrometer (Thermo Fisher Scientific). The mobile phases consisted of solvent A (water + 0.1% formic acid) and solvent B (ACN with 0.08% (v/v) formic acid). The peptides were separated by applying a gradient consisting of an increase from 2% to 7% of B in 5 min, then 7% to 31% of B in 55 min, then 31% to 41% B in 8 min, followed by column wash at 72% B for 9 min and column re-equilibration at 2% B for 14 min.

For analyses in Data-Dependent Acquisition (DDA) mode, MS1 spectra were acquired at a resolution of 60,000 with an AGC of 1e6, over an m/z range from 300 to 1300. MS2 spectra were acquired at a resolution of 15,000 with an AGC of 2e5, with a maximum injection time of 100 ms, a loop count of 20 and an isolation window of 1.5. Peptides were isolated for fragmentation using higher-energy collisional dissociation (HCD) with a collision energy of 30 and a dynamic exclusion of 10 s. The first fixed mass of MS/MS spectra was set at m/z 80, to be able to detect the cyclic immonium (CycIm) ion produced by monomethylated lysines (at m/z 98.09). For analyses by Parallel Reaction Monitoring (PRM), MS2 spectra were acquired at a resolution of 30,000, with an AGC of 1e6, a maximum injection time of 120 ms and an isolation window of 1.6.

Scheduled PRM analyses were designed to target (i) L-lactylated and acetylated peptides to start studying the presence of L-lactylation in mouse testis histones (**Suppl. Table 2**), (ii) L-, D-lactylated and acetylated peptides to determine the relative abundances of L- and D-lactylation (L/D), as well as of acetylation versus lactylation (ac/lac) (**Suppl. Table 3**). In the latter case, we included the targeting of oxidized forms of methionine-containing peptides H3 VTIMPK122DIQLAR and H4 K79TVTAMDVVYALK91R. Finally, to be able to get an estimate of the stoichiometry of L- and/or D-lactylation as a percentage of each H3 and H4 lysine residue, we determined the stoichiometry of each corresponding acetylation site. DDA analyses were relevant to estimate these values for N-terminal lysines where this PTM is quite abundant **(Suppl. Table 4**), whereas PRM analyses were designed to systematically characterize lysines from the histone fold domain (**Suppl. Table 5**).

### Pre-processing, identification and quantification of modified peptides from DDA and PRM analyses

For DDA analyses, the acquired RAW data were converted to .MGF files by the in-house developed tool MGFBoost (article in preparation; https://www.profiproteomics.fr/proline/other-tools/), which maintains the relative intensities of fragments in MS/MS spectra^21^. Identification of histone peptides was obtained with the Mascot search engine (Matrix Science v2.8) using an in-house histone database complemented with a list of classical contaminants^23^. The search parameters were as follows: Arg-C as the enzyme, Propionyl (N-term) as a fixed peptide modification, acetyl (K), butyryl (K) standing for the dual modification by endogenous monomethyl and chemical propionyl, dimethyl (K), trimethyl (K), lactyl (K), propionyl (K), oxidation (Q), and oxidation (M) as variable modifications. Peptide mass tolerance was set at 5 ppm and fragment mass tolerance at 20 ppm. For PRM analyses, .RAW files were imported directly into Skyline software and matched to Mascot .DAT identification files. Each peptide identification was visually inspected in Mascot, in particular to verify the presence of intense cyclic immonium ions corresponding to the modified lysine in first position^21^. Peptide quantification was calculated from PRM analyses based on specific y-type fragments indicated in **Suppl. Tables 3 and 5**, while considering the area under the curve.

### Experimental Design and Statistical Rationale

Samples of synthetic D- and L-lactylated peptides prepared in 1:1, 1:2 and 2:1 ratios were analyzed by LC-MS/MS in technical duplicates which are both shown in **Suppl. Figures section 3**. Histones from mouse testes were obtained from four mice. These four biological replicates were analyzed to estimate the relative abundances of L- and D-lactylation, as well as acetylation. Of note, when quantifying L-versus D-lactyl and lactyl versus acetyl at the same lysine sites, we compared mass spectrometry signals within a single PRM analysis, so that normalization of total histone amounts between samples was non-necessary.

## Results

### Identification by proteomics of potentially lactylated sites on histones H3 and H4 from mouse testis

We sought to obtain a global mapping of tentative lactylation sites on histones H3 and H4 from mouse testis, by following the workflow depicted in **Figure 1a**. After enrichment of nuclei from the tissues, histones were acid-extracted and then either separated by SDS-PAGE or directly processed in solution by the propionylation of endogenously non-modified lysines, digested by trypsin, treated by a second step of propionylation of peptide N-termini, and lastly with a basic solution to remove excessive propionylation at S/T/Y residues ^20,22^. Interpretation by the program Mascot of the proteomics data acquired on these samples by liquid chromatography-tandem mass spectrometry (LC-MS/MS) proposed the identification of a number of peptides bearing a precise mass increment of 72.021 u, which might correspond to lactylation. We carefully scrutinized each MS/MS spectrum matched to a lactylated peptide from histones H3 and H4, by checking the following criteria. If the lactylation site was attributed to the lysine in the first position of the peptide, we demanded that the b1 fragment and the CycIm ion corresponding to lactylated lysine (Kla) be detected with high intensity, as formerly reported ^21,24–26^ (**Figure 1b**). If lactylation was attributed to a lysine residue inside the peptide sequence, we verified the presence of y fragments directly surrounding the Kla site (**Figure 1b**). We finally established the global mapping of tentative lactylation sites on histones H3 and H4, and compared it to the sites of acetylation identified from the same LC-MS/MS analyses (**Figure 1c and MS/MS spectra in Suppl. Figures, section 1**). Interestingly, possible lactylation sites were detected on H3K56, H3K64, H3K122 as well as H4K77 and H4K91, whereas the acetylated counterparts were not identified. Of note, the latter lysines are known to be acetylated at very low stoichiometry, which likely explains the difficulty to identify them in exploratory (DDA) LC-MS/MS analyses, in which peptide ions are selected for fragmentation based on their MS1 signal intensity.

**Figure 1:**
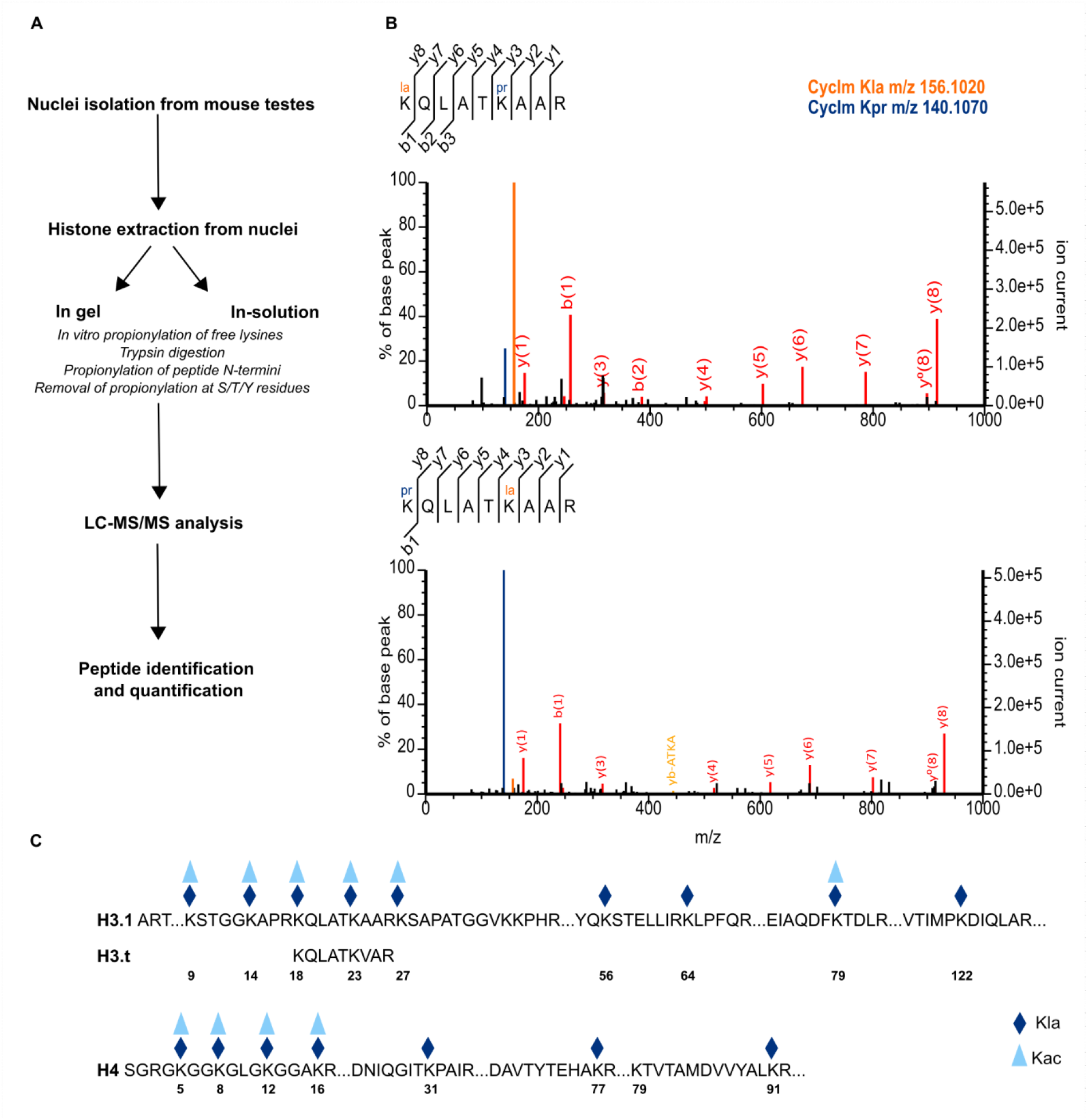
Establishment of a mapping of tentative lactylation sites on histones H3 and H4 from mouse testis. (A) Workflow followed to identify lactylation sites by DDA analysis. (B) Validation of the presence of a tentative lactylation (mass increment of 72.021 u) at lysines H3K18 and H3K23, by detection of the cyclic immonium ion expected for Kla at m/z 156.102 and of discriminating y fragments. (C) Global mapping of tentative lactylation and acetylation sites over the sequences of H3 and H4.

### False identifications of presumably lactylated peptides correspond to propionylated and oxidized peptides, which can be of prevailing abundance

When visually inspecting the MS/MS spectra attributed by the program Mascot to lactylated peptides, we rejected some identifications because they did not match the above validation criteria. Visual sequencing of these spectra usually confirmed the proposed amino acid sequence, yet the absence of the CycIm ion for Kla refuted the lactylated nature of the fragmented peptide. **Figure 2a** shows an MS/MS spectrum presumably identifying peptide K_18_laQLATK_23_prAAR from H3. Yet, the dominant fragment at m/z 140.10 corresponding to the CycIm ion of Kpr rather reveals the first lysine is propionylated, and then encourages to position the mass difference between lactyl and propionyl at an upcoming residue. Considering that this deltamass is 15.9949 u and matches an oxidation (ox), we re-interpreted our datasets with the possible oxidation of several residues, including glutamine. This led to the considered spectrum be interpreted as H3 K_18_prQoxLATK_23_prAAR, and specifically allowed interpreting the two highest-mass fragments as y/y_0_ ions (**Figure 2b**). Extraction of the chromatographic peaks of the peptides of very same mass H3 K_18_laQLATK_23_prAAR, K_18_prQLATK_23_laAAR and K_18_prQoxLATK_23_prAAR revealed a low, hardly quantifiable, MS signal for the two first peptides at very similar retention times (RT, about 46.7 min) and a much more intense MS signal for the oxidized peptide species at RT about 49 min (**Figure 2c)**. The oxidized peptide comes from H3 molecules which are endogenously non-modified at H3K18 and H3K23 and constitute the most abundant proteoform.

**Figure 2:**
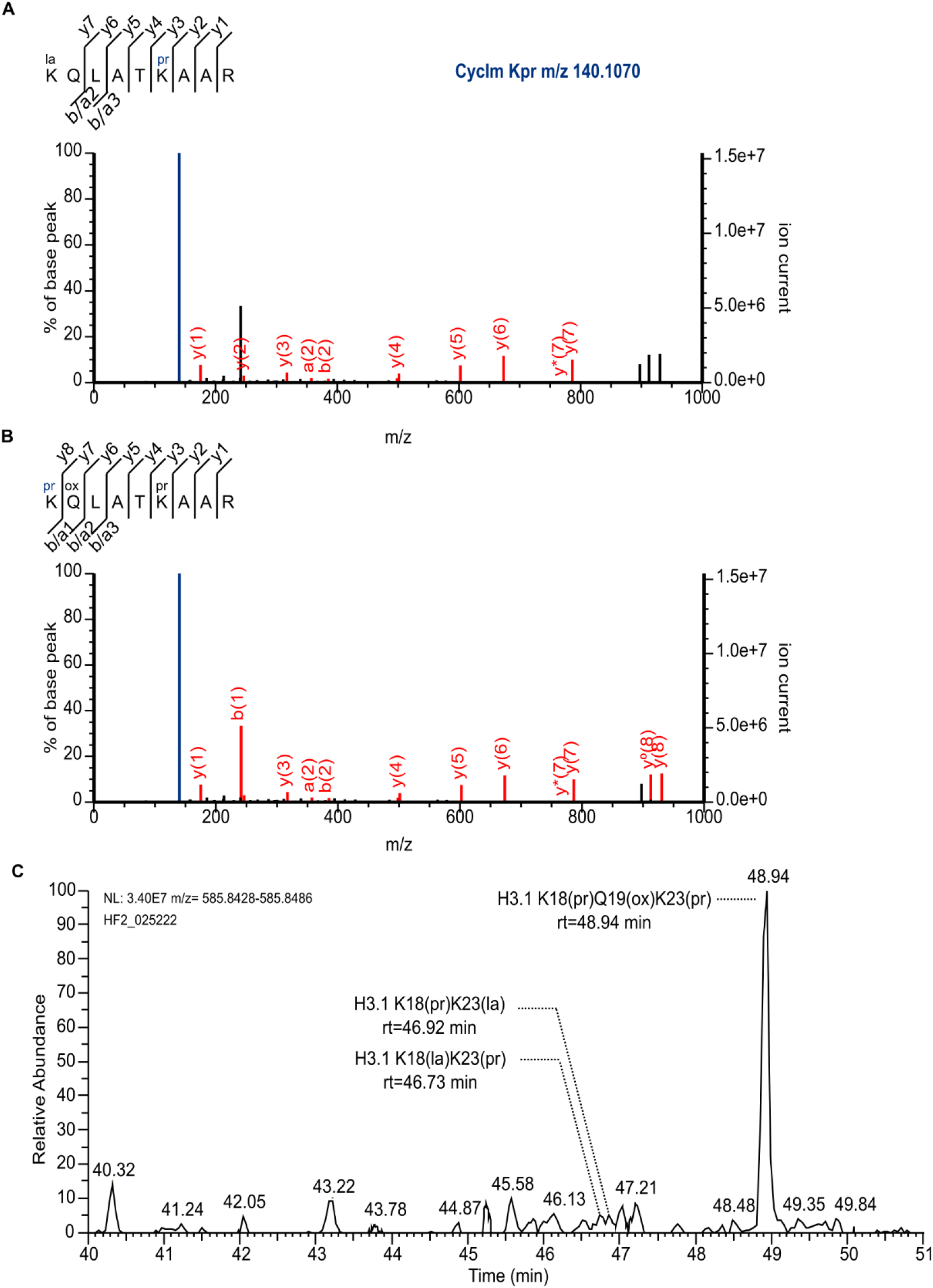
Oxidized peptides lead to erroneous identifications of lactylated sequences. (A) MS/MS spectrum tentatively matched by Mascot to H3 K18QLATK23AAR lactylated on the first lysine. The presence of an intense CycIm ion at m/z 140.10 that signs H3K18pr, and the absence of CycIm ion for Kla (expected at m/z 156.10) and of b1, lead to rejecting this theoretical sequence. (B) Taking into account the oxidation of glutamine leads to the correct identification of H3 KprQoxLATKprAAR. (C) Relative abundances estimated by their MS1 XICs of H3 KQLATKAAR lactylated at either K18 or K23, and of KprQoxLATKprAAR.

Comparing the MS intensities of K_18_prQoxLATK_23_prAAR and K_18_prQLATK_23_prAAR indicated that oxidation occurs at about 1.5% (**Suppl. Table 7**). Oxidation events likely occur during the harsh treatments applied to histones, at the time of their precipitation in TCA and during the reverse propionylation step at pH 12.

Histone peptides modified by propionylation and an oxidation on reactive residues can thus lead to the erroneous identification of pseudo-lactylated peptides, and their abundance sometimes largely surpasses the one of tentatively lactylated peptides which really bear a modification of 72.021 u on a specific lysine. To more widely assess this phenomenon, we considered other peptides from histones H3 and H4 and could confirm oxidation events at residues M (singly and doubly oxidized), Q, T, Y and H (see **Suppl. Figures, section 2**). All these species contribute to the multitude of (histone variant × modifications) combinations that share the very same mass, and then require careful inspection of their fragmentation spectra to identify the correct sequences^21^.

### Validating the presence of L-lactylation on H3K18 in mouse testis histones and the necessity to also envision D-lactylation

Beyond the impact on identification, the presence of propionylated and oxidized peptides in the samples hinders quantification of tentatively lactylated peptides based on their MS1 signal. To definitely validate the nature of lactylated peptides and reliably extract their corresponding MS signal, it appeared necessary to synthesize lactylated peptides containing a C-terminal arginine labeled with naturally stable heavy atoms, namely ^13^C and ^15^N (**Suppl. Table 1**). These synthetic peptides, propionylated and then spiked into the endogenous histone samples, would strictly elute and fragment identically with their endogenous counterparts if the latter are truly lactylated. Because histones have been described to be modified by the L-lactyl enantiomer rather than the D-lactyl one, we started with assessing this hypothesis ^1,27^.

Mouse testis histones were spiked with heavy labeled L-lactylated synthetic peptides covering a range of lysine residues from histones H3 and H4, and analyzed by LC-MS/MS, by DDA and then PRM (**Suppl. Table 2**). Because H3K18la has been described and functionally characterized in various contexts ^1,8,9,11,28,29^, we started with scrutinizing peptide K_18_laQLATK_23_prAAR and its positional isomer K_18_prQLATK_23_laAAR. Extraction of the chromatographic peaks detected in MS1 for the synthetic and the endogenous peptides (m/z = 590.8498, 2+ and m/z = 585.8457, 2+, respectively) showed a good retention time match between the light and heavy peptides modified on either H3K18 or H3K23 (**Figure 3a**). This observation suggests the presence of both endogenous peptides L-lactylated on H3K18 and H3K23. To fully validate the identity of the endogenous species, we examined the MS/MS spectra acquired on the light/heavy pairs within the two extracted chromatographic peaks. In the first peak, MS/MS spectra acquired on the light and heavy species were identical, except for the 10.008 mass shift due to the ^13^C/^15^N labeling, which validated the presence of peptide K_18_prQLATK_23_(L-la)AAR in mouse testis histones (**Figure 3b**). In contrast, in the second chromatographic peak, the MS/MS spectrum yielded by the light peptide corresponded to a composite spectrum generated by peptides lactylated at either H3K18 or H3K23 (**Figure 3b**). This allowed us to conclude that mouse testis H3 was indeed L-lactylated at residue H3K18, but another isobaric structure ‘mod’ was also present on peptide H3 K_18_prQLATK_23_modAAR, which co-eluted with peptide K_18_(L-la)QLATK_23_prAAR. We then hypothesized that D-lactylation could stand for this modification, thus corresponding to peptide H3 K_18_prQLATK_23_(D-la)AAR.

**Figure 3:**
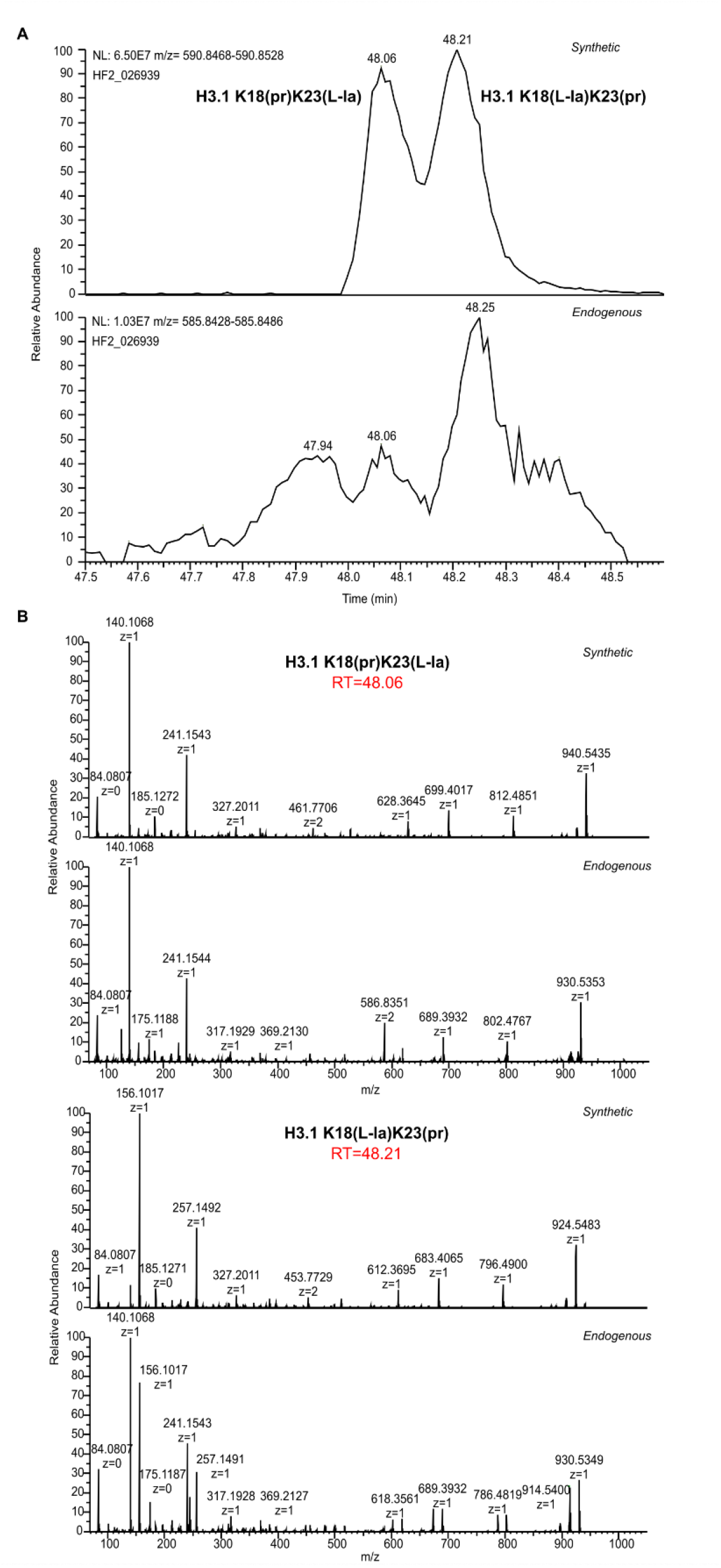
Assessing the presence of L-lactylation at lysines H3K18 and H3K23 in histones from mouse testis. (A) MS1 signal detected for the heavy-labeled peptides of sequences H3 K18(L-la)QLATK23prAAR and H3 K18prQLATK23(L-la)AAR (top panel) and corresponding MS1 signal in endogenous (light) mouse testis histones. (B) MS/MS spectra acquired on the pairs of heavy/endogenous peptides at the apex of the two chromatographic peaks defined by the synthetic peptides.

### Determination of the chromatographic behavior of synthetic peptides from histones H3 and H4 bearing L- or D-lactylated lysines

The peptides formerly synthesized with an L-lactylation on their lysine residues and bearing a heavy C-terminal arginine were similarly produced with D-lactylated lysines (**Suppl. Table 1**). Then, to assess whether enantiomeric peptides bearing either lactyl moiety could be chromatographically separated, all synthetic peptides were propionylated, mixed in variable D:L abundance ratios (1:1, 1:2, and 2:1) and analyzed by LC-MS/MS.

The four peptidoforms corresponding to sequence K_18_QLATK_23_AAR from histone H3 yielded three chromatographic peaks (**Figure 4a)**. The fragmentation spectra at the apex of these peaks in the 1:1 mixture allowed identifying successively the modified peptides K18(la)K23(pr), then K18(pr)K23(la), and finally both positional isomers in the third chromatographic peak which exhibited a doubled intensity (**Figure 4b**). The elution order between the D- and L-lactylated peptides could be established from the signal intensities measured in MS in the mixtures of imbalanced ratios: the K18(D-la)K23(pr) form elutes first, followed by K18(pr)K23(L-la), and finally the two positional isomers K18(pr)K23(D-la) and K18(L-la)K23(pr) co-elute. Of note, even though the two latter positional isomers co-elute, they can be distinguished by their fragmentation profiles, which allows considering MS2-based quantification to assess all four peptidoforms.

**Figure 4:**
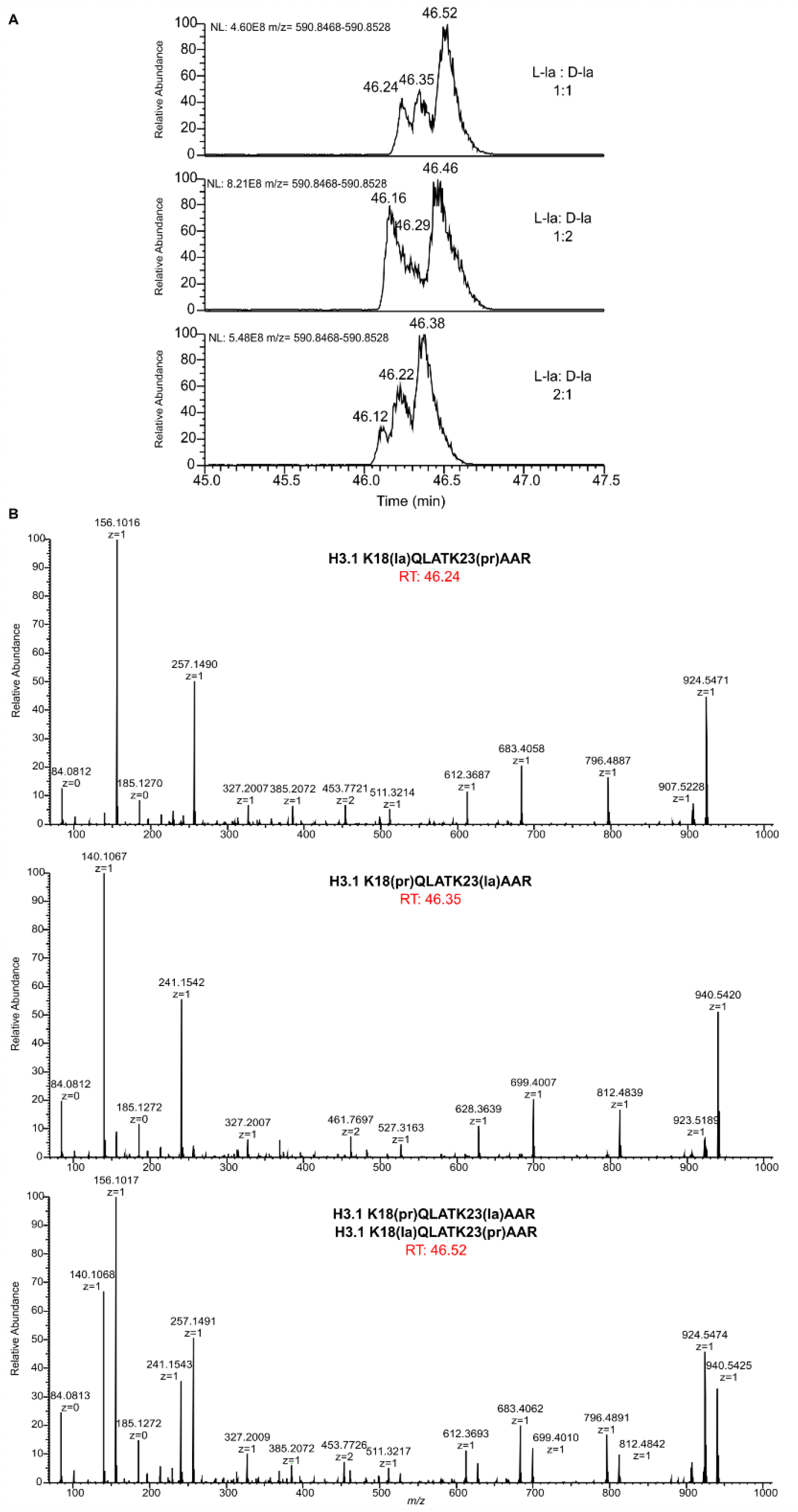
(A) MS1 signals detected when analyzing three mixtures of synthetic peptides H3 K18QLATK23AAR, being L- or D-lactylated at either H3K18 or H3K23, in relative molar ratios 1:1, 1:2 and 2:1. (B) MS/MS spectra acquired at the apex of the three successive chromatographic peaks.

The LC-MS/MS analyses of the three mixtures of L- and D-lactylated peptides revealed that a series of enantiomeric peptides could be chromatographically separated on the C18 Aurora column: these were peptides bearing lactylation on H3K9, H3K18 and H3K23 from both canonical H3 and testis-specific H3 (H3.t, sequence K_18_QLATK_23_<u>V</u>AR), and H3K56 (**Suppl. Figures, section 3**). Enantiomers modified at H3K79 and H4K77 were partially separated. However, no chromatographic separation was obtained of peptides being L- or D-lactylated at H3K14, H3K27 (from both canonical H3 and variant H3.3), H3K64, H3K122, H4K5/K8/K12/K16, H4K31, H4K79 and H4K91. Of note, the peptide bearing the two latter lysines (i.e., K_79_TVTAMDVVYALK_91_R) also contains a methionine that was observed in various oxidation levels, which impacted the chromatographic behavior of D/L-lactylated peptides. Due to the complexity of following this peptide distributed over variable forms, we did not retain it in the following parts of the study.

### Systematic analysis of L- and D-lactylation in histones H3 and H4 from mouse testis and comparison to their acetylated counterparts

To assess the existence of L- and D-lactylation in mouse testis histones, four biological replicates of this sample were spiked with synthetic peptides bearing the two enantiomeric PTMs in a 1:1 ratio, and analyzed by targeted LC-MS/MS (**Suppl. Table 3**). The four peptidoforms of H3 K_18_QLATK_23_AAR could be reliably detected based on discriminative fragments shared between the endogenous and the synthetic peptides (**Figure 5a**), which validated our original assumption that this sequence is modified by L- and D-lactyl on its two lysine residues. We determined that the enantiomers which could be chromatographically resolved had a relative L/D abundance lying between 0.4 and 1.5 (**Figure 5b**). The lowest L/D ratios were measured on H3K18 and the highest on H3K23 (**Suppl. Table 8**). Statistical assessment of MS2 signals indicated a significant excess of D-lactylation at H3K18 and inversely an excess of L-lactylation at H3K23, both on canonical H3 and testis-specific H3 (H3.t) (**Suppl. Figures, section 4**). Besides, H3K9, H3K56 and H4K77 anchored balanced amounts of L- and D-lactylation. H3K31la could not be properly quantified, due to signal interferences coming from a highly abundant species (**Suppl. Figures, section 5**). Finally, intriguingly, the endogenous signal for H3K79la essentially matched the D-lactylated synthetic peptide, indicating that this specific residue was mostly D-lactylated (**Suppl. Figures, section 6**).

**Figure 5:**
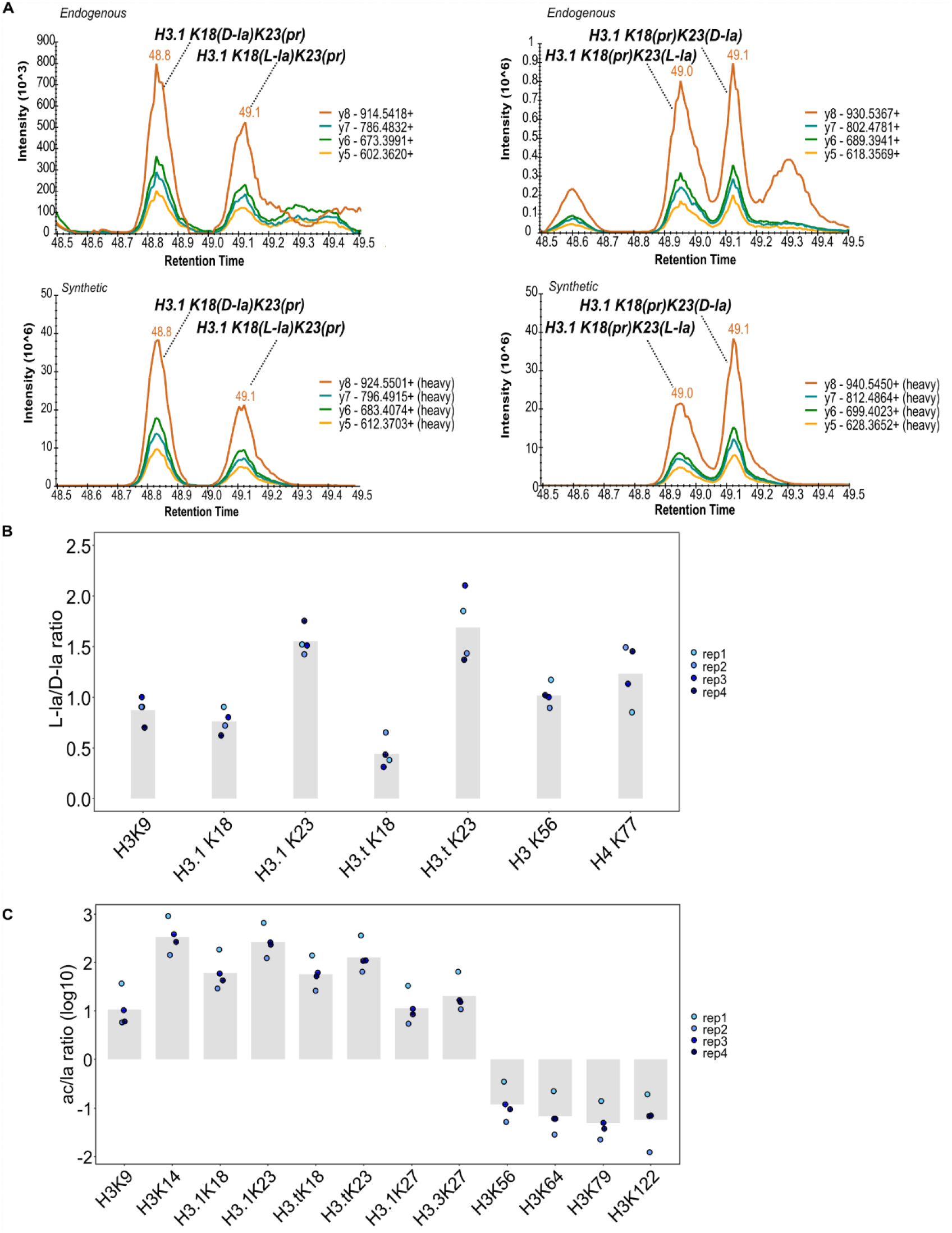
Validation of the co-existence of L- and D-lactylation on lysine residues of mouse testis histones H3 and H4 for which the two enantiomers are chromatographically separated, and relative abundance to one another and compared to acetylation. (A) Detection by selected y fragments of the presence of L- and D-lactylation on H3K18 (left panels) and H3K23 (right panels) in mouse testis histones. These fragments exhibit a perfect match between the endogenous (light) and heavy (synthetic) peptides. (B) Comparison of the abundances of L- and D-lactylation at the lysine sites for which the two enantiomers are chromatographically separated, in four biological replicates. (C) Abundance ratio of acetylation and lactylation at lysine residues of histone H3, in four biological replicates.

Next, we aimed to compare the relative abundance of lactylation to the canonical PTM acetylation at the same lysine sites. We then performed a targeted analysis of the four biological samples of mouse testis histones spiked with the L- and D-lactylated, as well as the corresponding acetylated synthetic peptides. Of note, we observed that the retention times of lactylated and acetylated peptides were very close, whatever the peptide sequence (**Suppl. Figures, section 7**). The relative abundance of variably modified peptides was determined by integrating the signals of a few y-type fragments, the same series being selected for acetylated and lactylated peptides (see **Suppl. Table 3**). Interestingly, acetylation appeared to be much more abundant at lysines of the N-terminal tail of both histones H3 and H4, whereas lactylation started becoming more abundant in the histone fold domain (**Figure 5c and Suppl. Figures, section 8**).

Finally, we sought to get an estimation of the stoichiometry of lactylation at all investigated lysine residues. Having formerly determined the relative abundance ac/la, we obtained the stoichiometry of acetylation. Exploratory analyses by DDA provided this information for all lysines of the N-terminal tails of histones H3 and H4, which are of sufficient abundance to be automatically selected for fragmentation (**Suppl. Table 4**). Besides, to estimate the stoichiometry of acetylation at lysines of the histone fold domains, we designed PRM analyses targeting the acetylated peptide and all variably modified peptide forms of same sequence, whose summed MS signals represented more than 90% of the MS signal for this sequence (**Suppl. Table 5**). The quantitative results of these analyses are provided in **Suppl. Table 6**. Finally, the estimated stoichiometry of acetylation and lactylation at all lysine sites are shown in **Table 1**. The well-known discrepancies in acetylation levels between H4K5/K8/K12 and H4K16, or between H3K9 and H3K14, were observed in our analyses of mouse testis histones. By contrast, lactylation levels at these compared sites were of very similar relative abundances, lying around 0.1%.

**Table 1:**
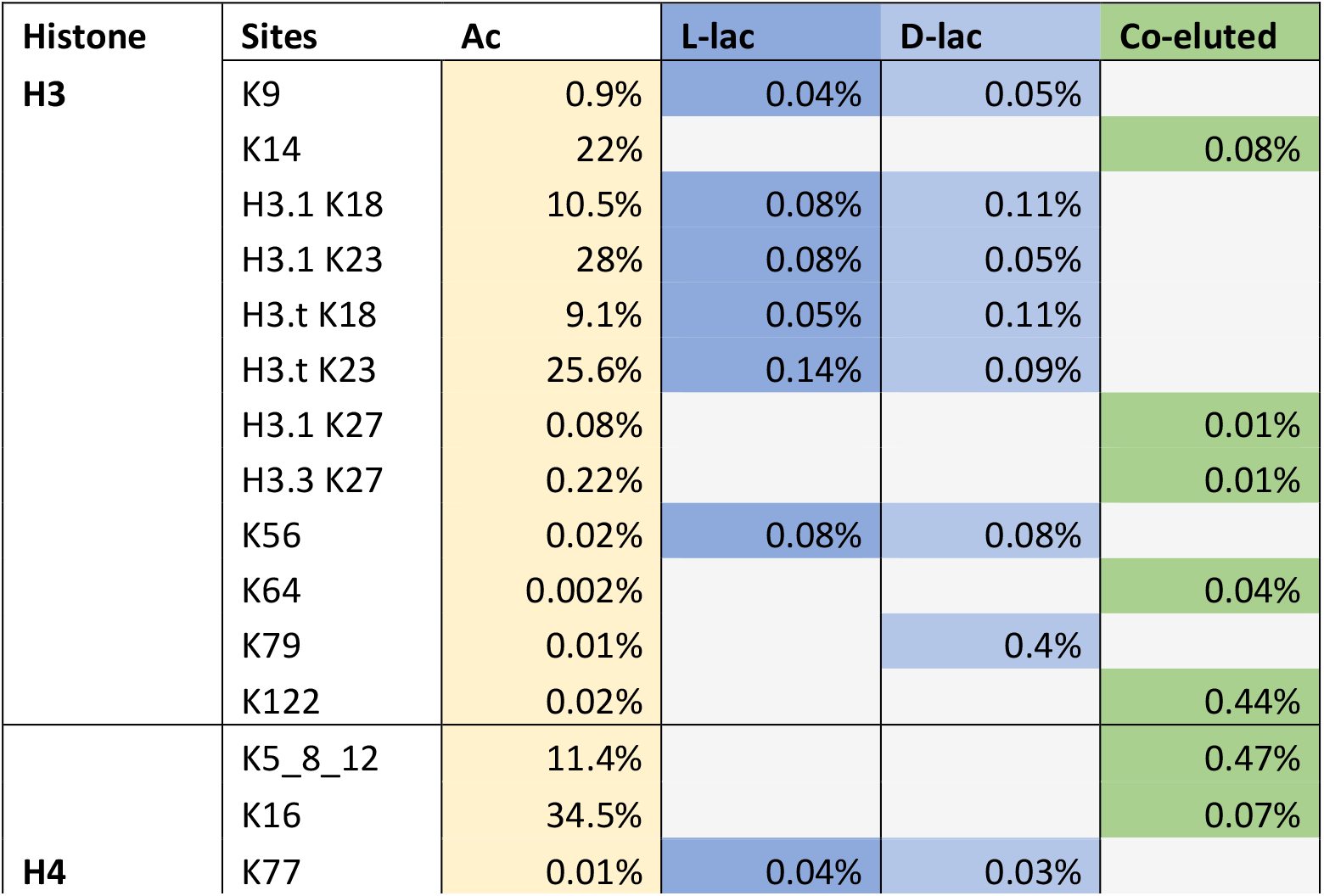
Estimated relative abundances of acetylation and lactylation (with the indication of the respective amounts of L- and D-lactylation when the enantiomers could be chromatographically separated) at all investigated lysines from histones H3 and H4. A summed abundance is indicated for residues H4K5, H4K8 and H4K12, whose positional isomers (both the acetylated and the lactylated ones) are insufficiently separated by LC to get their individual quantification.

| Histone | Sites | Ac | L-lac | D-lac | Co-eluted |
| --- | --- | --- | --- | --- | --- |
| **H3** | K9 | 0.9% | 0.04% | 0.05% |  |
|  | K14 | 22% |  |  | 0.08% |
|  | H3.1 K18 | 10.5% | 0.08% | 0.11% |  |
|  | H3.1 K23 | 28% | 0.08% | 0.05% |  |
|  | H3.t K18 | 9.1% | 0.05% | 0.11% |  |
|  | H3.t K23 | 25.6% | 0.14% | 0.09% |  |
|  | H3.1 K27 | 0.08% |  |  | 0.01% |
|  | H3.3 K27 | 0.22% |  |  | 0.01% |
|  | K56 | 0.02% | 0.08% | 0.08% |  |
|  | K64 | 0.002% |  |  | 0.04% |
|  | K79 | 0.01% |  | 0.4% |  |
|  | K122 | 0.02% |  |  | 0.44% |
| **H4** | K5_8_12 | 11.4% |  |  | 0.47% |
|  | K16 | 34.5% |  |  | 0.07% |
|  | K77 | 0.01% | 0.04% | 0.03% |  |

## Discussion

Our study pioneers in characterizing by proteomics the presence of L- and D-lactylation in a normal mouse tissue, when the majority of reports describing histone lactylation worked on cultured cells or tumors and most often relied on antibody-based experiments. We determined that lactylation occurred on a large number of lysine residues from histones H3 and H4 at a quite stable stoichiometry of 0.05-0.4% in mouse testis. This observation is in stark contrast with relative abundances of acetylation, which are highly variable between lysine sites, including the neighboring H3K9 versus H3K14 and H4K5/K8/K12 versus H4K16, being of low (1 to 5%) or high (20 to 35%) stoichiometry, respectively. Besides, while acetylation dominates on the N-terminal tails of H3 and H4 which are well accessible to HATs, lactylation becomes more abundant than acetylation in the histone fold domains. Our work further shows for the first time that some histone peptides modified by either L- or D-lactyl can be separated on a C18 column, when performing *in vitro* propionylation of free lysines before trypsin digestion. We could thus distinguish modification by the two enantiomers on H3K9, H3K18 and H3K23 on both canonical H3 and testis-specific H3, as well as H3K79 and H4K77. Some of these residues bore a balanced amount of D- and L-lactyl, with the exception of H3K18 and H3K79 which exhibited more or almost exclusive D-lactylation, whereas H3K23 appeared more modified by L-lactylation.

To reliably identify and quantify lactylation in histones from mouse testis, we considered synthetic peptides singly modified with this acylation at a lysine residue. However, most of these proteolytic peptides contain two or more lysines, and the lactylation site may co-exist with other modifications *in vivo*. For example, we detected H3K18lac with H3K23ac in our testis histone samples. Besides, H3K9lac was identified by proteomics in association with H3K14ac in HeLa cells^7^. Of note, both H3K14 and H3K23 are acetylated at high stoichiometries in most cell types, so that the combination of these acetylation sites with a lactylation on the neighboring lysine may be quite abundant and thus detectable by proteomic analysis. A similar reasoning may apply to the stretch H3K27-R40, in which acetylation of H3K27 is more abundantly detected in association with H3K36me2 in various tissues (e.g. mouse testis and brain), especially on variant H3.3. It might thus be worth exploring the hypothesis of the combination of PTMs H3 K27lac/K36me2, using doubly modified synthetic peptides.

L-lactylation has been most often considered to be the enantiomer that modifies protein lysines^5,6,27^, due to the much higher concentration of L-lactate in mammalian tissues and plasma compared to D-lactate^30^. This assumption is either explicitly expressed or is occulted in studies describing enzymes endowed with lactyltransferase activity. Besides, the non-enzymatic addition of D-lactylation has been proposed to occur on non-histone^2^ and histone proteins^11^. Our present study reinforces the importance to investigate whether one or both mechanisms may be at stake to modify mouse testis histones.

Indeed, D-lactate can be produced from MGO via the successive action of the glyoxalases GLO1 and GLO2^2^. Before being described as a source of protein lysine D-lactylation, MGO was shown to induce substantial modification of arginine residues, while the modification of lysines by carboxyethyl (Kce), an isomer of lactyl, was of minor abundance^31^. We indeed could not detect H3K18ce in our samples (**Suppl. Figures, section 9**). These chemical modifications occur over the whole histone sequences, impact the pattern of biological PTMs^31^ and perturb the assembly and stability of nucleosomes^32^. Accumulation of MGO glycation products was identified in cancer-related cells and not in normal cultured cells^32^. Zheng *et al*. reported PAD4 (protein arginine deiminase 4) to be a potent deglycase that converts arginine glycation, even due to long-term treatment with MGO, into citrulline, thus further antagonizing MGO-mediated chemical modification of this residue^33^. The enzyme DJ-1/PARK7 was also described as a histone deglycase which prevents and removes the chemical modifications at arginine^31,32^, yet with an action restricted to short-stage MGO adducts^33^. It is noteworthy that DJ-1 was originally characterized in 2012 as a glyoxalase that converts methylglyoxal to lactic acid^34^. To precisely assess whether DJ-1 bears deglycase activity and/or detoxifies MGO by glyoxalase activity, Gao *et al*. used mass spectrometry in addition to Western Blot to rule out the first enzymatic action^35^. These authors further demonstrated that DJ-1 is allosterically activated by glutathione (GSH) and that it produces a mixture of L- and D-lactate, whose relative amount, while always in favor of L-lactate, depends on the presence of GSH or of a glycated peptide. In parallel, Zhou *et al*. described the production of a racemic mixture of L- and D-lactate from MGO by DJ-1 in the absence of GSH^36^. In all, our observation of histone D-lactylation in mouse testes combined with these former publications would support the hypothesis of a production of a mixture of D- and L-lactate from MGO by GLO1 and/or DJ-1 in the chromatin vicinity. In particular, DJ-1 was formerly described as a chromatin-linked protein^37,38^. Furthermore, lysine-site-specific mechanisms may be at stake, as Trujillo *et al*. stressed the fact that deletion of GLO2 (and therefore accumulation of LGSH and lactyl-CoA) led to a significant increase solely at H3K18 and H3K79 lactylation^10^. It is interesting to note that we observed these two sites are enriched or almost specifically modified by D-lactylation, respectively, which may be in support of a mechanism of lactylation involving the MGO pathway.

Beyond the production of L- or D-lactate, the question whether lactylation occurs by spontaneous chemical reaction or is mediated by an enzyme, such as p300, AARS1/2, HBO1 and/or KAT2A remains open. Yet, given that we showed lactylation to be present at a stable stoichiometry over all lysine sites of histones H3 and H4, in stark contrast to acetylation events, a chemical addition may be a valid hypothesis to explore. In all cases, the tentative regulatory functions of lactylation on gene expression, both at lysines located in the N-terminal tail and in the histone fold domain should be explored. When pursuing this quest, the herein determined low stoichiometries of lactylation should prompt researchers to systematically acquire in parallel the whole genomic distributions of the lactylated and acetylated counterparts (e.g. H3K18lac and H3K18ac) by Chromatin immunoprecipitation followed by high-throughput sequencing (ChIP-seq) or cleavage under targets and tagmentation (CUT&Tag) technologies. This would allow verifying the occurrence of specific signals for lactylation, which would not be due to cross-reactivity of the anti-Kla antibody with the more abundant acetylated site. This was carefully performed in the original description of histone lactylation focusing on H3K18^1^ but was not systematically carried out in later studies.

Based on the herein presented results, future work should enquire (i) whether the modification by L- and D-lactyl is also observable on other lysines of histones H3 and H4 from mouse testis, and beyond, on histones H2A, H2B and H1, possibly by implementing a chiral reaction to increase the chromatographic separation of variably modified peptides^27,39^ or by ion mobility^40^; (ii) whether the two enantiomers modify histone lysines in other mouse organs, but also in organs of other mammals including humans; (iii) what enantiomers of lactyl modify non-histone proteins in the former tissues; (iv) how this landscape of lactylation evolves during spermatogenesis at stages where glycolysis is particularly active, and lactate becomes the predominant source of energy (i.e. in meiotic and postmeiotic cells). At these stages, does L-lactylation dominate over D-lactylation? Of note, the latter point was precisely studied in MCF7 cells, i.e. breast cancer cells, where glycolysis is expected to fuel cells with L-lactate, and the authors concluded that the major enantiomer modifying histone lysine was indeed L-lactyl ^27^.

## Acknowledgements

We thank our colleagues at EDyP for excellent technical support in proteomic analyses and informatics handling of data. The proteomic experiments were supported by Agence Nationale de la Recherche under projects ProFI (Proteomics French Infrastructure, ANR-10-INBS-08) and GRAL, a program from the Chemistry Biology Health (CBH) Graduate School of University Grenoble Alpes (ANR-17-EURE-0003). This project was largely funded by ANR-21-CE44-0035-01 to DP and JC. PhD funding for JM was also from ANR (ANR-21-CE44-0035-01) and to HH was from GRAL.

