## Supplementary figures for "Both L-lactyl and D-lactyl enantiomers modify histones in mouse testis"

**Section 1**: Identification by proteomics of potentially lactylated sites on histones H3 and H4 from mouse testis

### H3 K9STGGK14APR

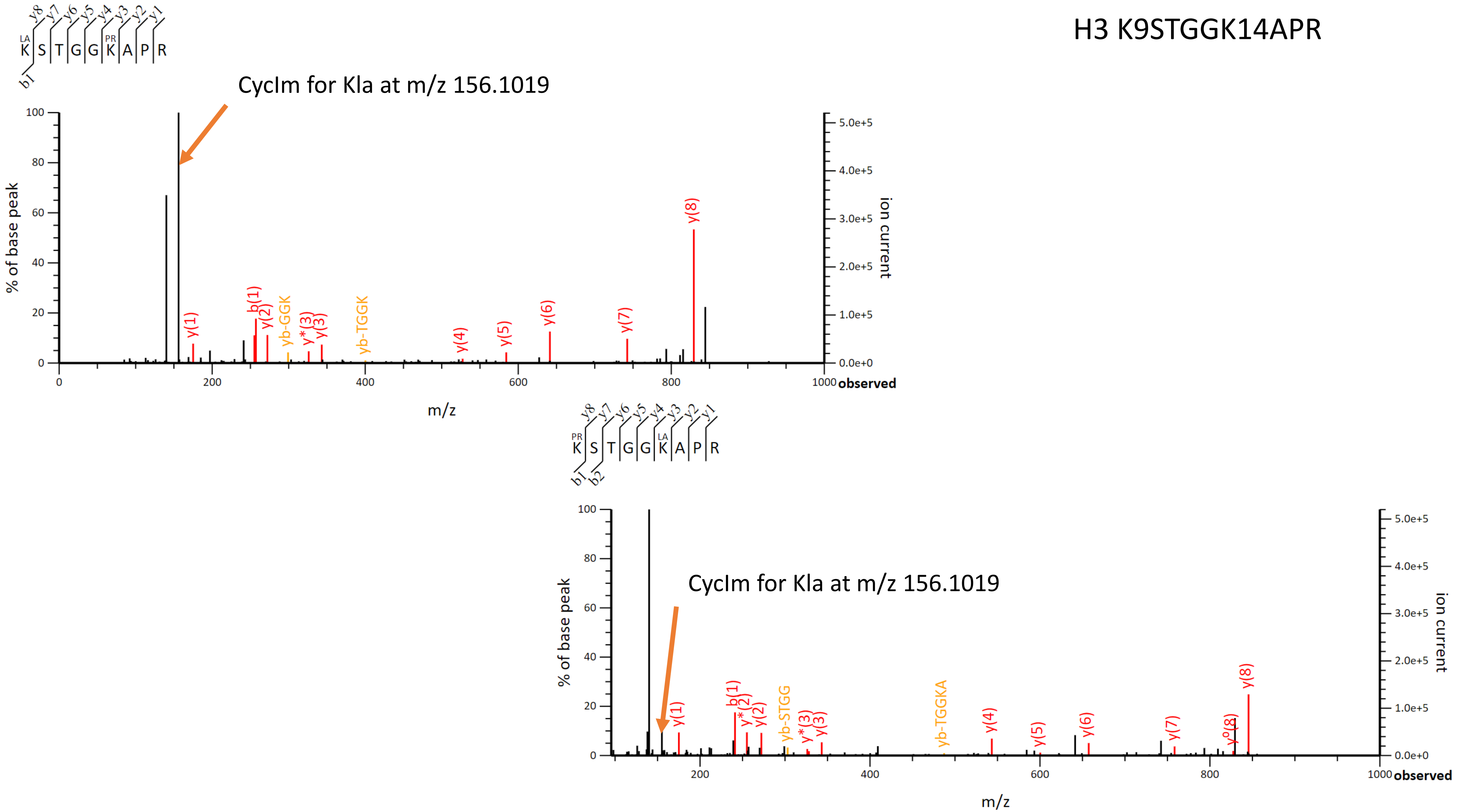

#### H3.1 K18QLATK23AAR

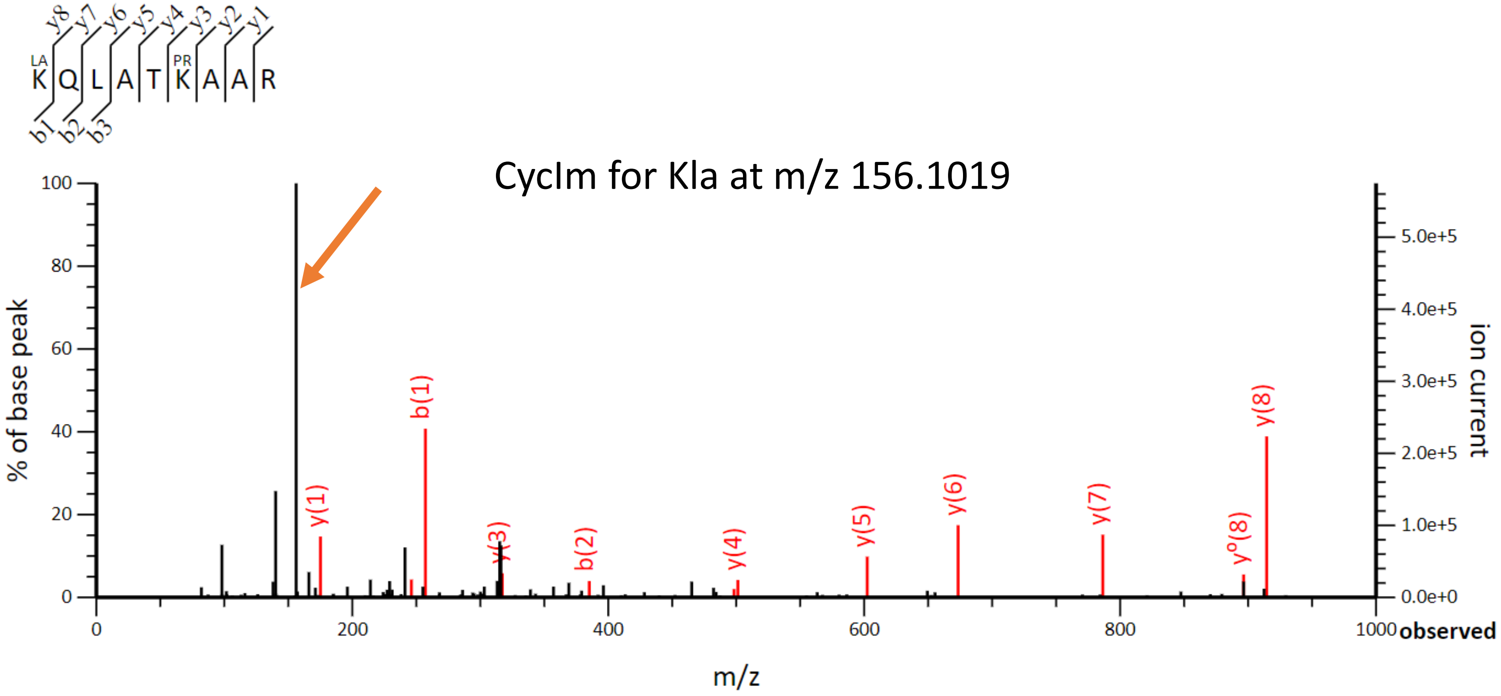


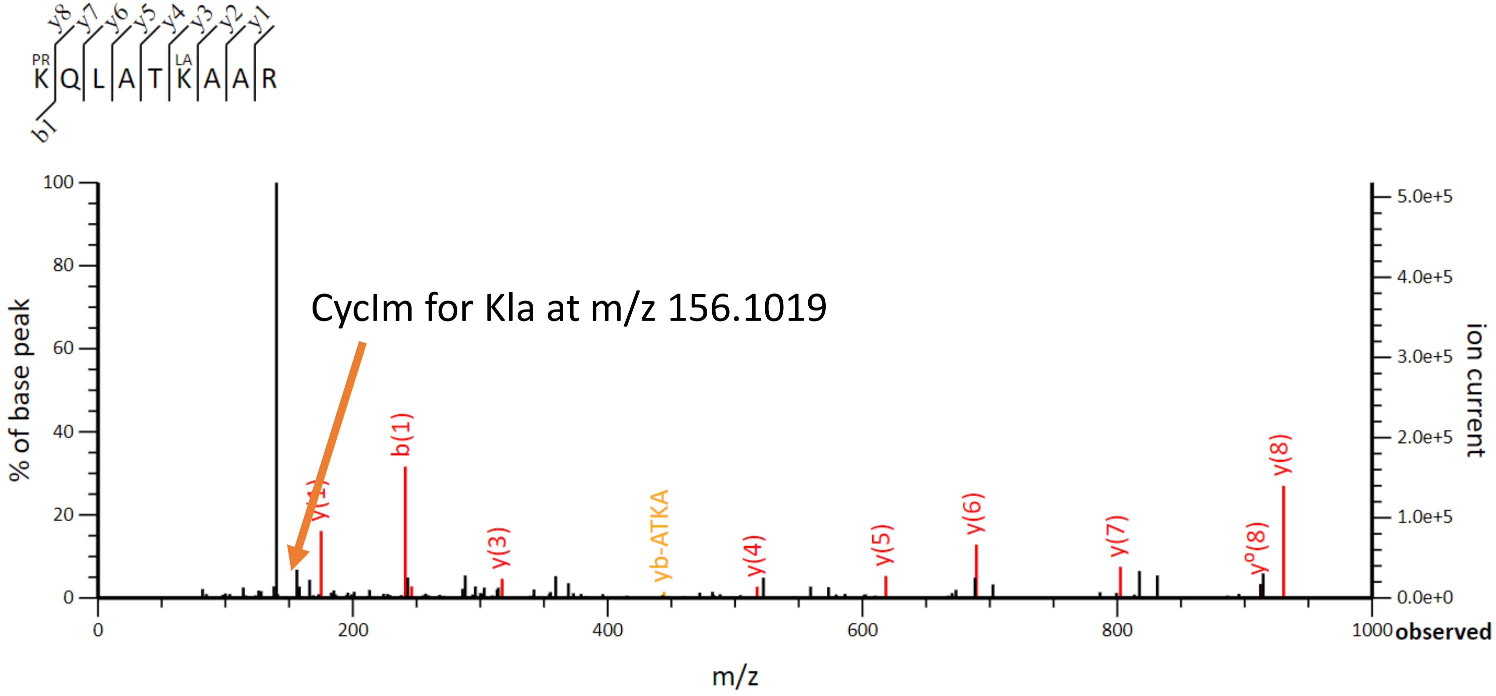

### H3.T K18QLATK23VAR

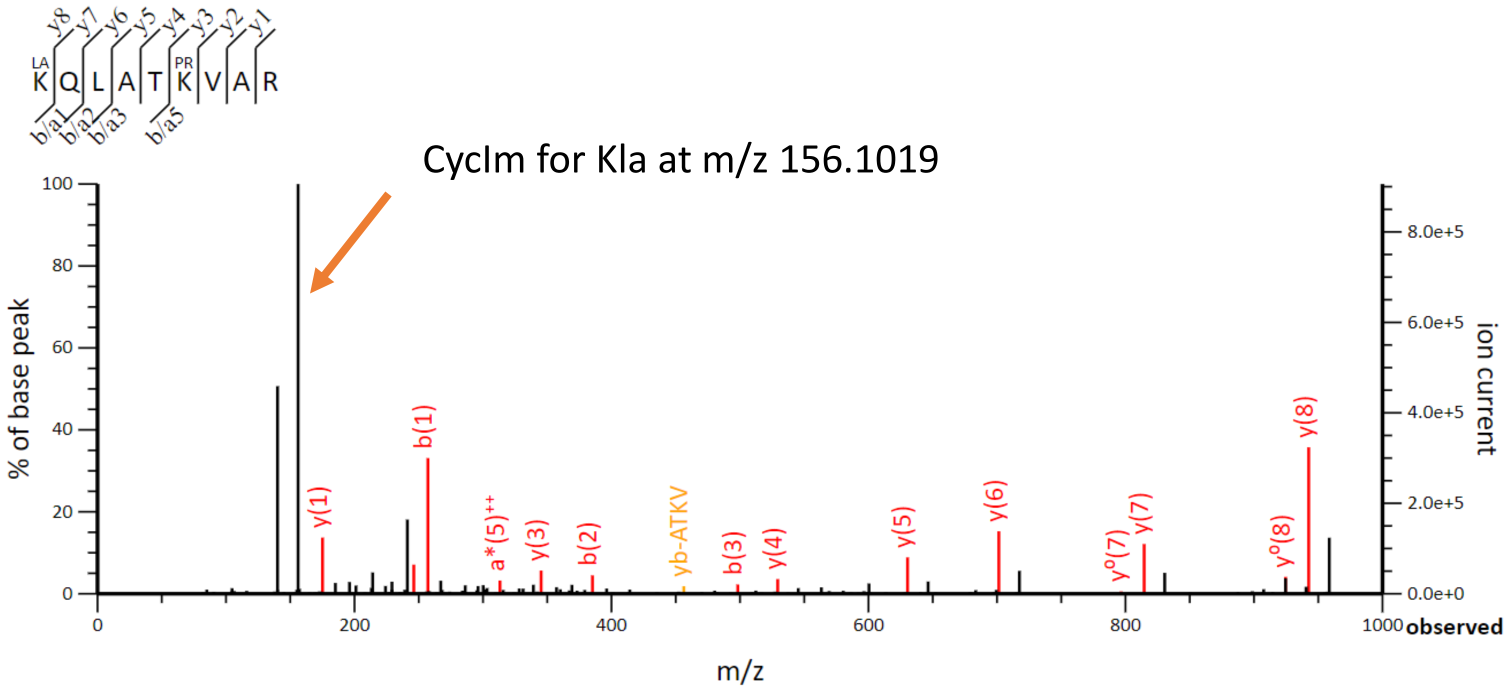


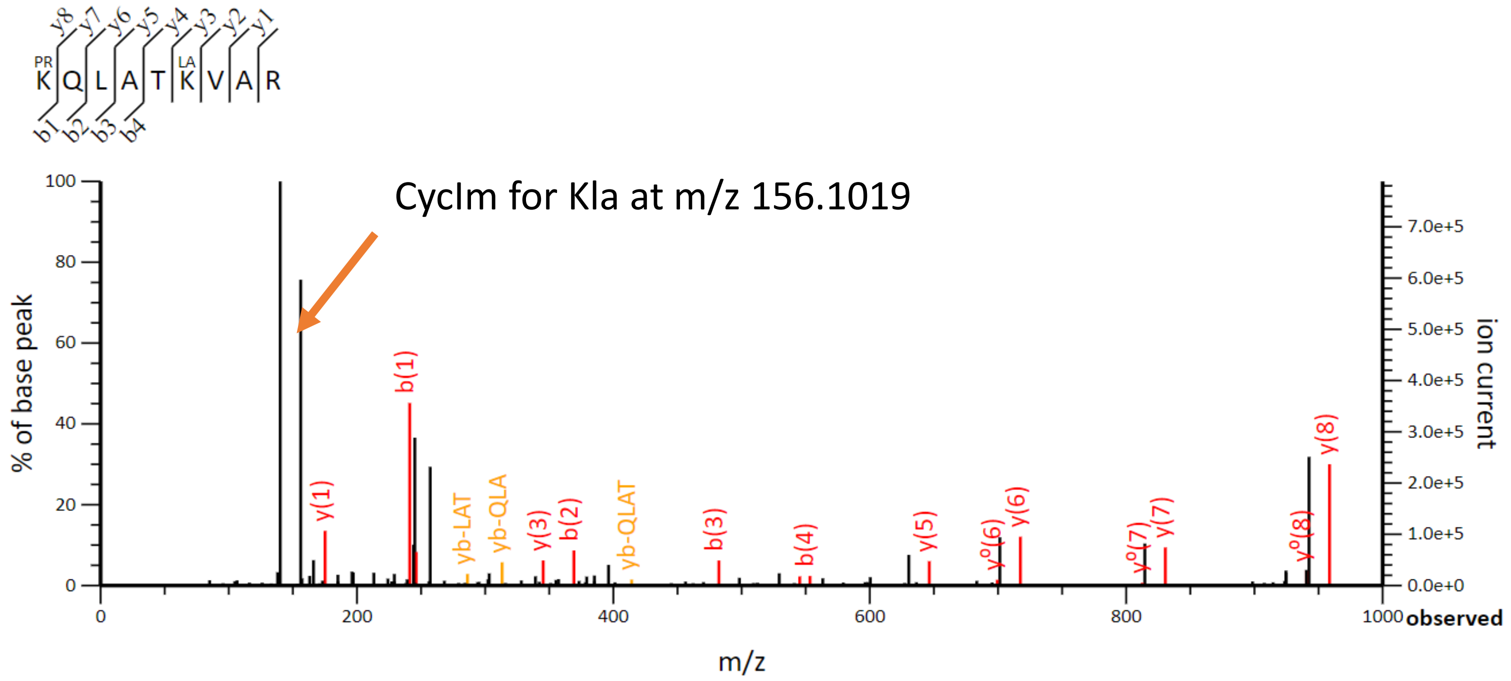

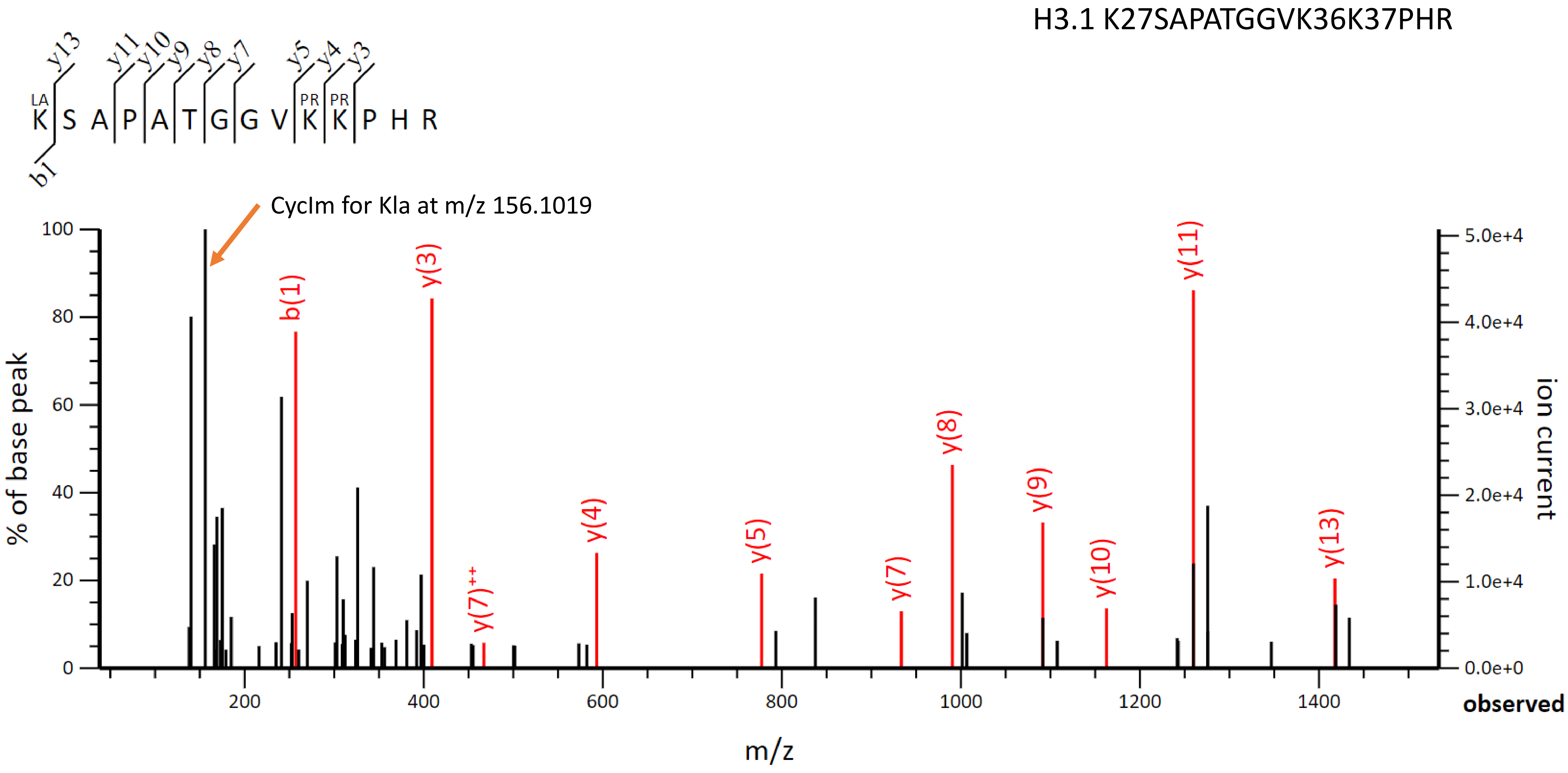

H3.1 K27SAPATGGVK36K37PHR
K S A P A T G G V K K P H R
LA
PR
PR
b1
y13
y11
y10
y9
y8
y7
y5
y4
y3
CycIm for Kla at m/z 156.1019
b(1)
y(3)
y(7)++
y(4)
y(5)
y(7)
y(8)
y(9)
y(10)
y(11)
y(13)
% of base peak
ion current
m/z
observed

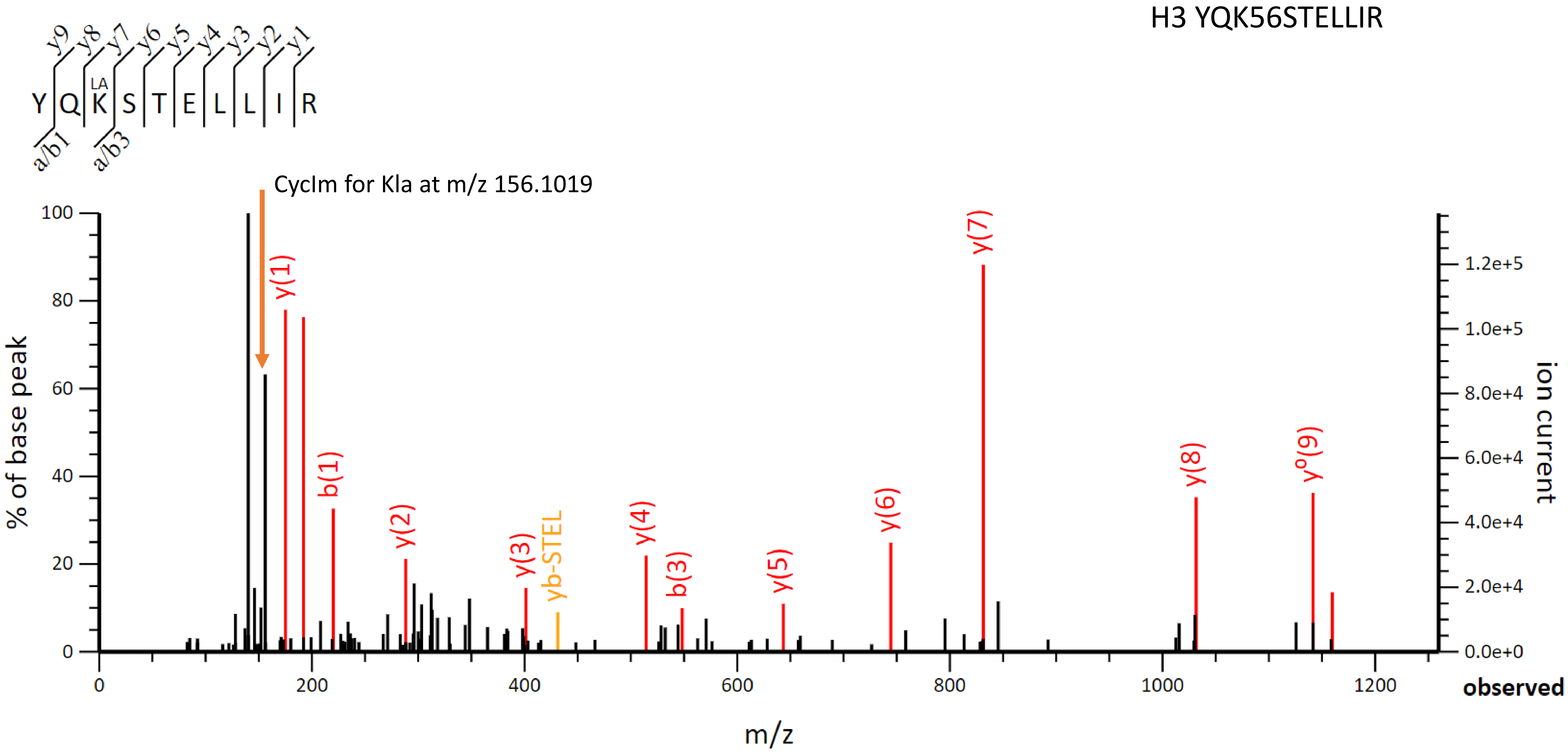

H3 YQK56STELLIR
y9 y8 y7 y6 y5 y4 y3 y2 y1
Y Q LA K S T E L L I R
a/b1 a/b3
CycIm for Kla at m/z 156.1019
% of base peak
ion current
m/z
observed
y(1)
b(1)
y(2)
y(3)
yb-STEL
y(4)
b(3)
y(5)
y(6)
y(7)
y(8)
y°(9)

H3 K64LPFQR

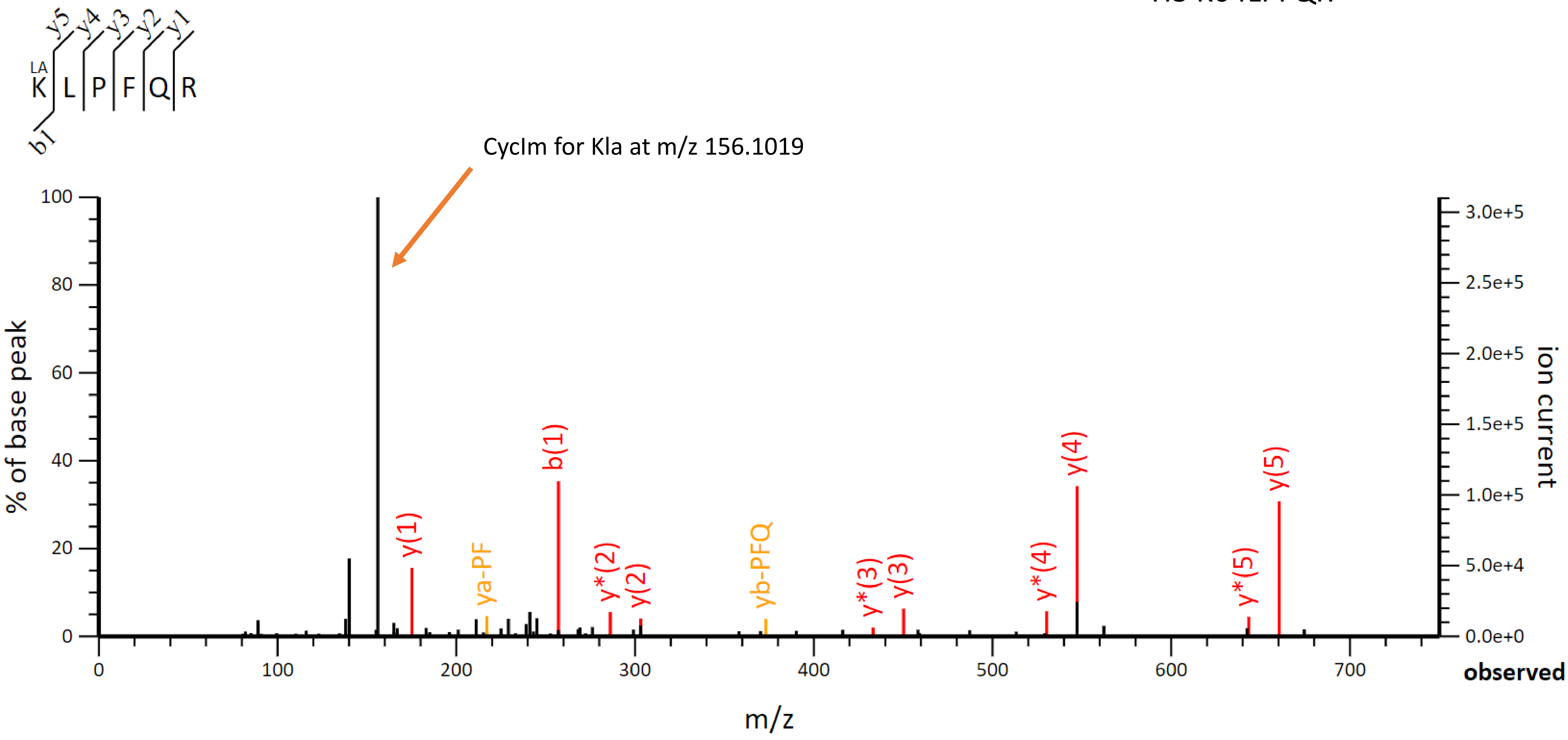

y5 y4 y3 y2 y1
LA
K L P F Q R
b1
CycIm for Kla at m/z 156.1019
100
80
60
40
20
0
% of base peak
y(1)
ya-PF
b(1)
y*(2)
y(2)
yb-PFQ
y*(3)
y(3)
y*(4)
y(4)
y*(5)
y(5)
3.0e+5
2.5e+5
2.0e+5
1.5e+5
1.0e+5
5.0e+4
0.0e+0
ion current
0
100
200
300
400
500
600
700
observed
m/z

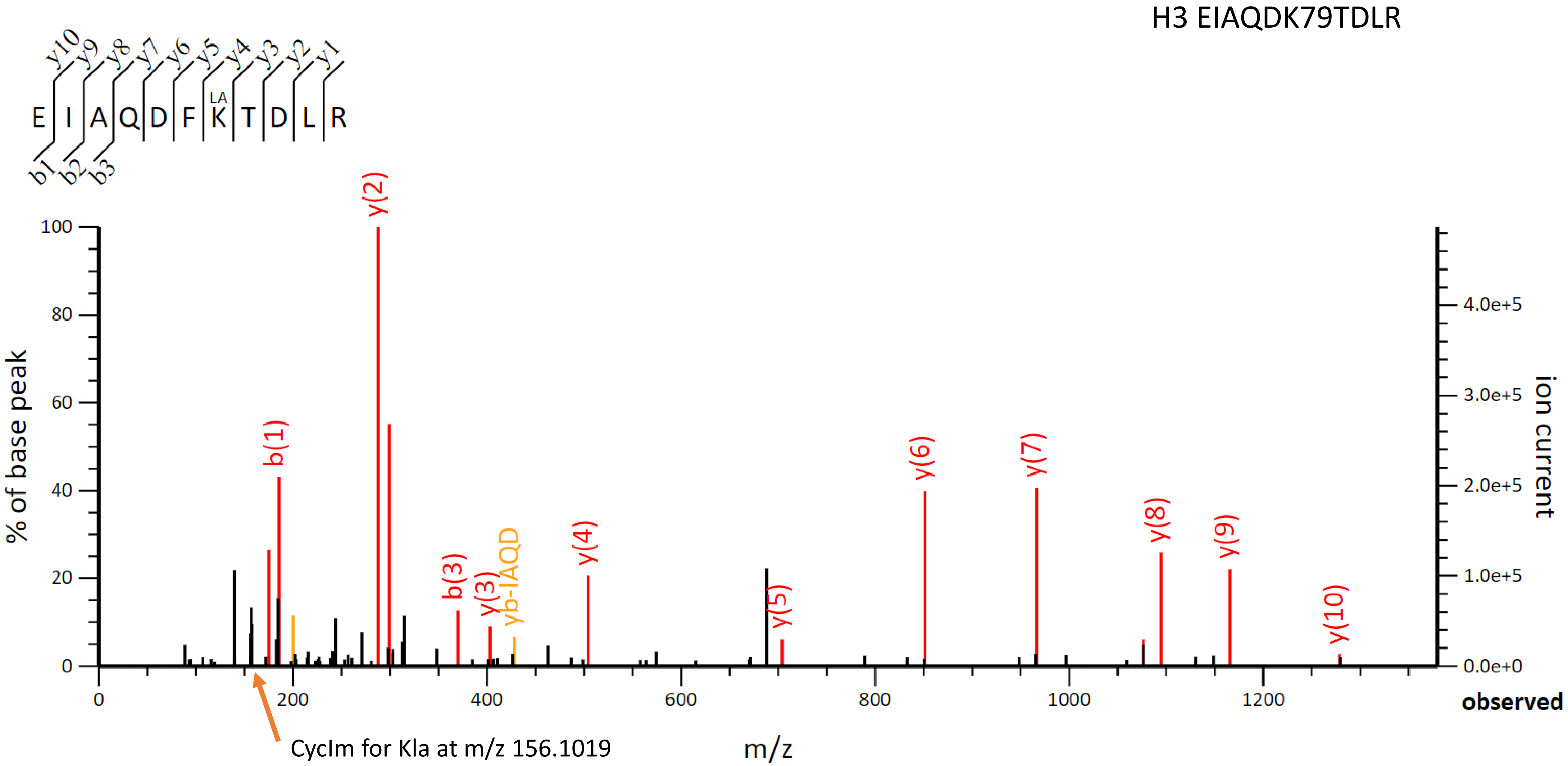
H3 EIAQDK79TDLR
E I A Q D F K T D L R
LA
y10 y9 y8 y7 y6 y5 y4 y3 y2 y1
b1 b2 b3
% of base peak
ion current
observed
m/z
y(2)
b(1)
b(3)
y(3)
yb-IAQD
y(4)
y(5)
y(6)
y(7)
y(8)
y(9)
y(10)
CycIm for Kla at m/z 156.1019

b1 ion for Valine is expected at mass 156.1019, that is the same m/z as for the CycIm of Klac.
We validated K122(lac) based on fragments y6 and y7.

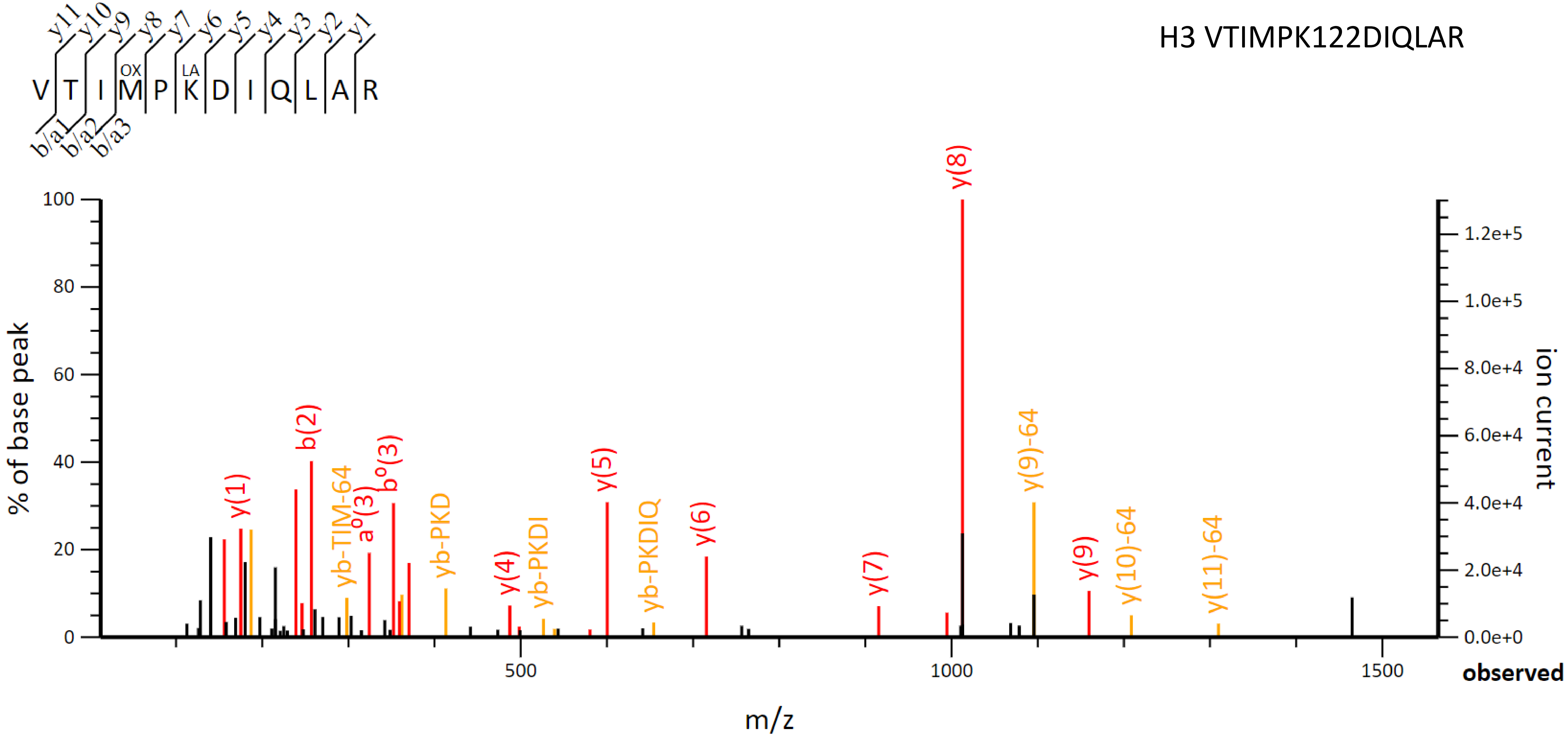

H4 GK5GGK8GLGK12GGAK16R

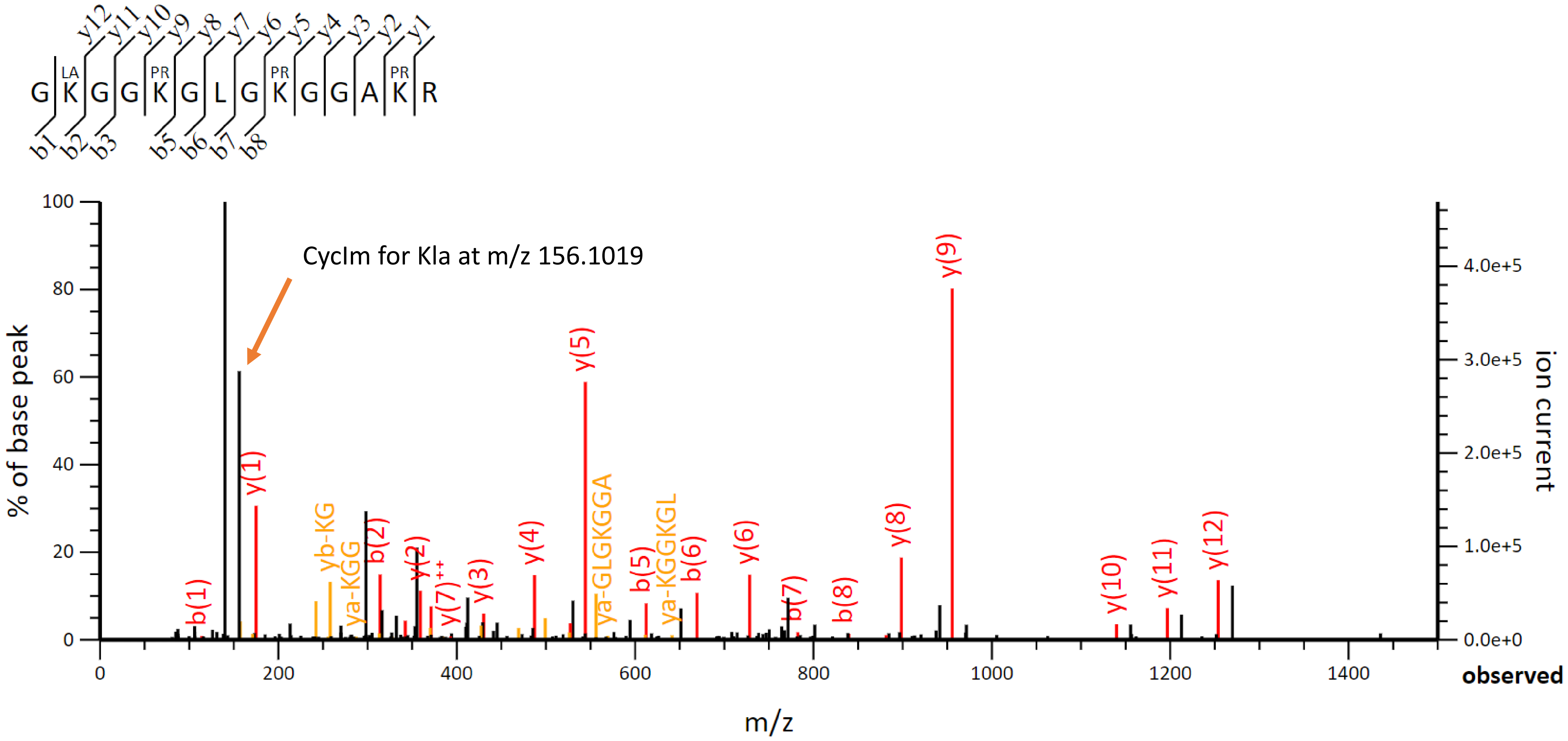

H4 GK5GGK8GLGK12GGAK16R

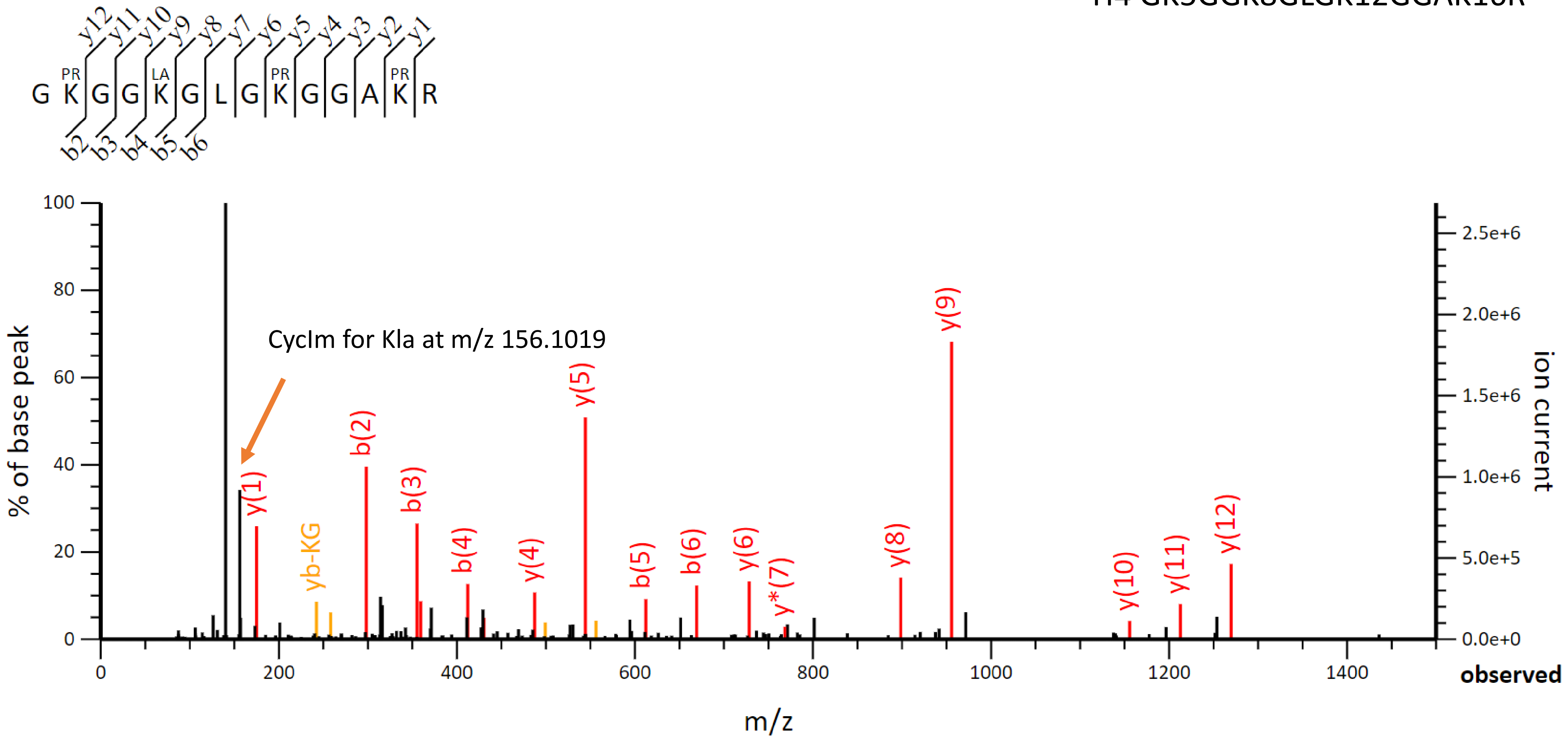

H4 GK5GGK8GLGK12GGAK16R

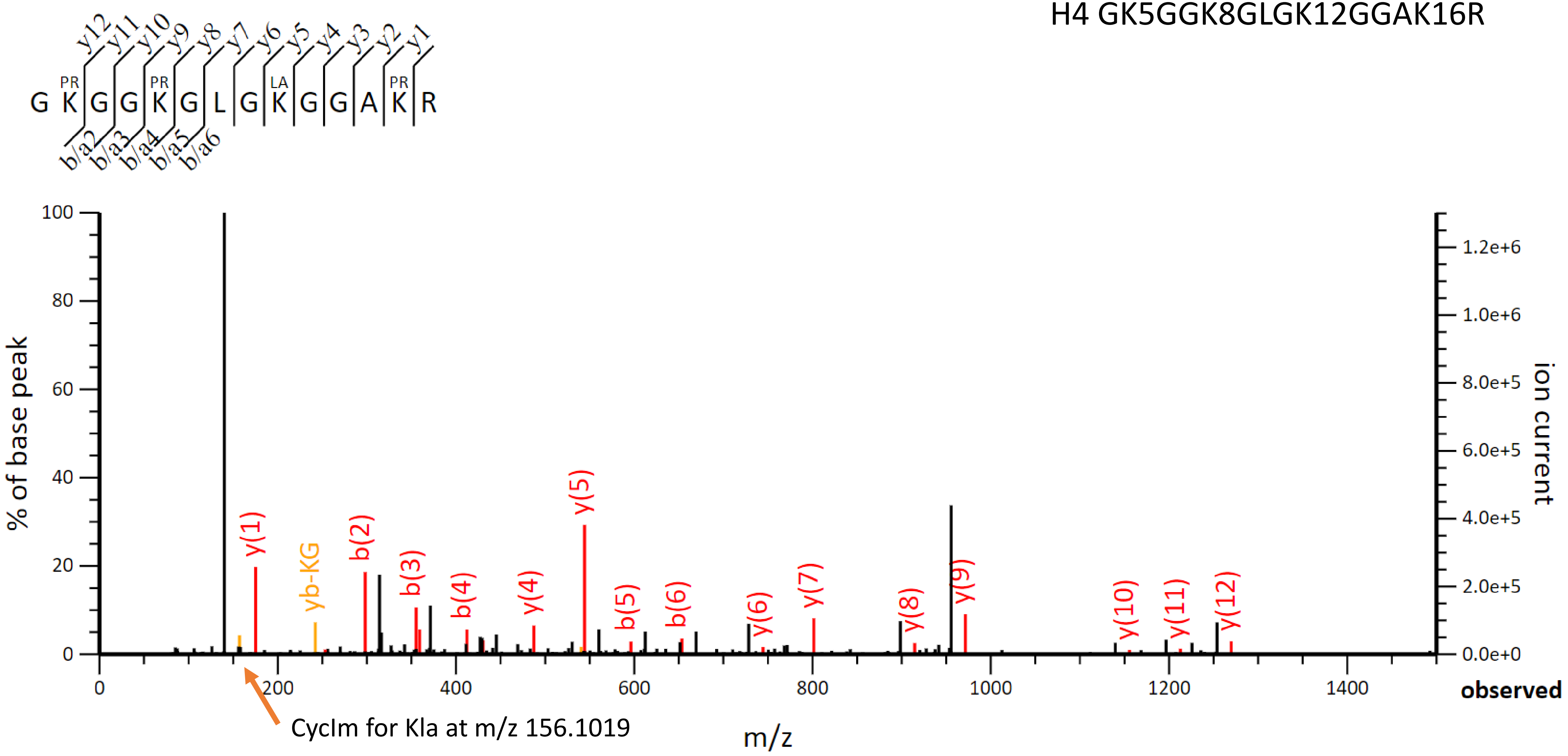


CycIm for Kla at m/z 156.1019

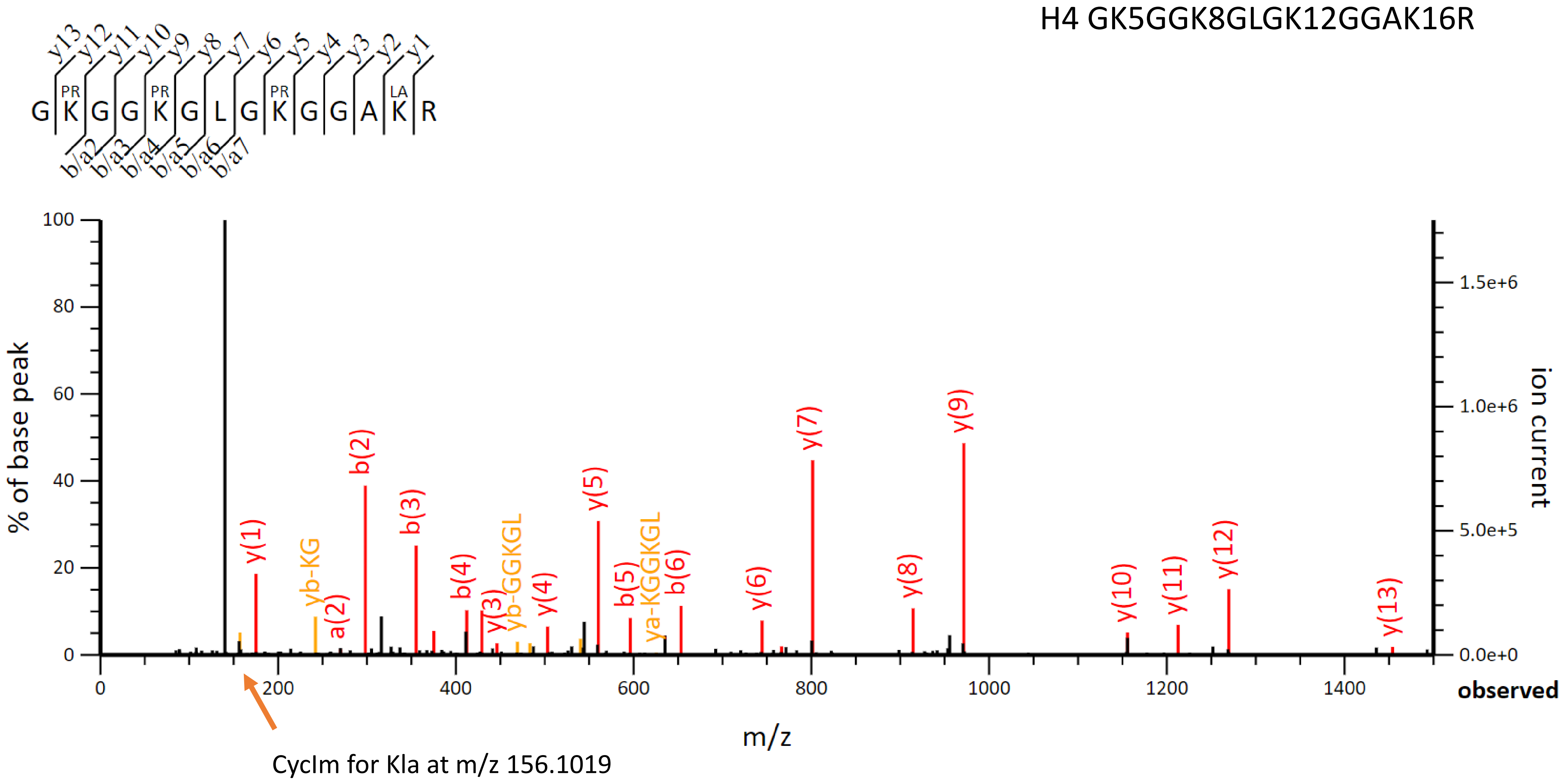
H4 GK5GGK8GLGK12GGAK16R
y13 y12 y11 y10 y9 y8 y7 y6 y5 y4 y3 y2 y1
G PR K G G PR K G L G PR K G G A LA K R
b/a2 b/a3 b/a4 b/a5 b/a6 b/a7
% of base peak
ion current
m/z
observed
y(1) yb-KG a(2) b(2) b(3) b(4) y(3) yb-GGKGL y(4) y(5) b(5) ya-KGGKGL b(6) y(6) y(7) y(8) y(9) y(10) y(11) y(12) y(13)
CycIm for Kla at m/z 156.1019

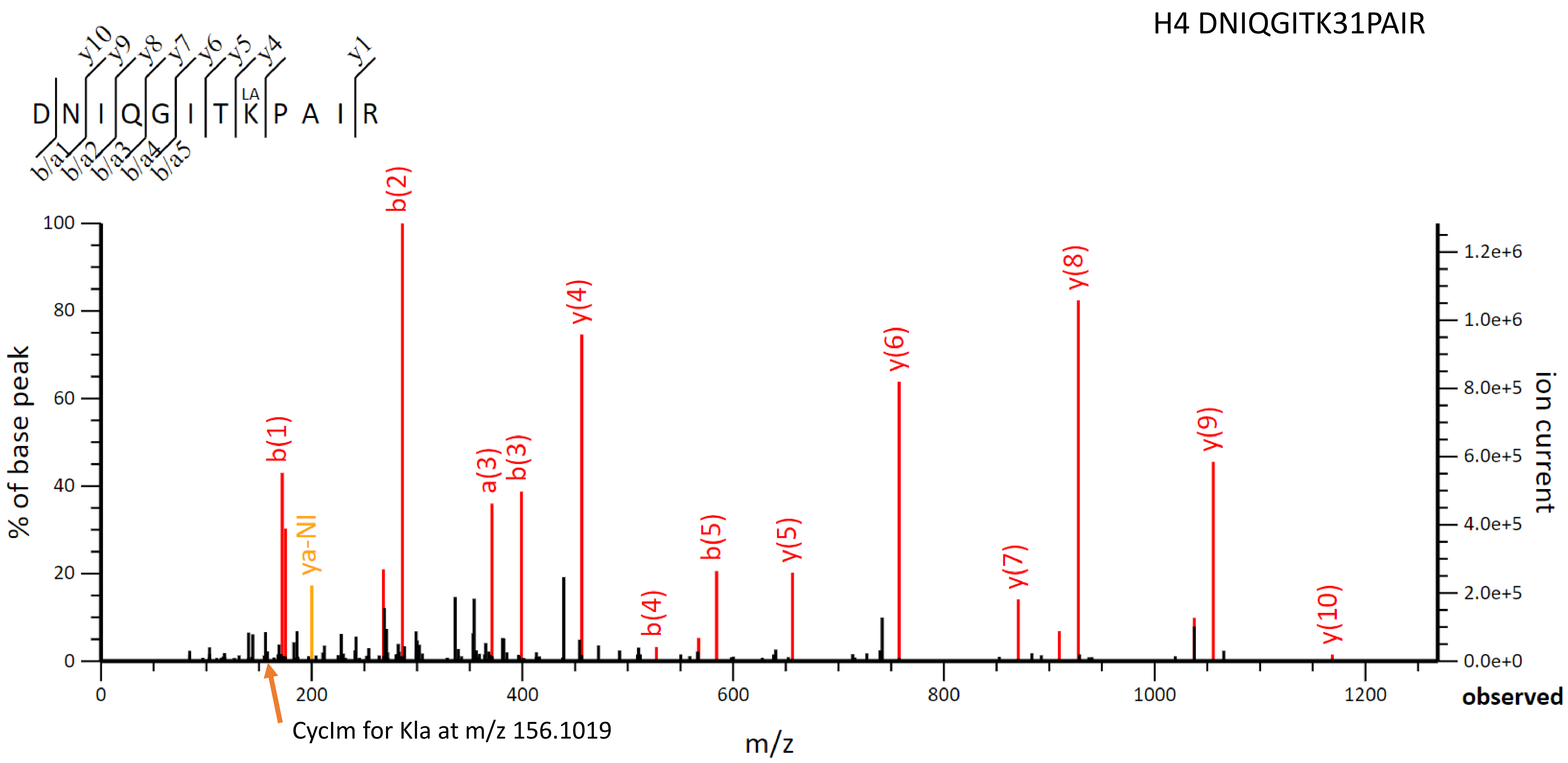
H4 DNIQGITK31PAIR
D N I Q G I T K P A I R
LA
y10 y9 y8 y7 y6 y5 y4 y1
b/a1 b/a2 b/a3 b/a4 b/a5
b(1)
ya-NI
b(2)
a(3)
b(3)
y(4)
b(4)
b(5)
y(5)
y(6)
y(7)
y(8)
y(9)
y(10)
% of base peak
ion current
m/z
observed
CycIm for Kla at m/z 156.1019

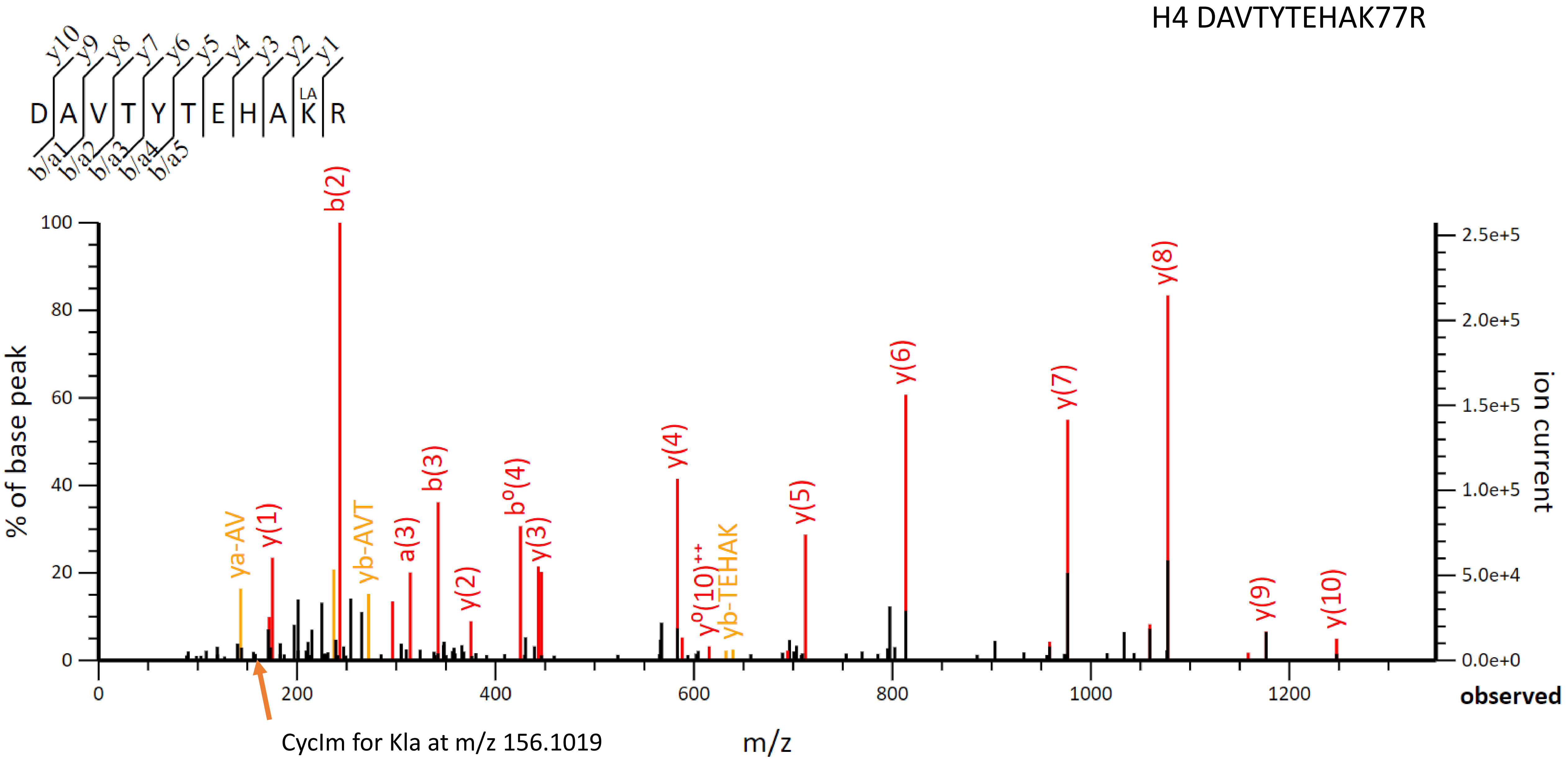
H4 DAVTYTEHAK77R
y10 y9 y8 y7 y6 y5 y4 y3 y2 y1
D A V T Y T E H A K R
LA
b/a1 b/a2 b/a3 b/a4 b/a5
% of base peak
ion current
b(2)
y(8)
y(6)
y(7)
y(4)
b(3)
b°(4)
y(5)
ya-AV
y(1)
yb-AVT
a(3)
y(3)
y(2)
y°(10)++
yb-TEHAK
y(9)
y(10)
observed
m/z
CycIm for Kla at m/z 156.1019

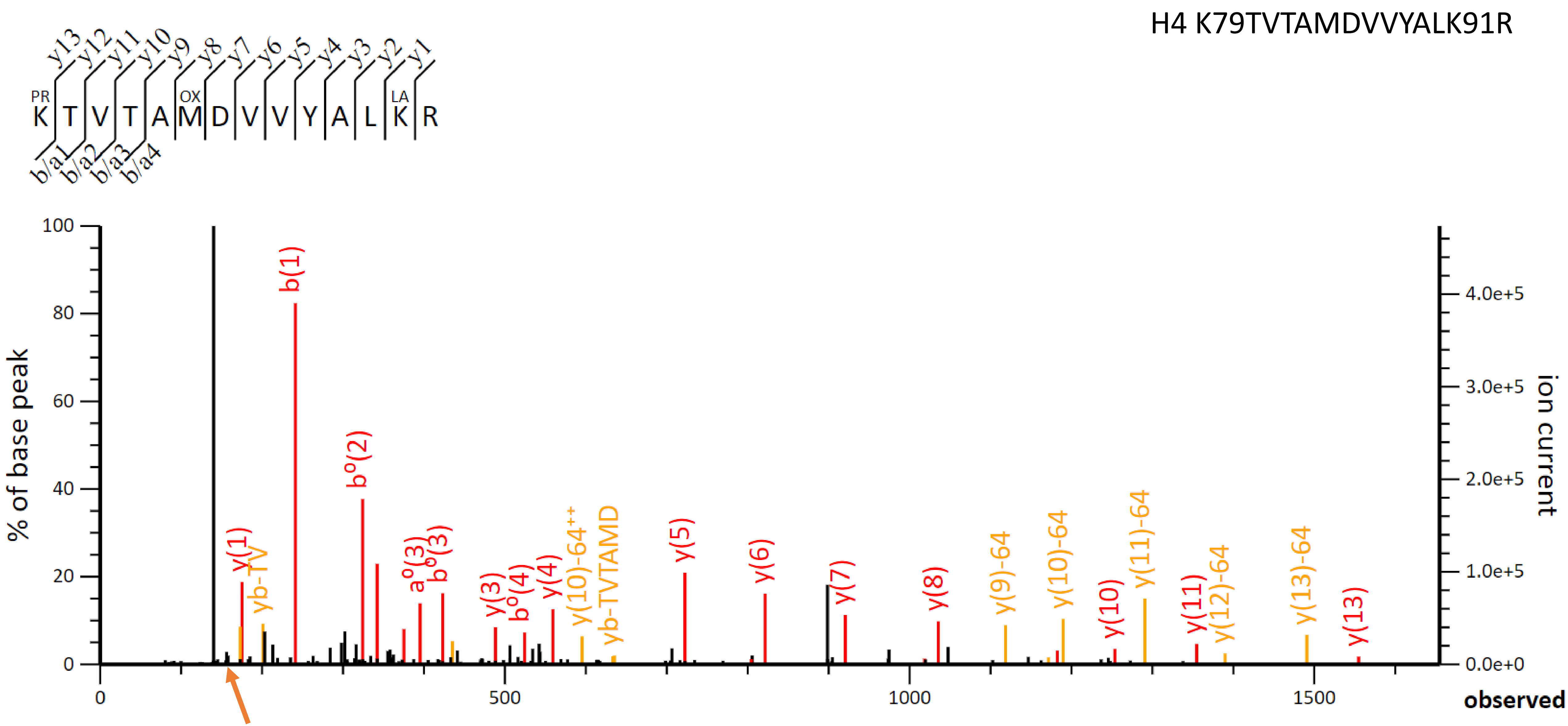


CycIm for Kla at m/z 156.1019

**Section 2:**

False identifications of presumably lactylated peptides correspond to propionylated and oxidized peptides, which can be of prevailing abundance

#### Oxidation events on histone H4 DNIQGITK$_{31}$PAIR peptide, m/z =727.4041,2+

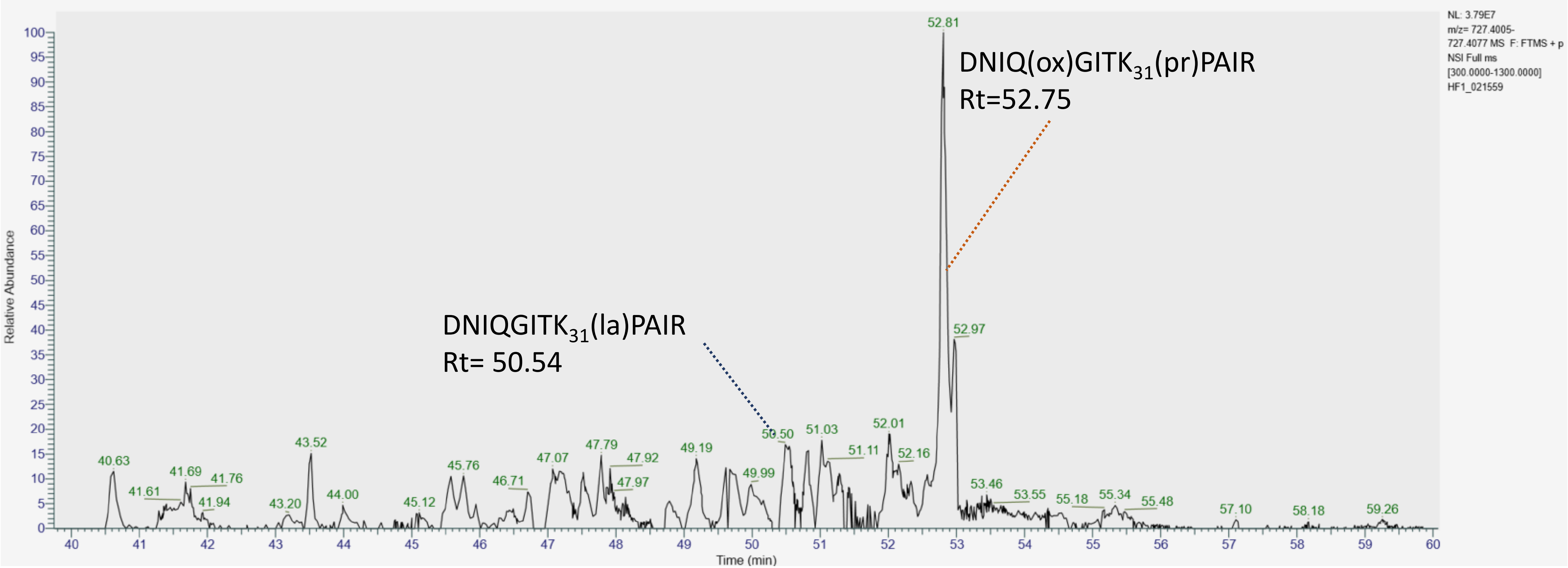

Scan 24196 , rt=52.751, HF1_021559.raw, 171924.dat

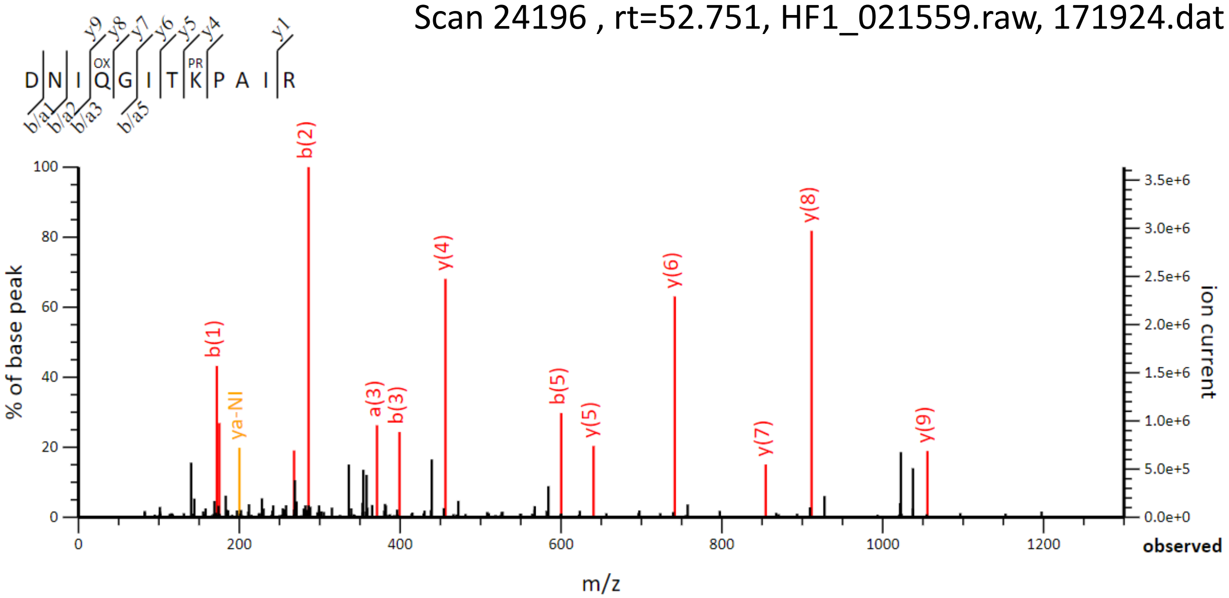


Scan 23089 , rt=50.537, HF1_021559.raw, 171924.dat

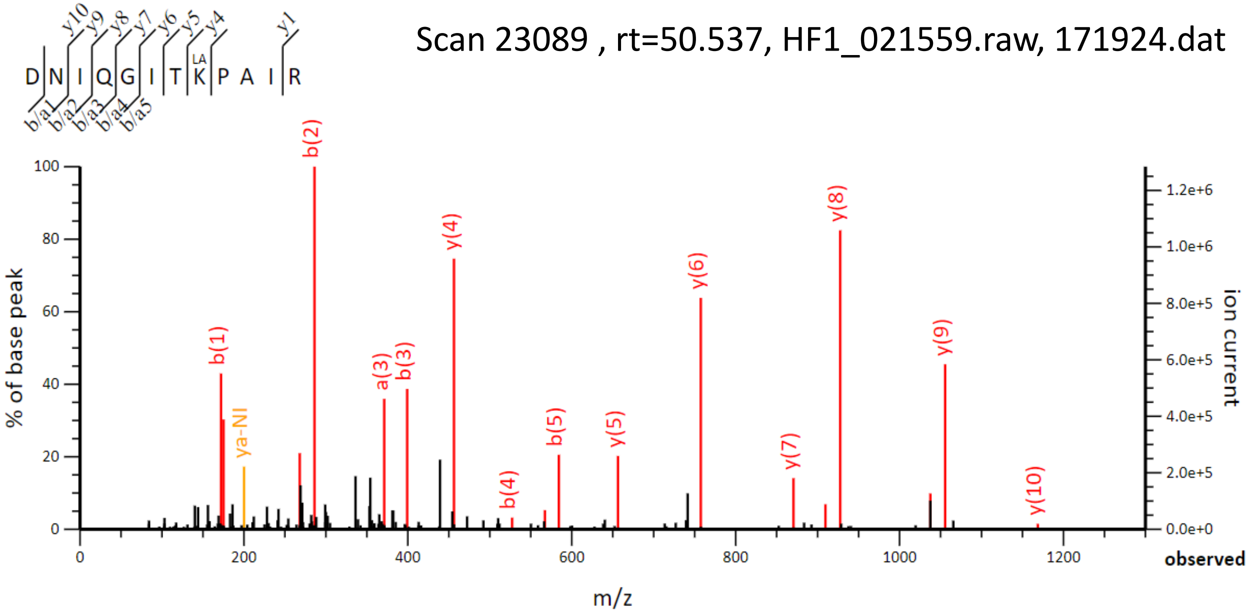

### Oxidation events on histone H4 DAVTYTEHK$_{77}$R peptide, m/z = 709.8491 (2+)

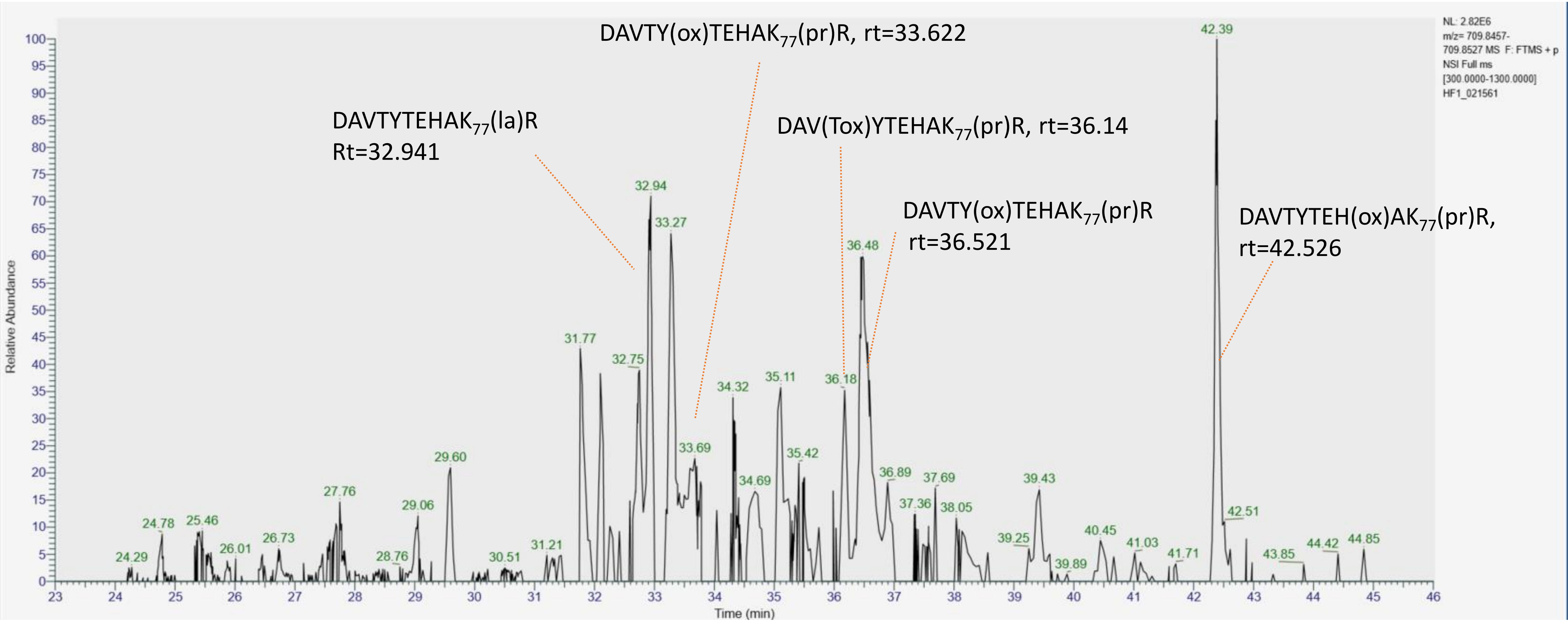

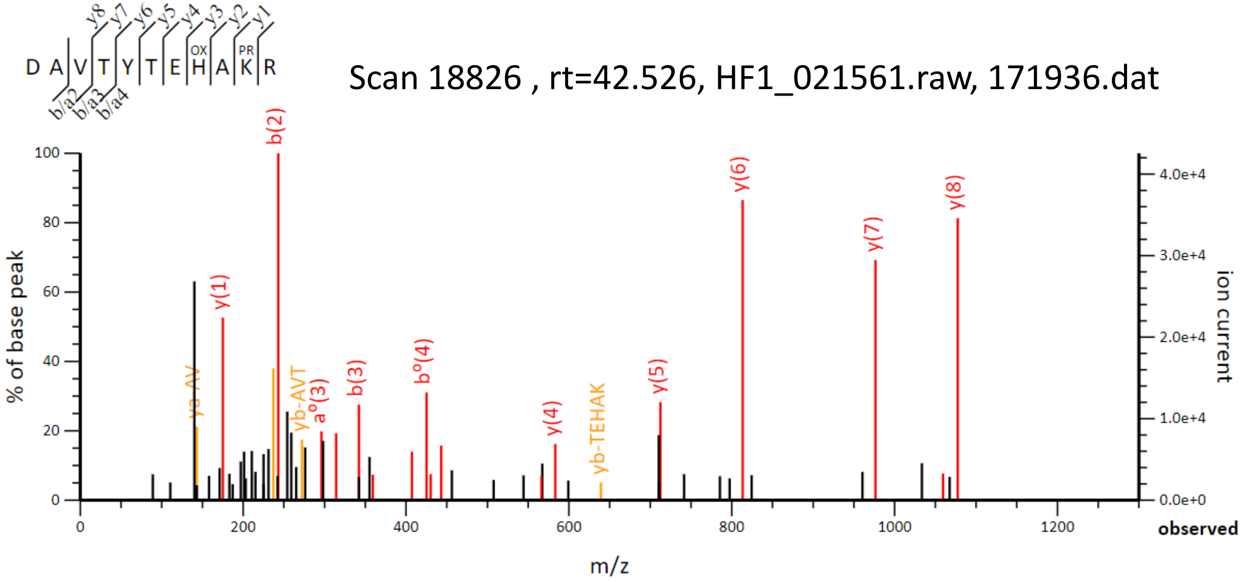

D A V T Y T E H A K R
Scan 18826 , rt=42.526, HF1_021561.raw, 171936.dat
% of base peak
ion current
m/z
observed


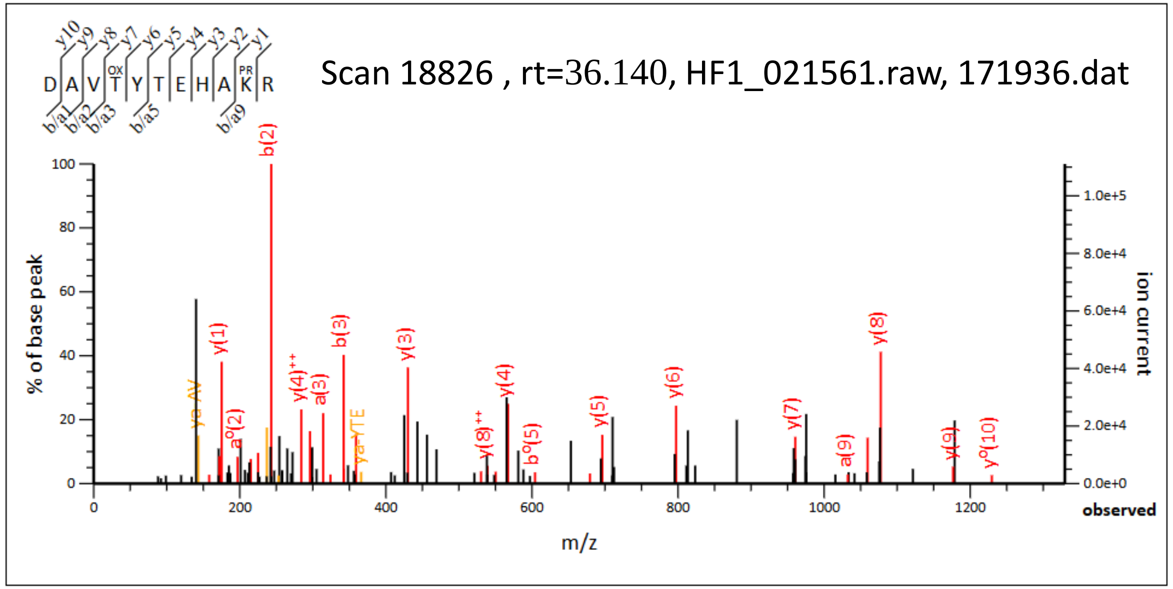

D A V T Y T E H A K R
Scan 18826 , rt=36.140, HF1_021561.raw, 171936.dat
% of base peak
ion current
m/z
observed


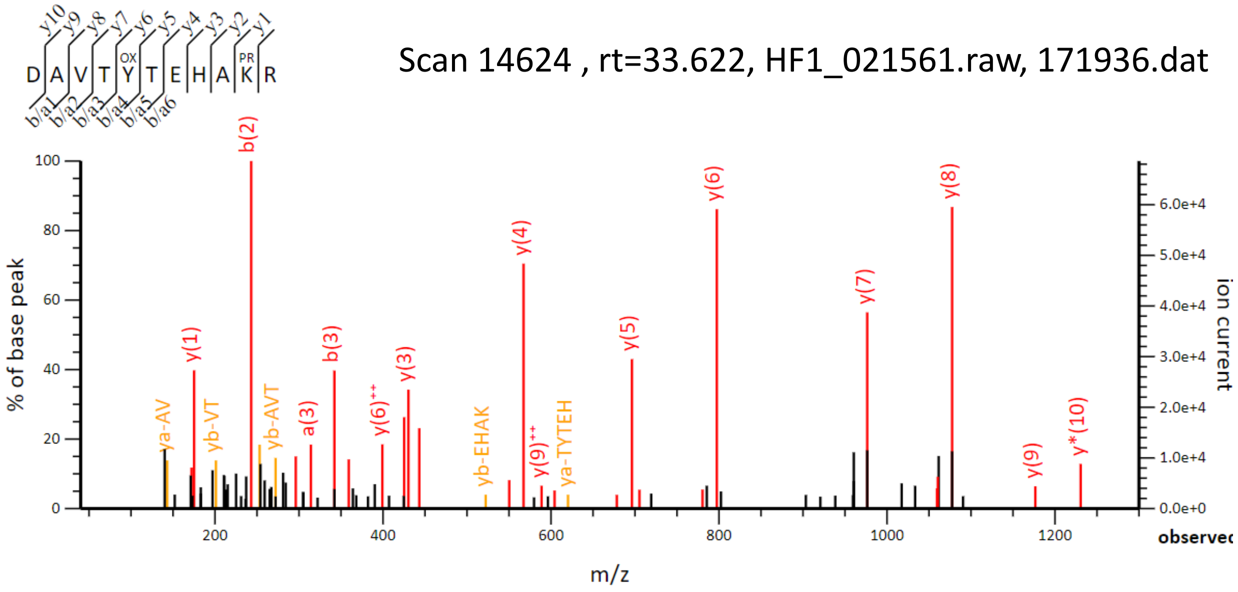

D A V T Y T E H A K R
Scan 14624 , rt=33.622, HF1_021561.raw, 171936.dat
% of base peak
ion current
m/z
observed


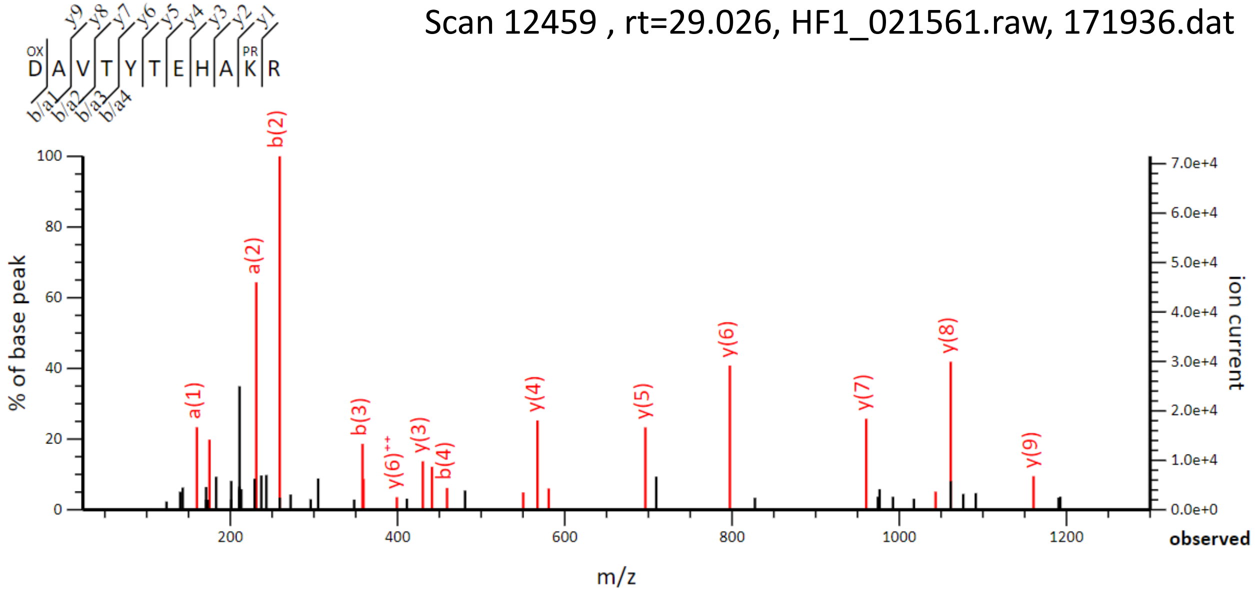

D A V T Y T E H A K R
Scan 12459 , rt=29.026, HF1_021561.raw, 171936.dat
% of base peak
ion current
m/z
observed

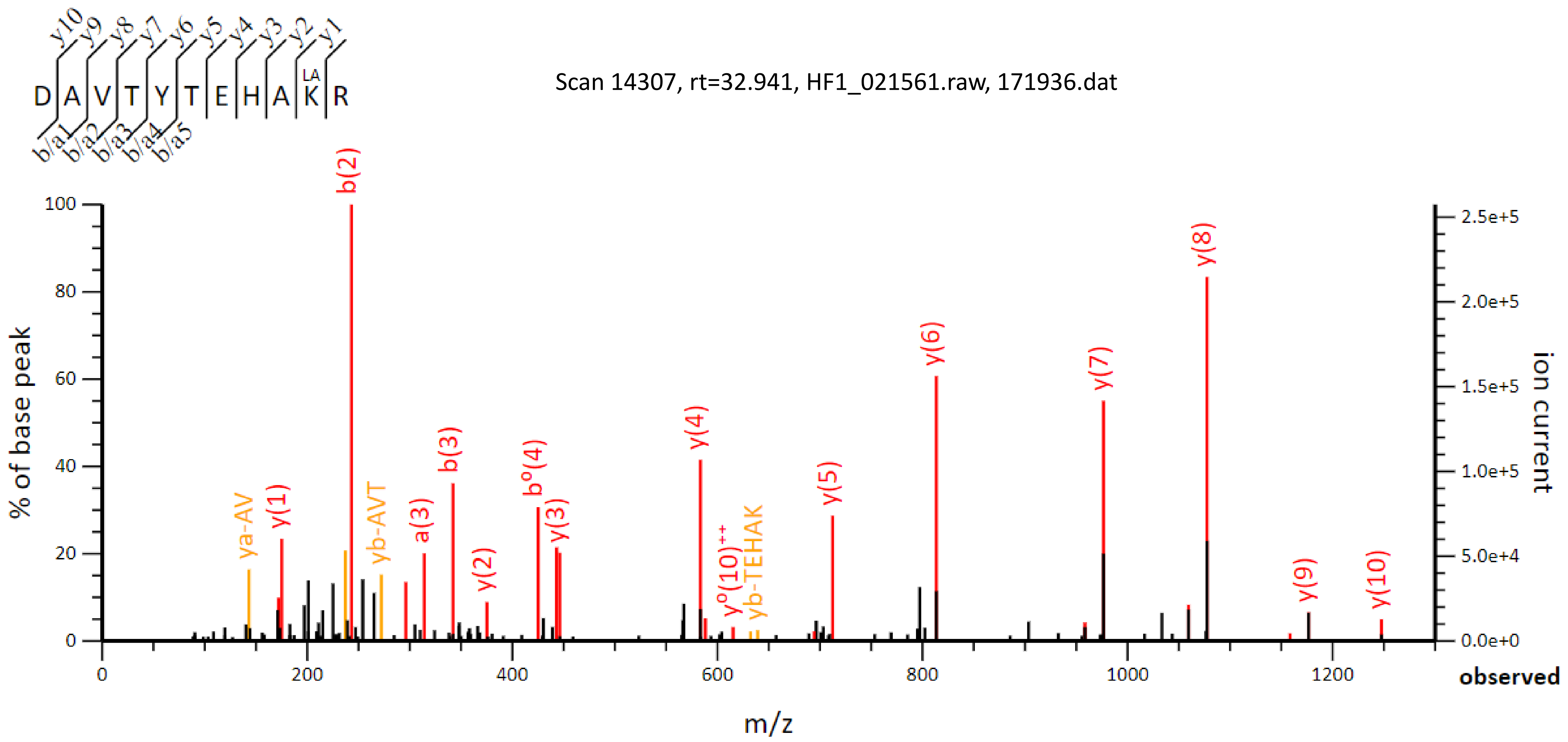

Scan 14307, rt=32.941, HF1_021561.raw, 171936.dat
D A V T Y T E H A K R
LA
y10 y9 y8 y7 y6 y5 y4 y3 y2 y1
b/a1 b/a2 b/a3 b/a4 b/a5
% of base peak
ion current
m/z
observed
b(2)
y(8)
y(6)
y(7)
y(4)
b(3)
bº(4)
y(5)
y(1)
ya-AV
yb-AVT
a(3)
y(3)
y(2)
yº(10)++
yb-TEHAK
y(9)
y(10)

### Oxidation events on histone H4 $K_{79}TVTAMDVVYALK_{91}R$ peptide, m/z=897.9871 (2+)

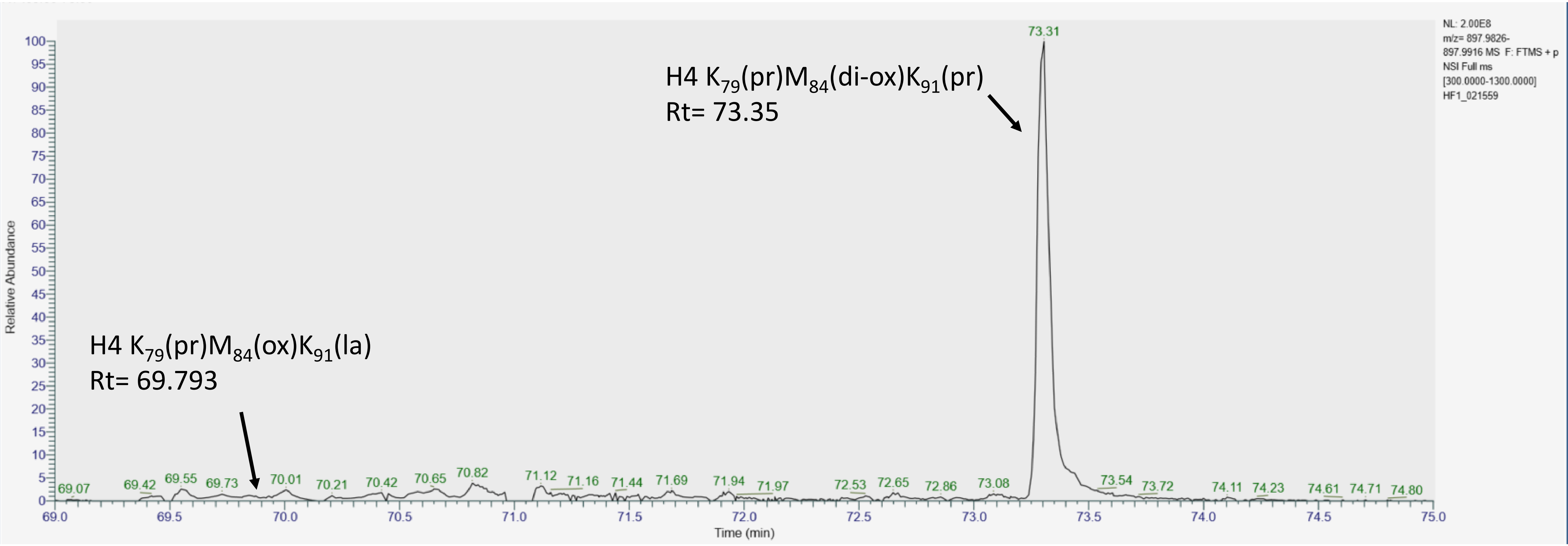

Scan 32593, rt=69.793, HF1_021559.raw, 172461.dat

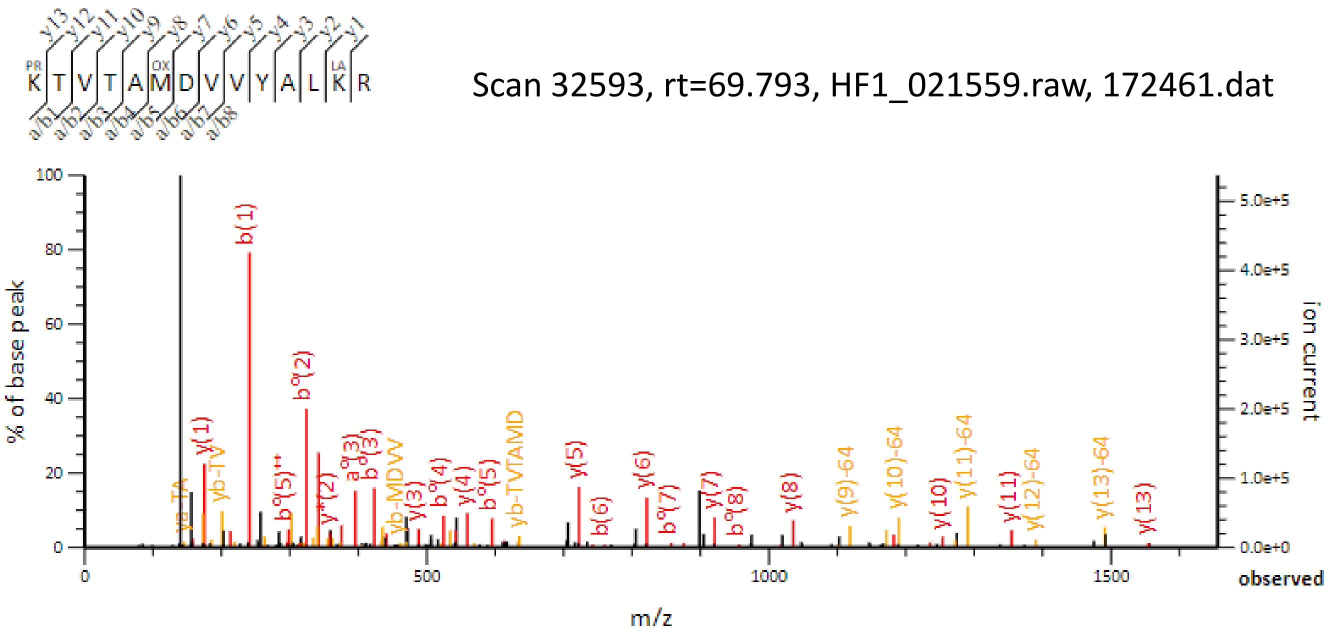


Scan 34357, rt=73.356, HF1_021559.raw, 172461.dat

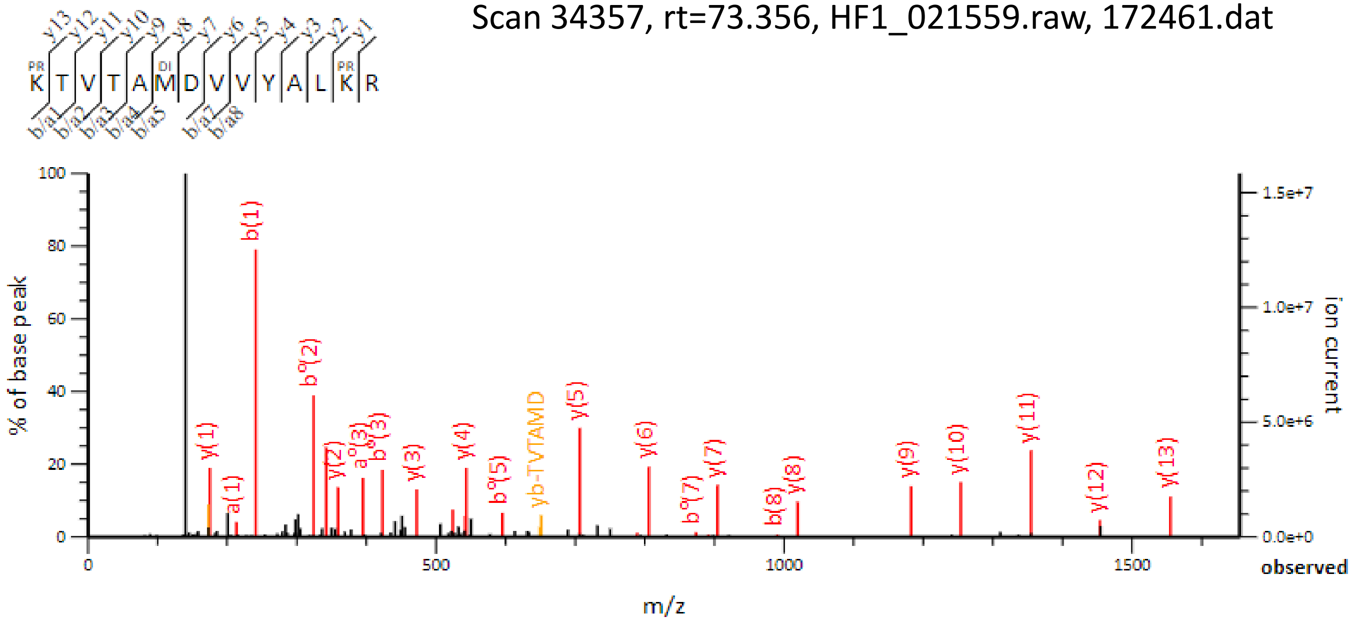

### Oxidation events on histone H3 YQK$_{56}$STELLIR peptide, m/z=689.8825 (2+)

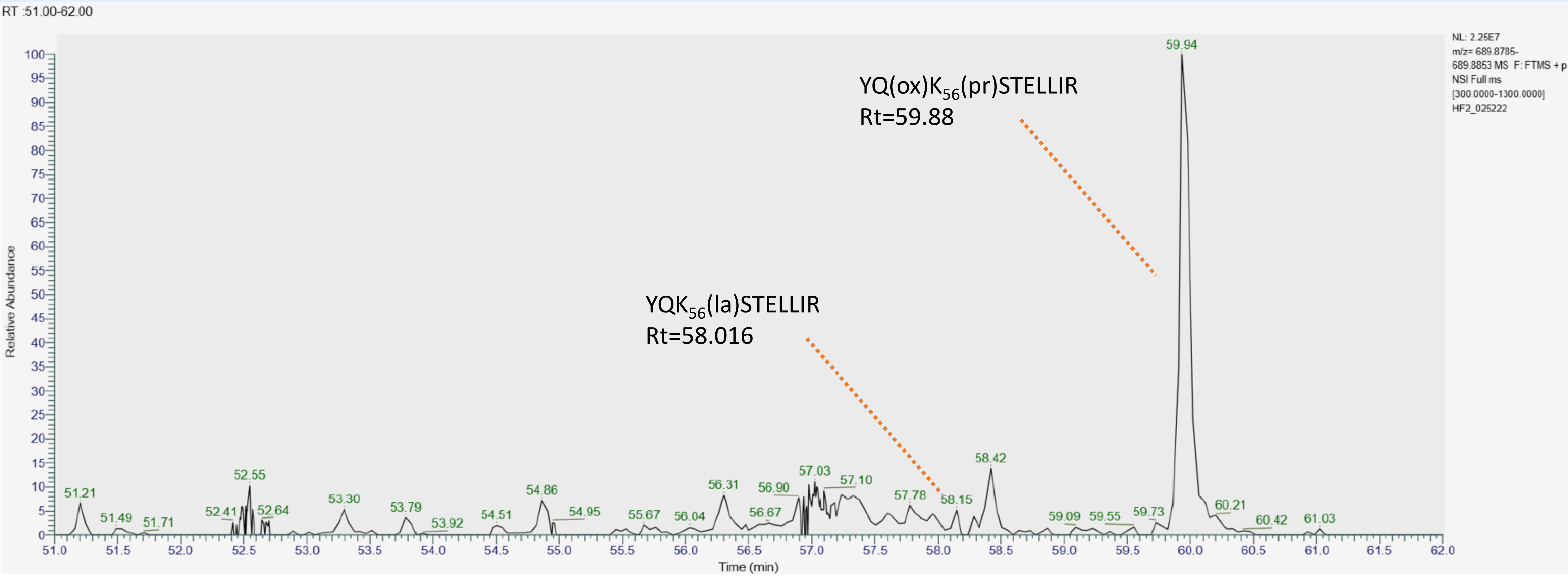

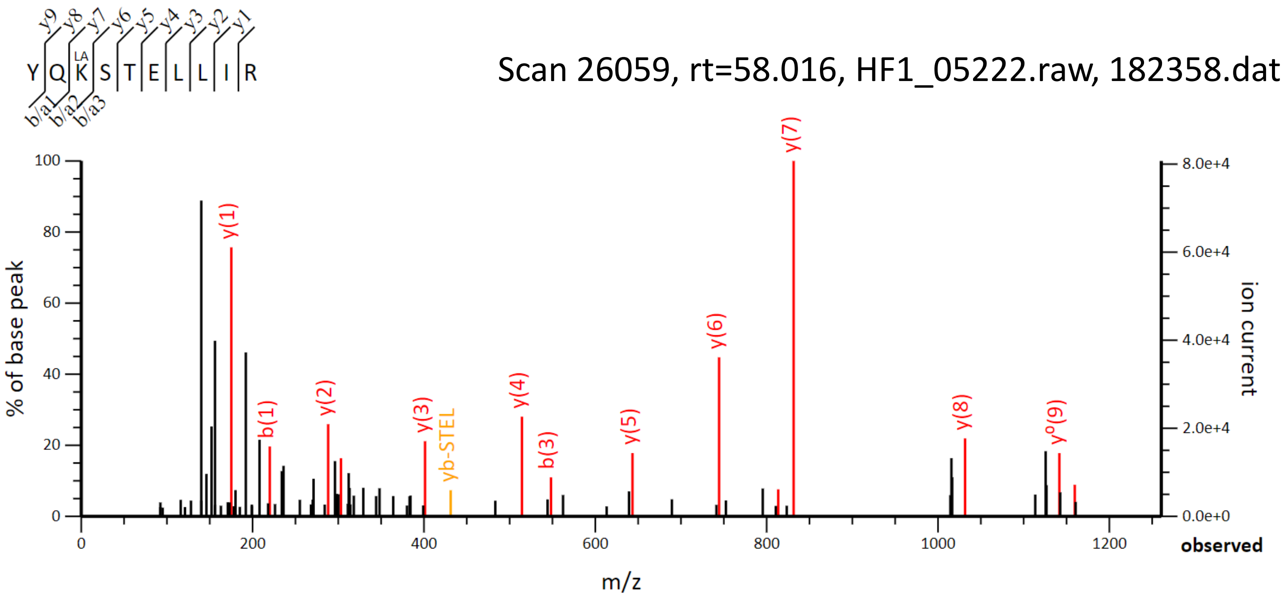

Scan 26059, rt=58.016, HF1_05222.raw, 182358.dat
Y Q K S T E L L I R
LA
% of base peak
ion current
m/z
observed
y(1)
b(1)
y(2)
y(3)
yb-STEL
y(4)
b(3)
y(5)
y(6)
y(7)
y(8)
y°(9)


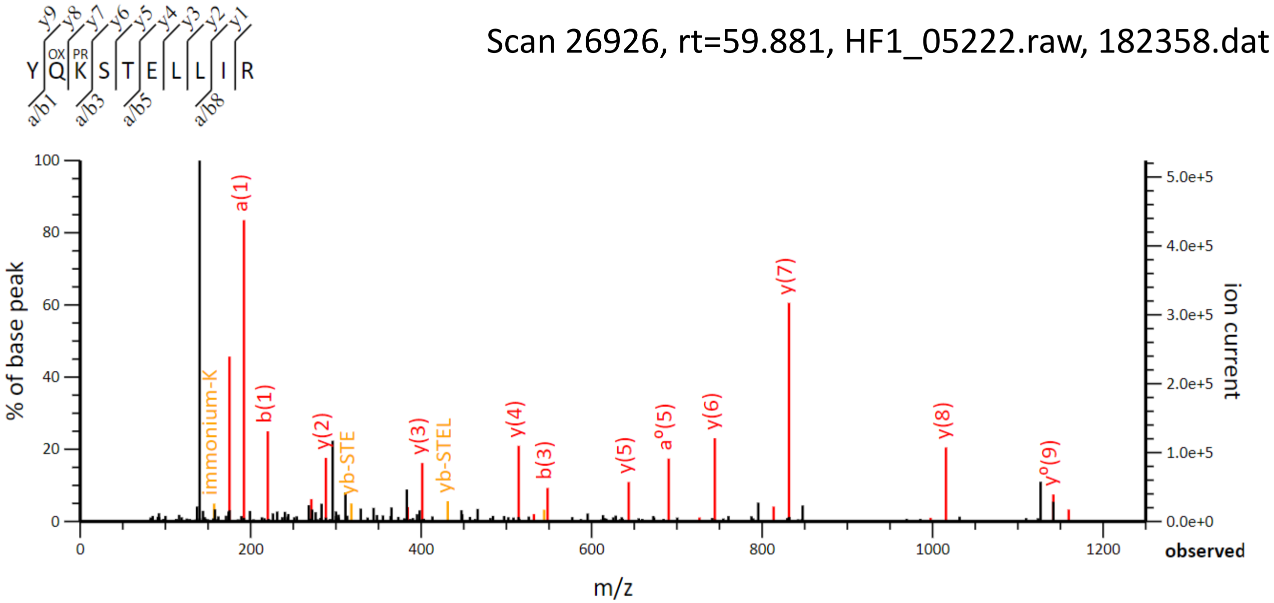

Scan 26926, rt=59.881, HF1_05222.raw, 182358.dat
Y Q K S T E L L I R
OX
PR
% of base peak
ion current
m/z
observed
immonium-K
a(1)
b(1)
y(2)
yb-STE
y(3)
yb-STEL
y(4)
b(3)
y(5)
a°(5)
y(6)
y(7)
y(8)
y°(9)

**Section 3**:
Determination of the chromatographic behavior of synthetic peptides from histones H3 and H4 bearing L- or D-lactylated lysines

#### Extracted ion chromatogram of H3.1 L- and D-lactyl $K_9STGGK_{14}APR$ (m/z=548.3052,2+)

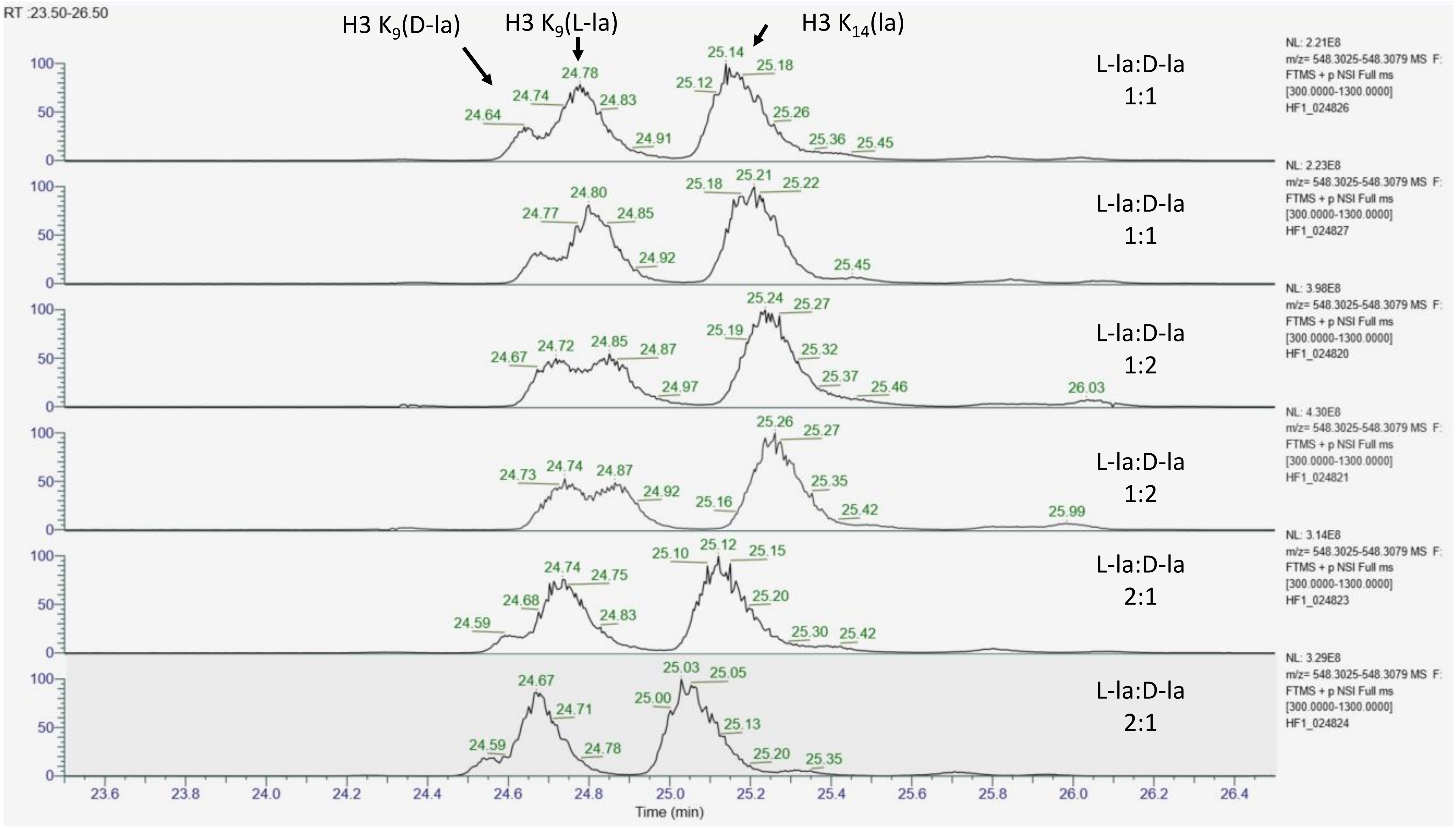

#### MS/MS spectra for H3 L- and D-lactyl $K_9STGGK_{14}APR$ in three different ratios

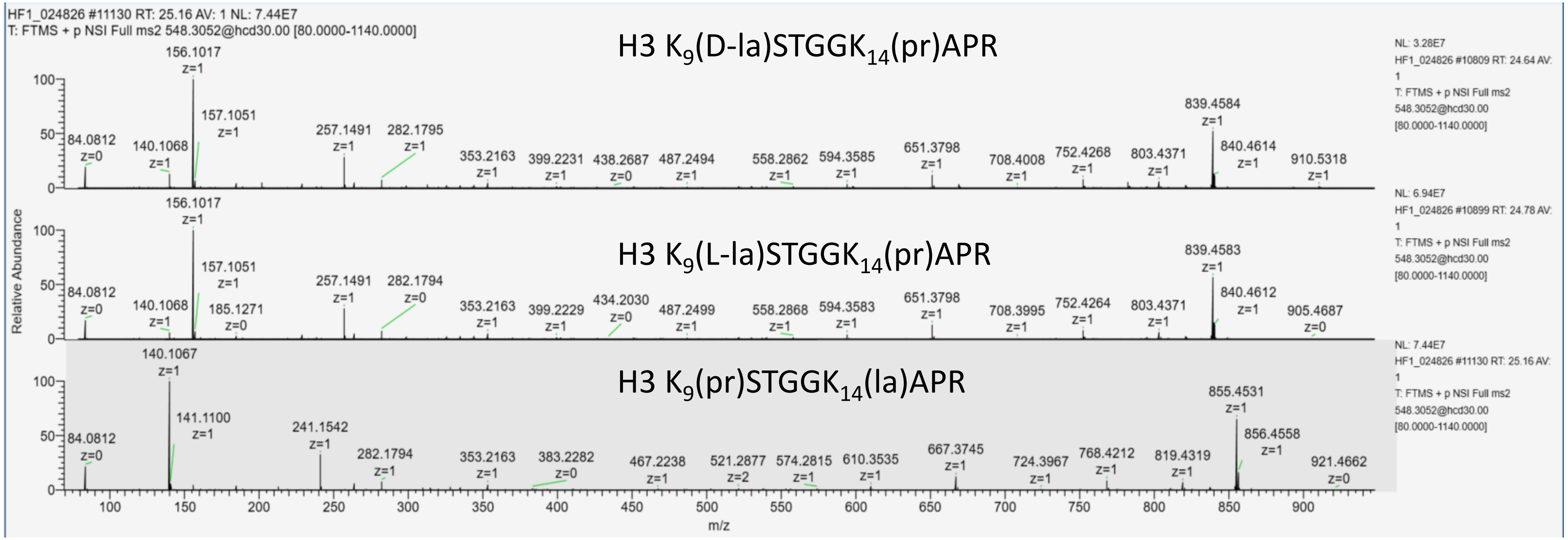


L-la:D-la
1:1

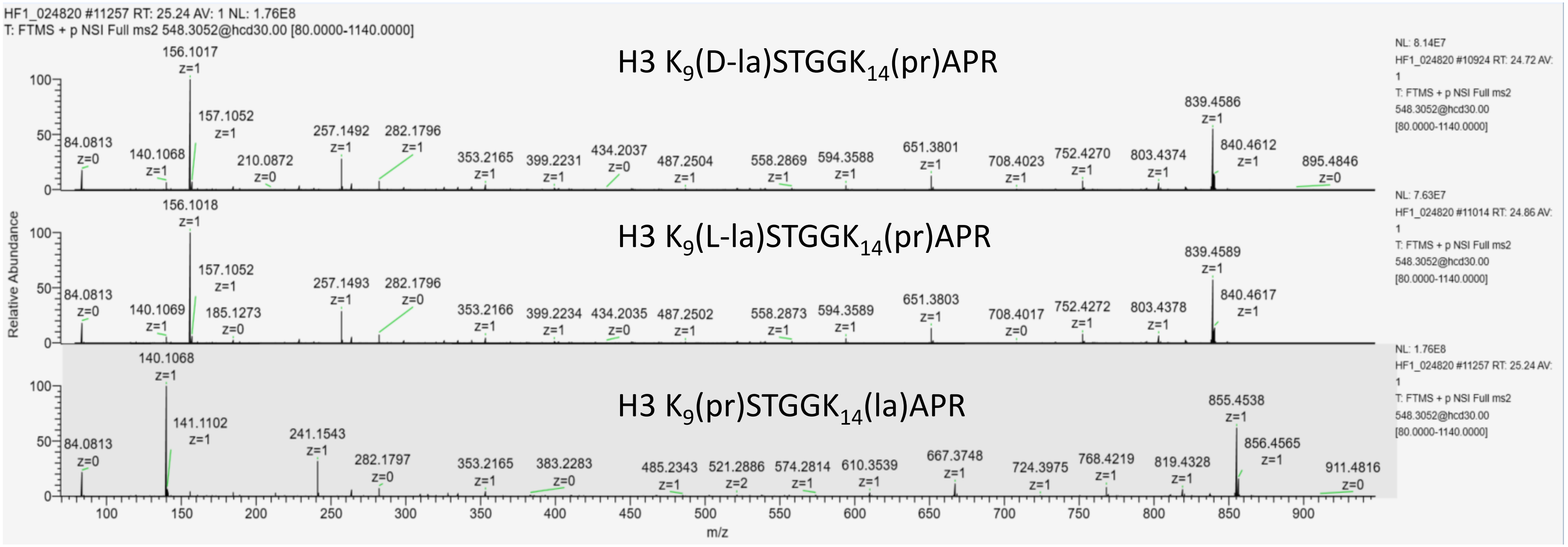


L-la:D-la
1:2

### MS/MS spectra for H3 L- and D-lactyl $K_9STGGK_{14}APR$ in three different ratios

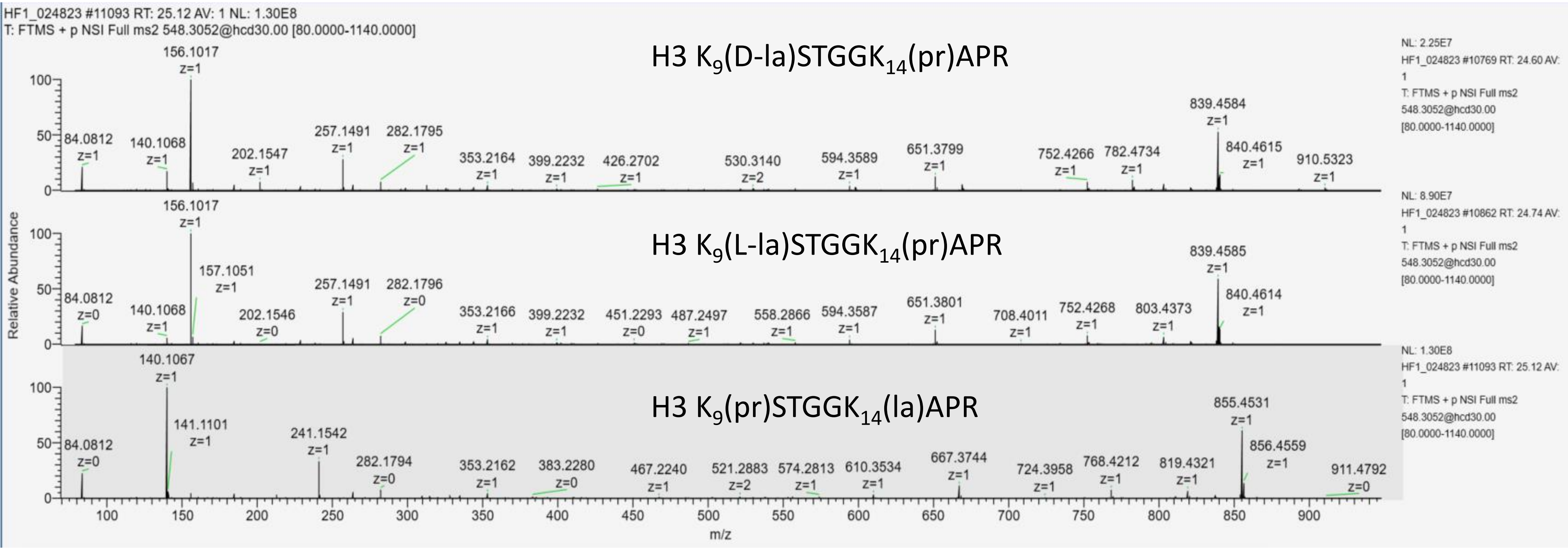


L-la:D-la
2:1

#### Extracted ion chromatogram of H3.1 L- and D-lactyl $K_{18}$QLATK$_{23}$AAR (m/z=590.8498,2+)

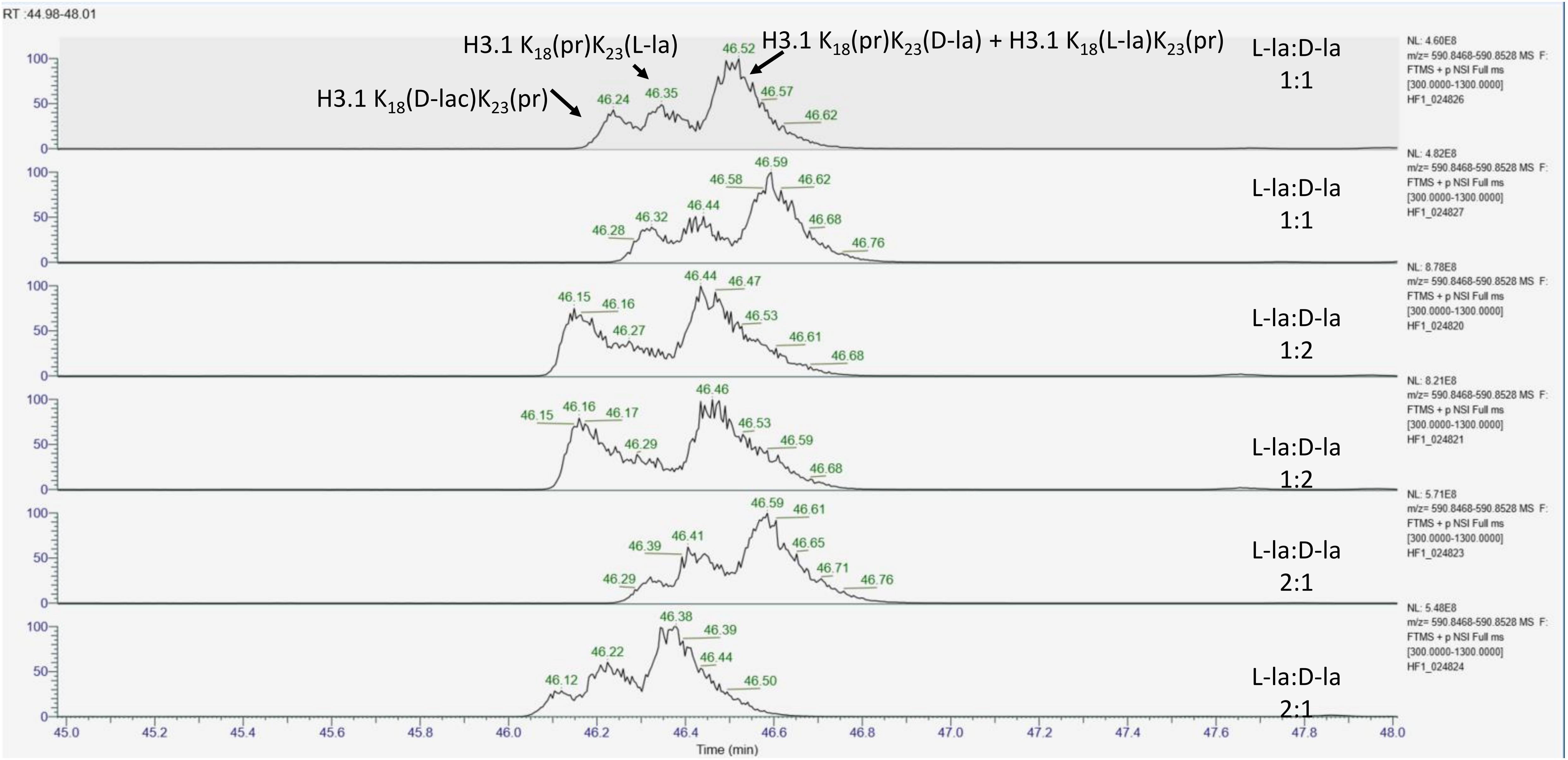

### MS/MS spectra acquired at apex of each chromatographic peak for H3.1 L- and D-lactyl $K_{18}QLATK_{23}AAR$ in three injections

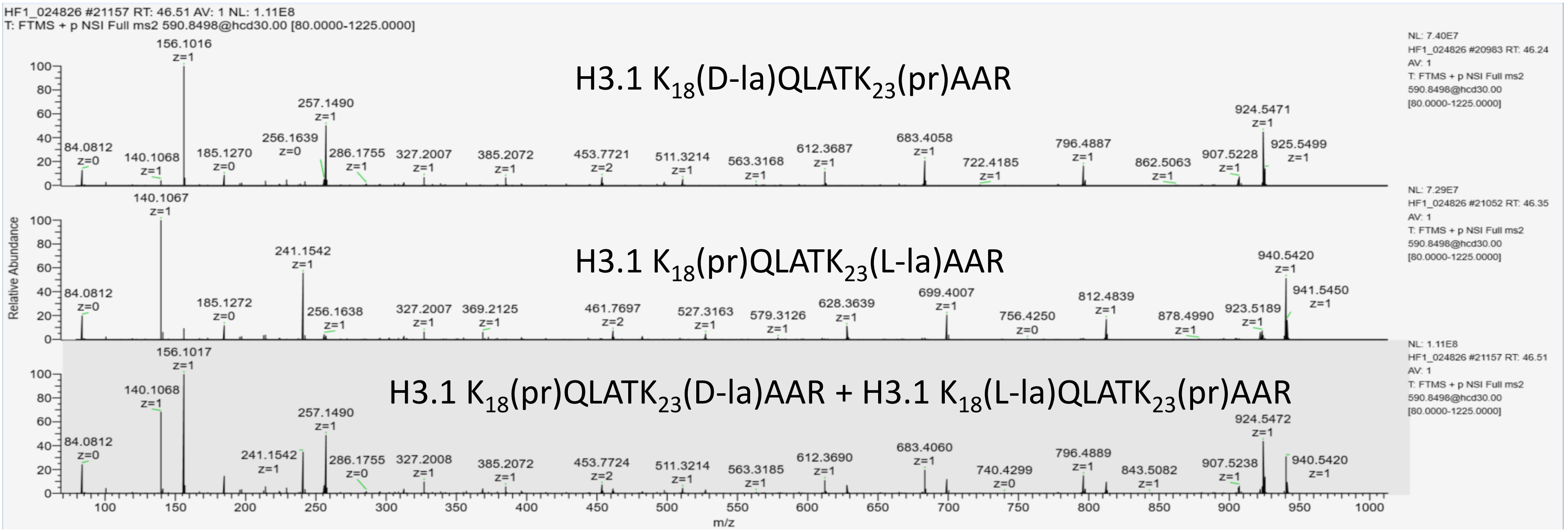


L-la:D-la
1:1

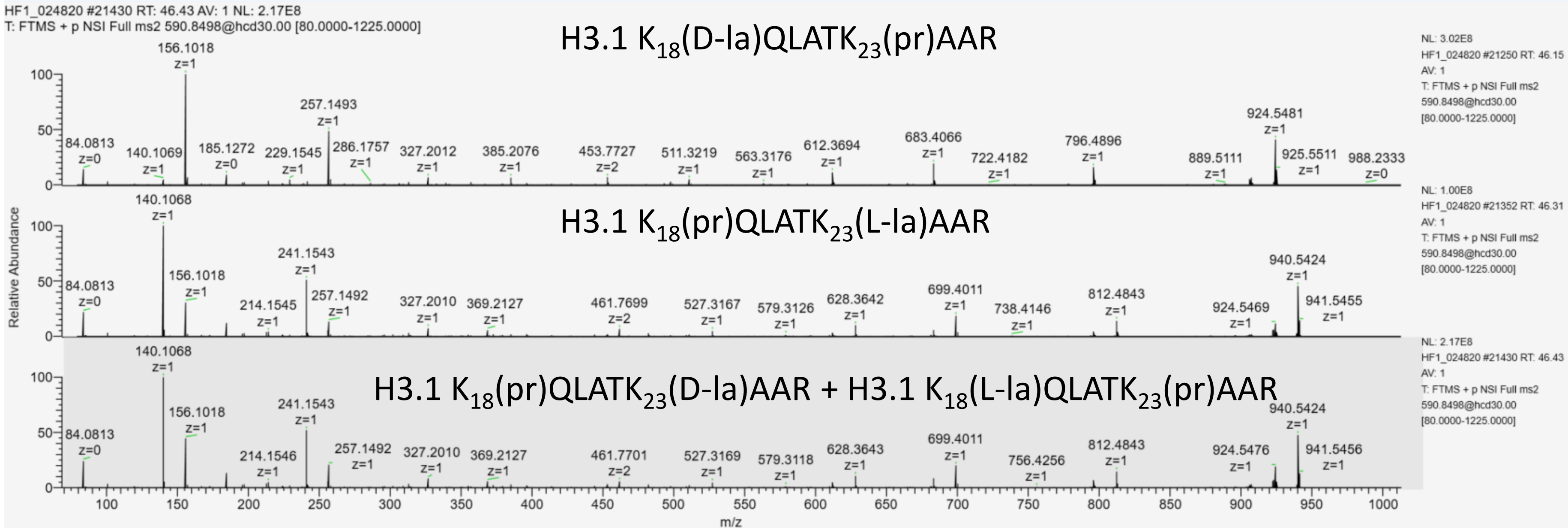


L-la:D-la
1:2

**MS/MS spectra acquired at apex of each chromatographic peak for H3.1 L- and D-lactyl $K_{18}QLATK_{23}AAR$**

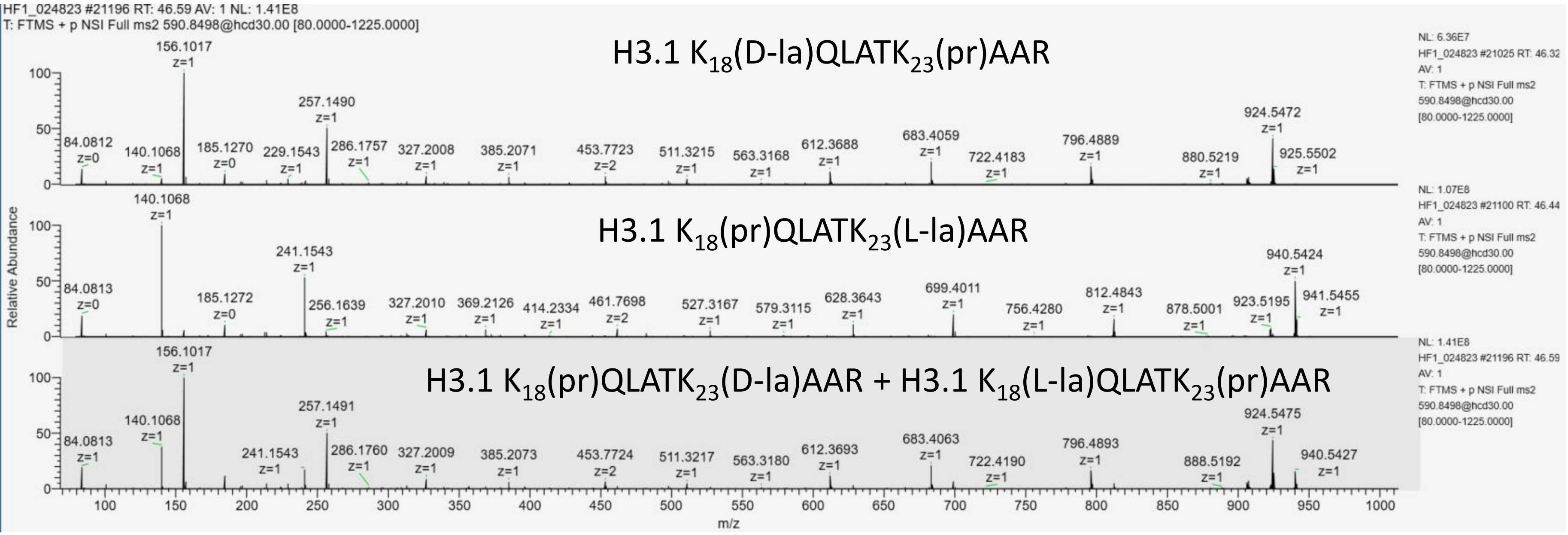


L-la:D-la

2:1

**Extracted ion chromatogram of H3.t L- and D-lactyl $K_{18}QLATK_{23}VAR$ (m/z=604.8655,2+)**

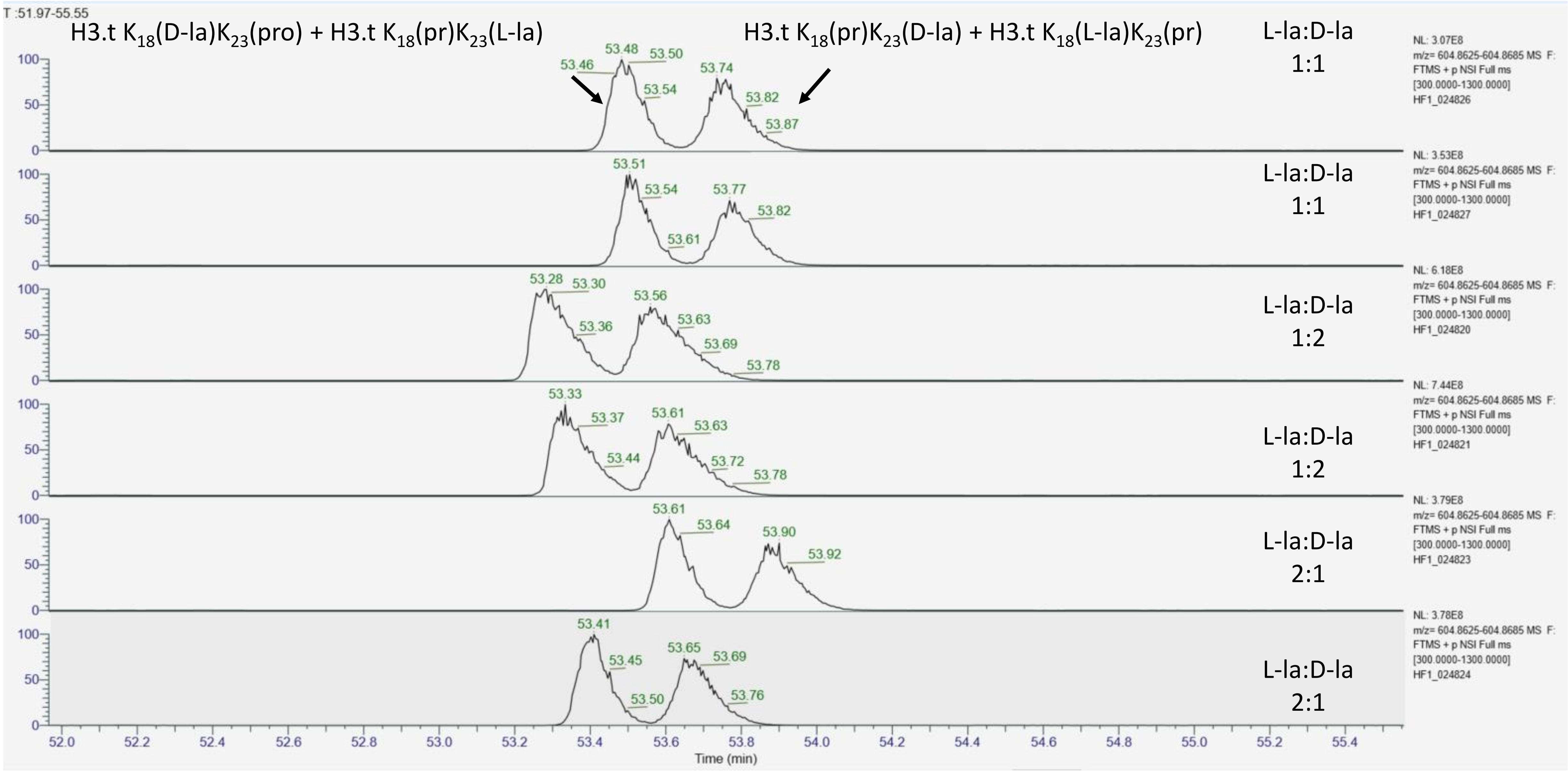

### MS/MS spectra acquired at apex of each chromatographic peak for H3.t L- and D-lactyl $K_{18}QLATK_{23}VAR$ in three injections

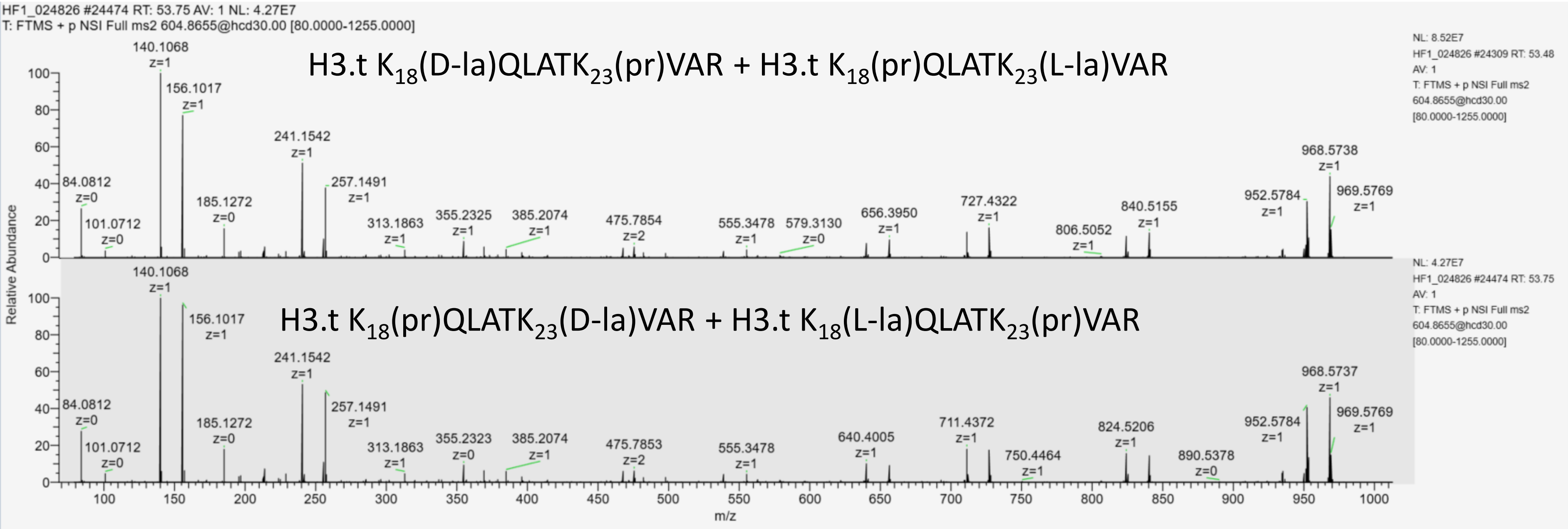


L-la:D-la
1:1

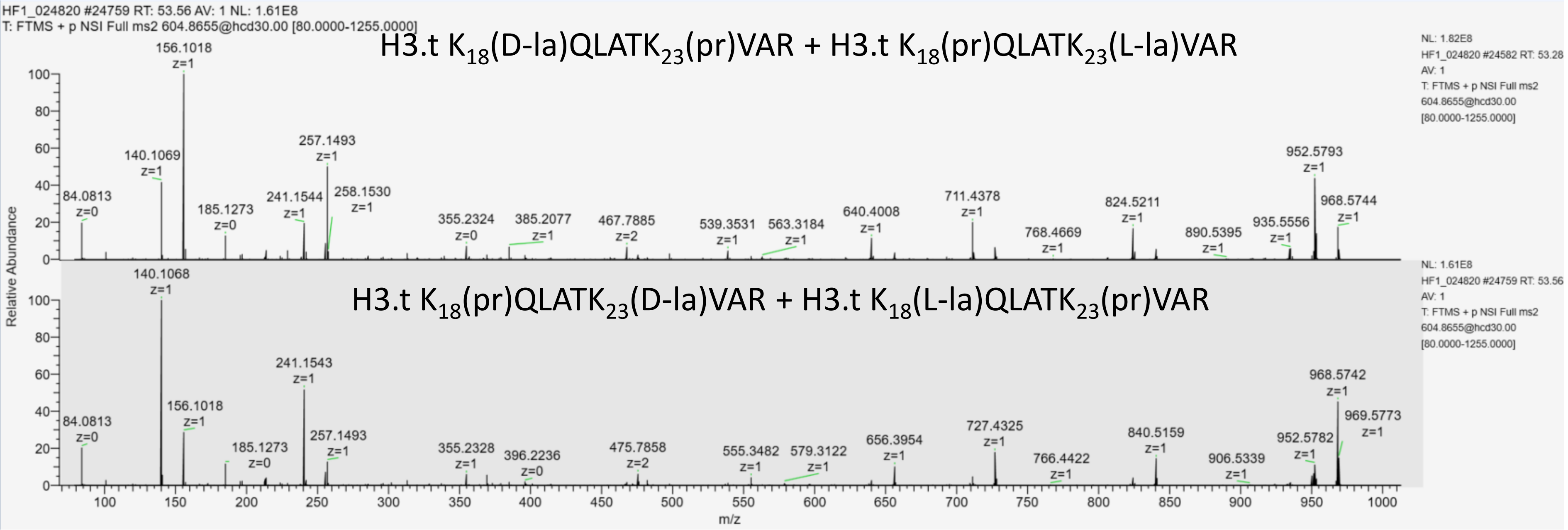


L-la:D-la
1:2

#### MS/MS spectra acquired at apex of each chromatographic peak for H3.t L- and D-lactyl $K_{18}QLATK_{23}VAR$

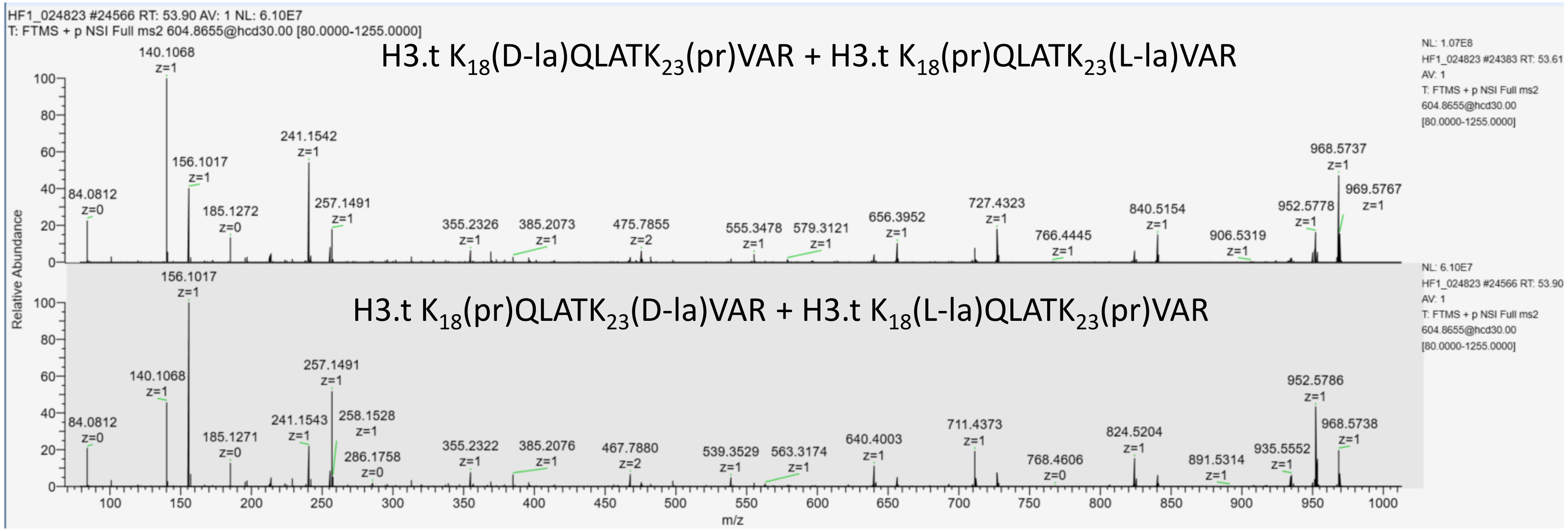


L-la:D-la
2:1

### Extracted ion chromatogram of H3.1 $K_{27}$(la)SAPATGGV$K_{36}$(pr)$K_{37}$(pro)PHR (m/z=842.4744,2+)

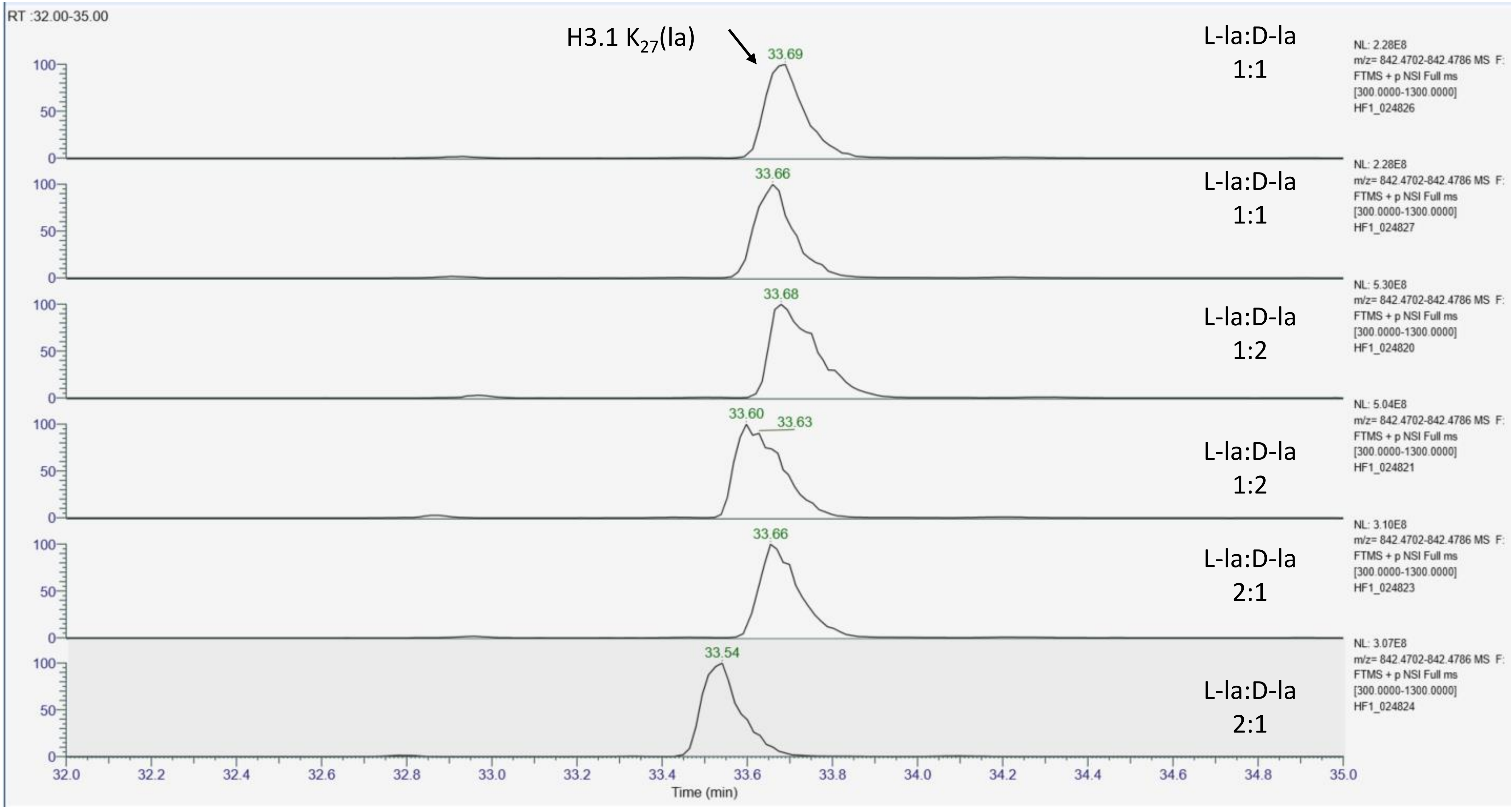

#### MS/MS spectra acquired at the apex of each chromatographic peak for H3.1 $K_{27}$(la)SAPATGGV$K_{36}$(pr)$K_{37}$(pr)PHR in three injections

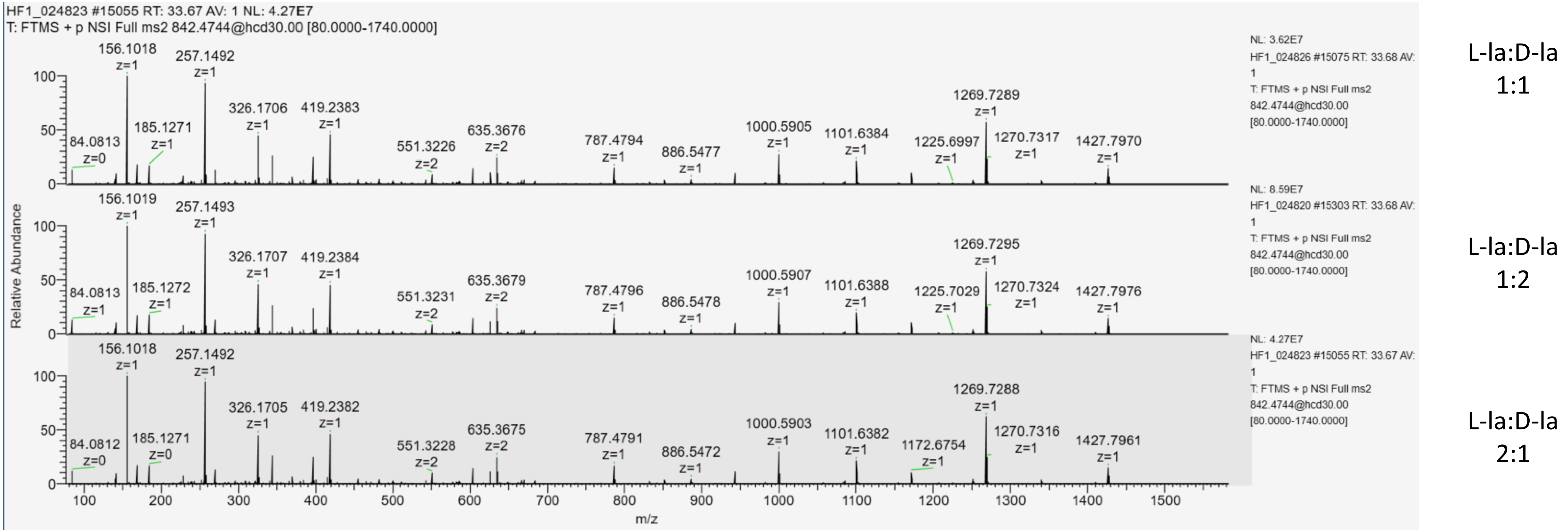

#### Extracted ion chromatogram of H3.3 $K_{27}$(la)SAPSTGV$K_{37}$(pr)$K_{37}$(pr)PHR (m/z=850.4719,2+)

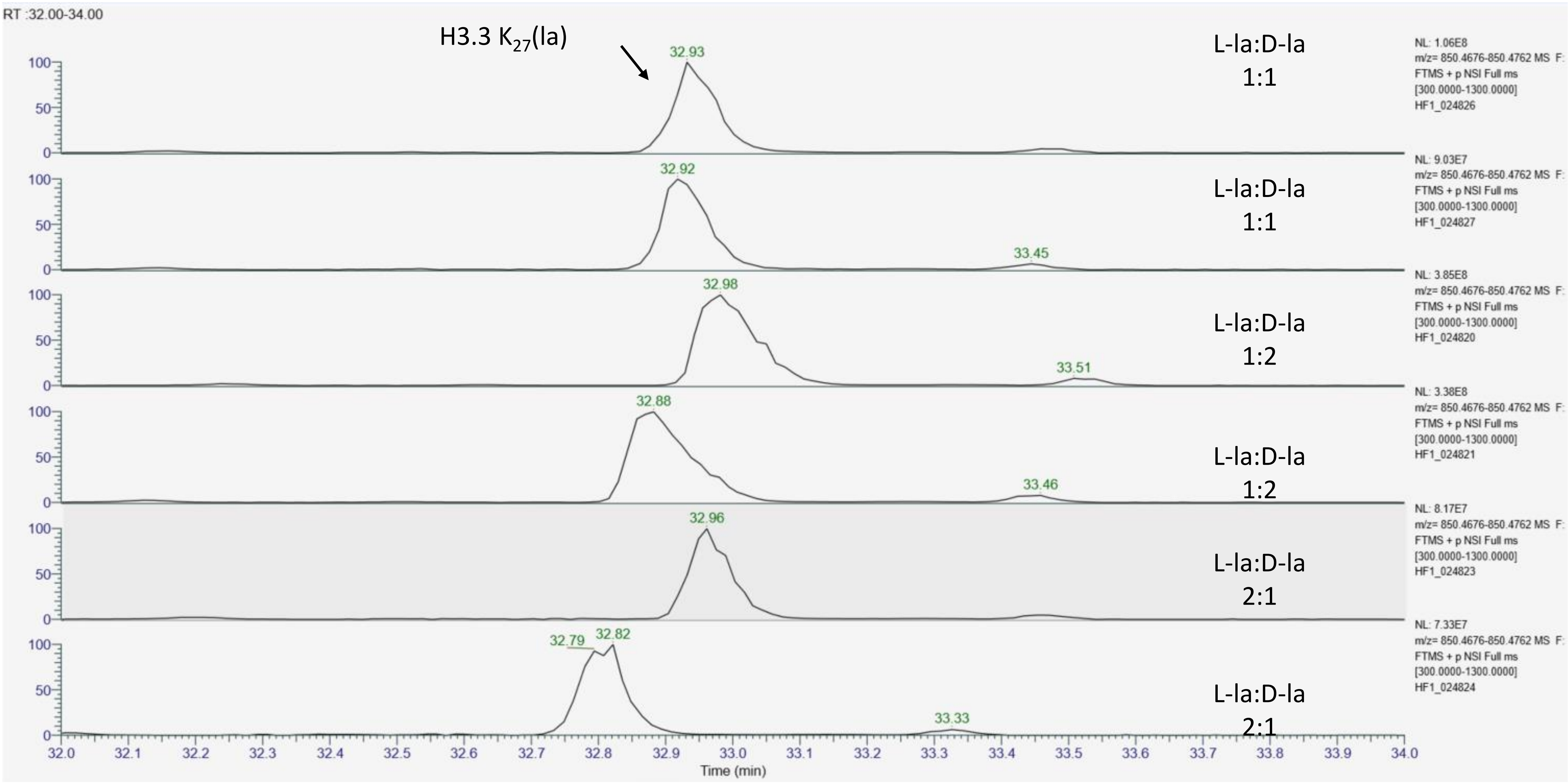

#### MS/MS spectra acquired at the apex of chromatographic peaks for H3.3 $K_{27}$(la)SAPSTGVK$_{37}$(pr)K$_{37}$(pr)PHR in three injections

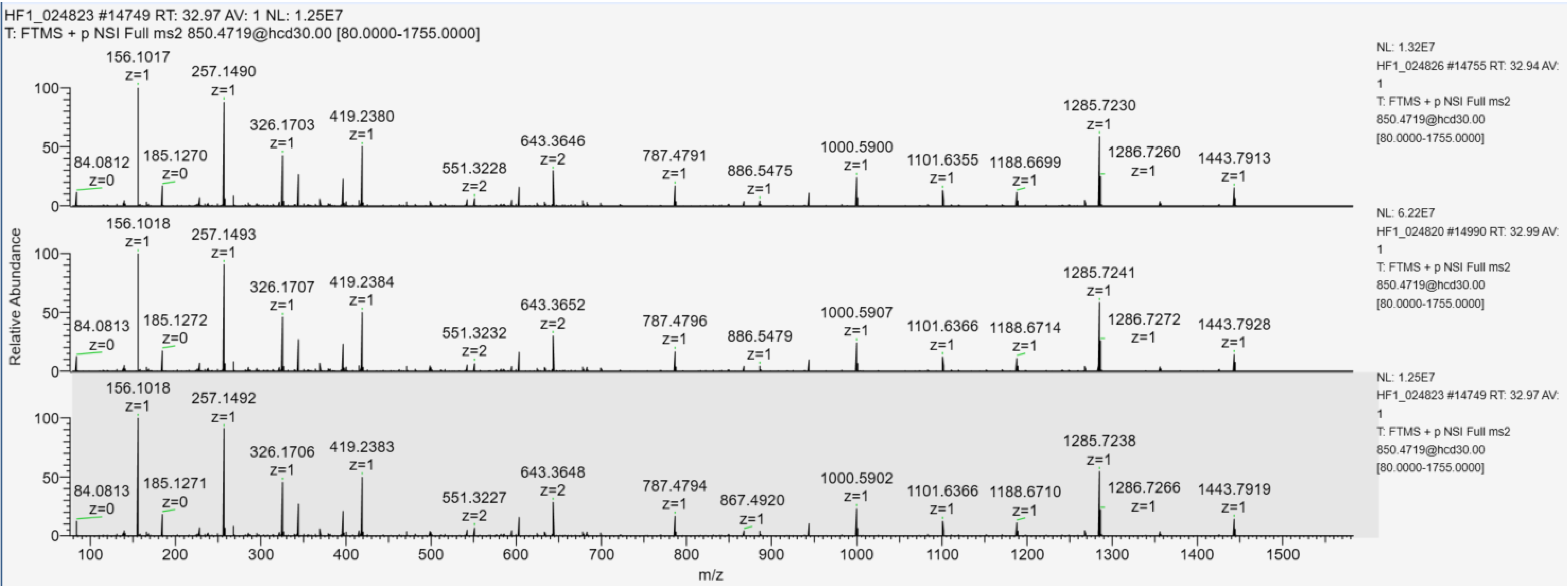


L-la:D-la 1:1

L-la:D-la 1:2

L-la:D-la 2:1

### Extracted ion chromatogram of L- and D-lactylated H3 YQK$_{56}$STELLIR (m/z=694.8866,2+)

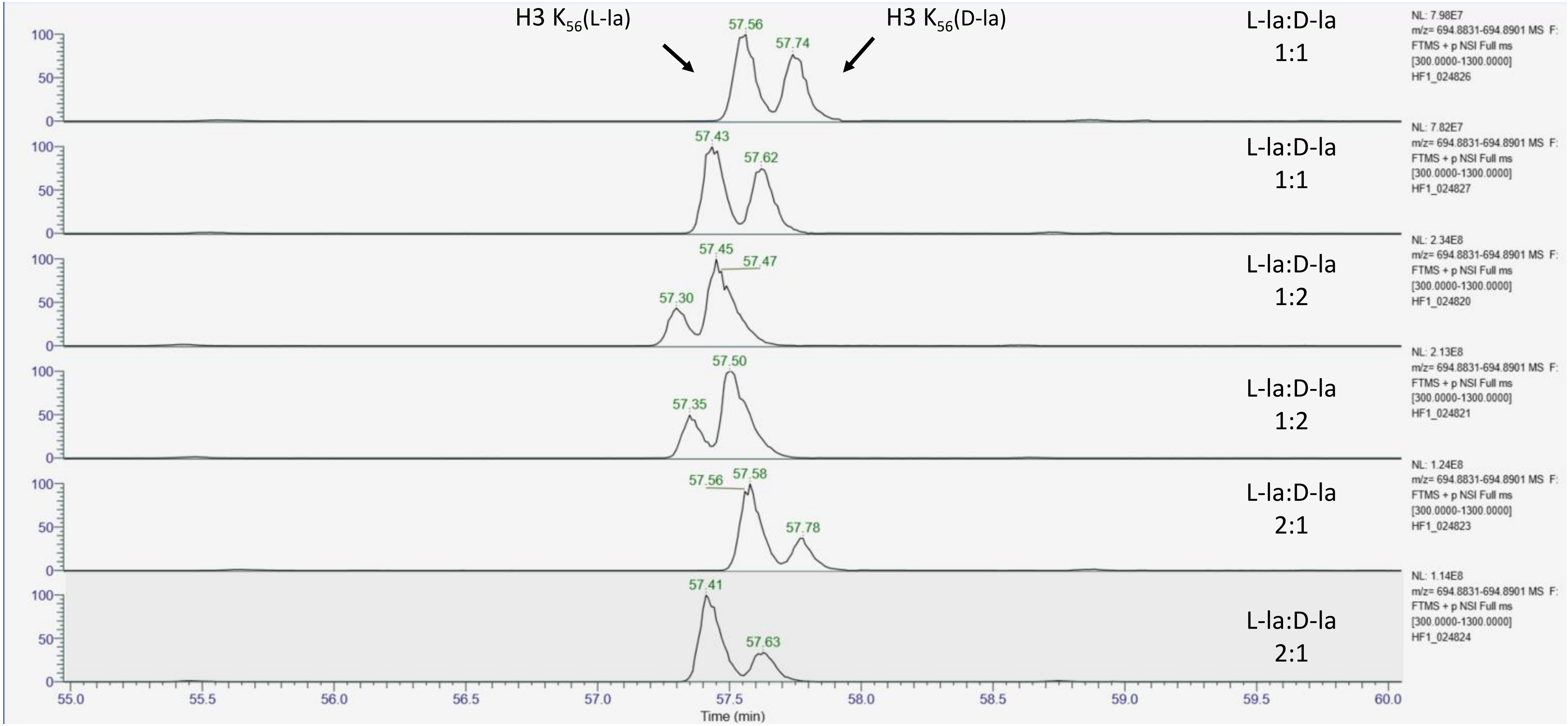

### MS/MS spectra acquired at apex of each chromatographic peak for L- and D-lactylated H3 YQK$_{56}$STELLIR

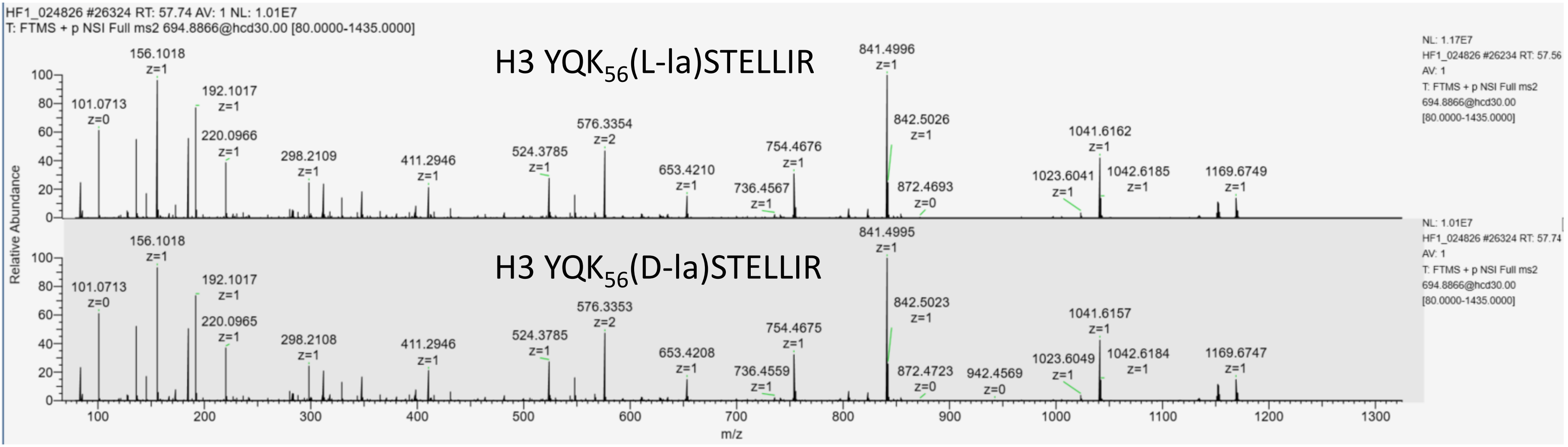


L-la:D-la
1:1

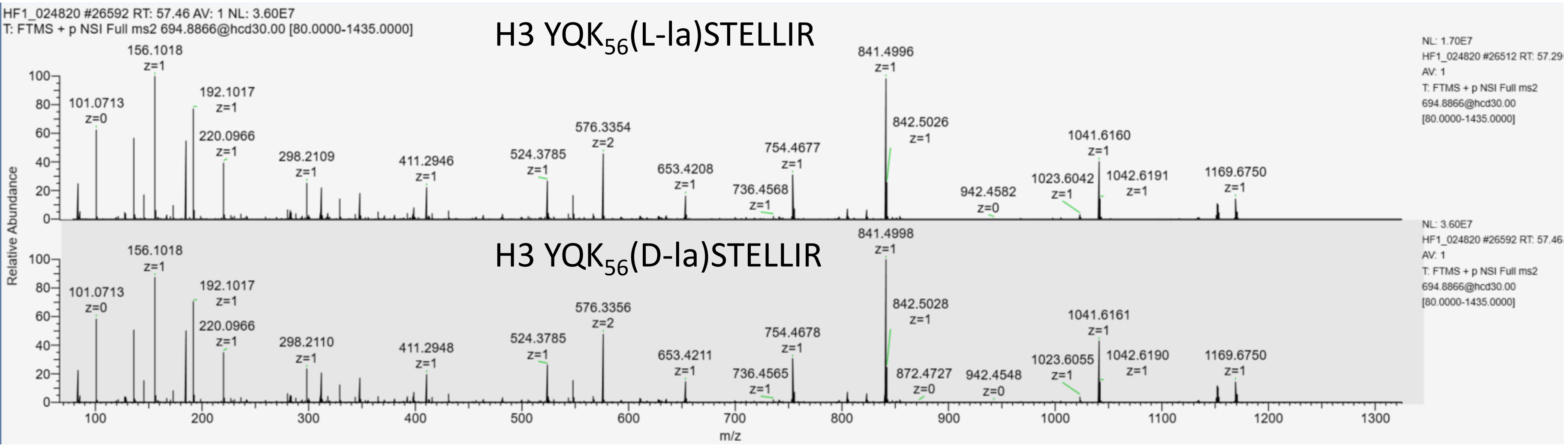


L-la:D-la
1:2

### MS/MS spectra acquired at apex of each chromatographic peak for L- and D-lactylated H3 YQK$_{56}$STELLIR

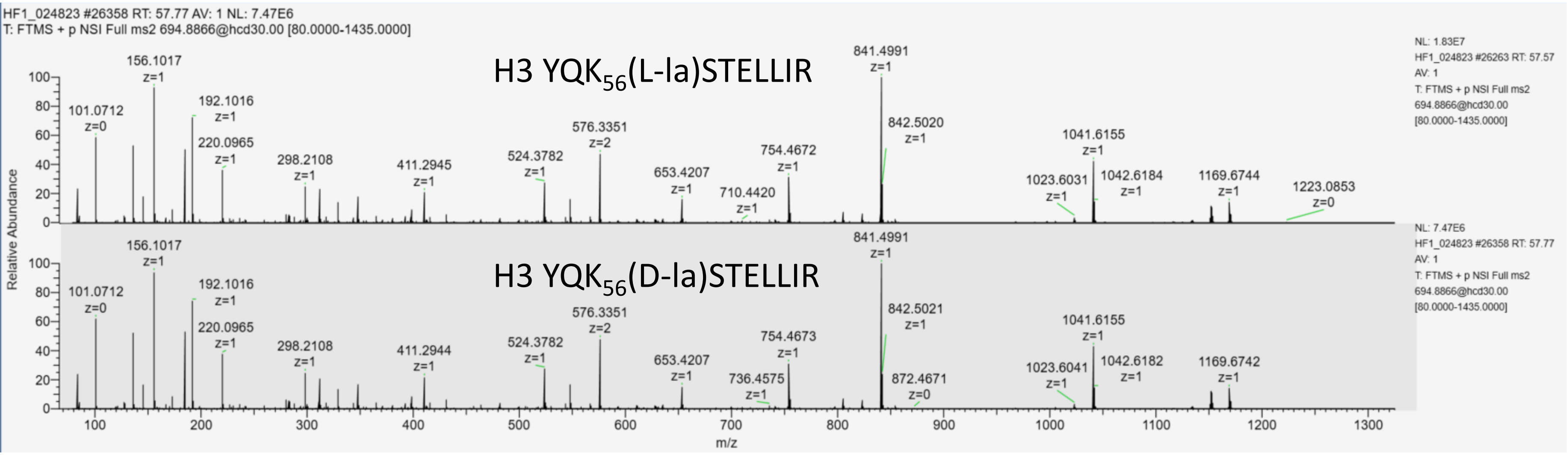


L-la:D-la
2:1

### Extracted ion chromatogram of H3 $K_{64}$(la)LPFQR (m/z=463.7703,2+)

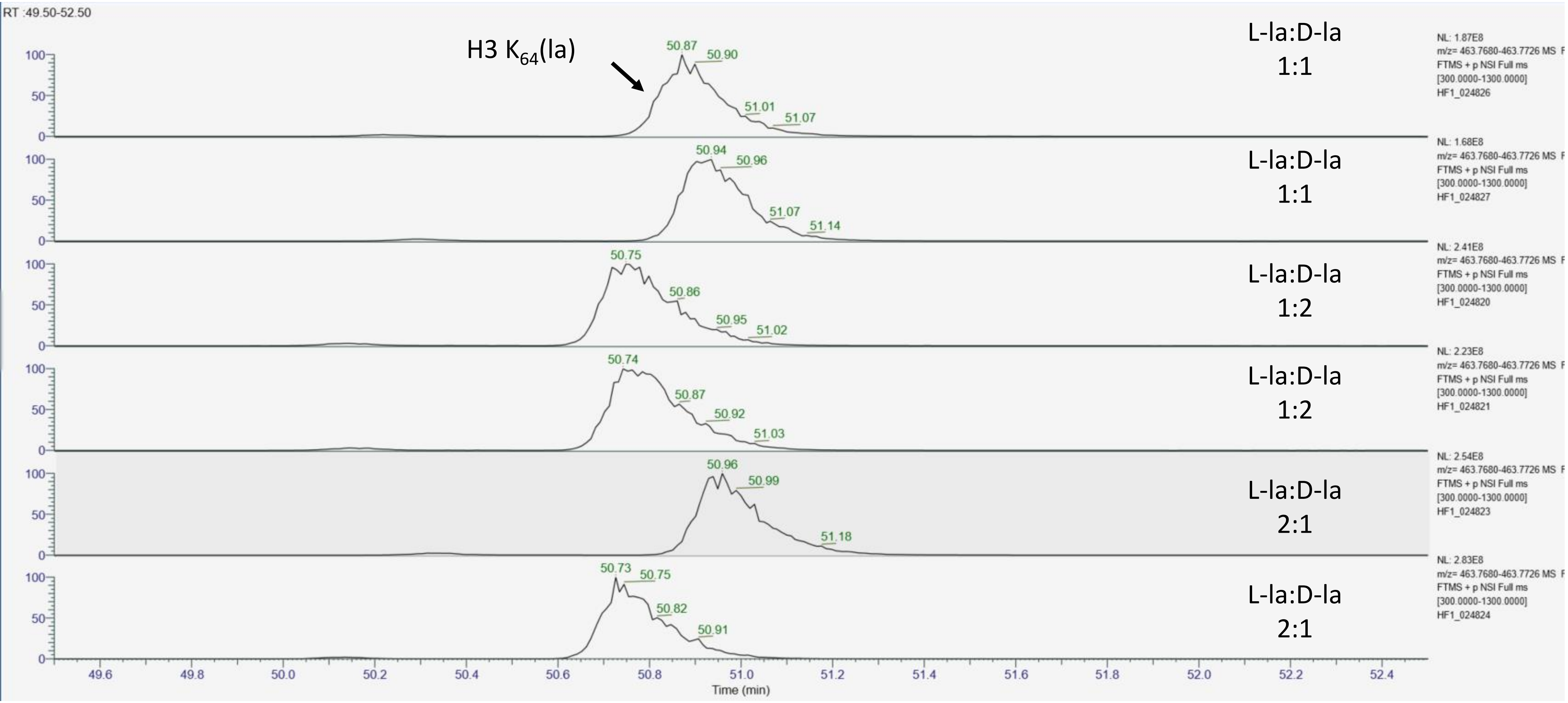

#### MS/MS spectra acquired at apex of each chromatographic peak for H3 $K_{64}$(la)LPFQR

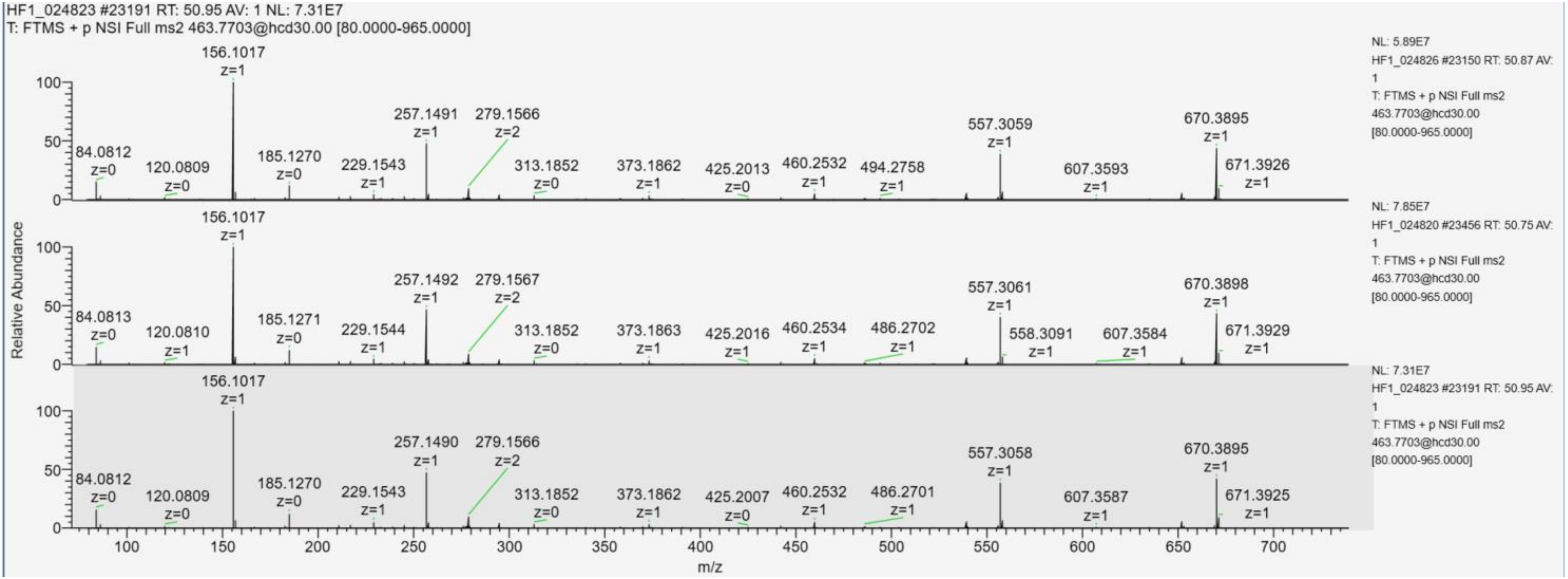


L-la:D-la 1:1

L-la:D-la 1:2

L-la:D-la 2:1

**Extracted ion chromatogram of L- and D-lactylated H3 EIAQDFK$_{79}$TDLR (m/z=737.3766,2+)**

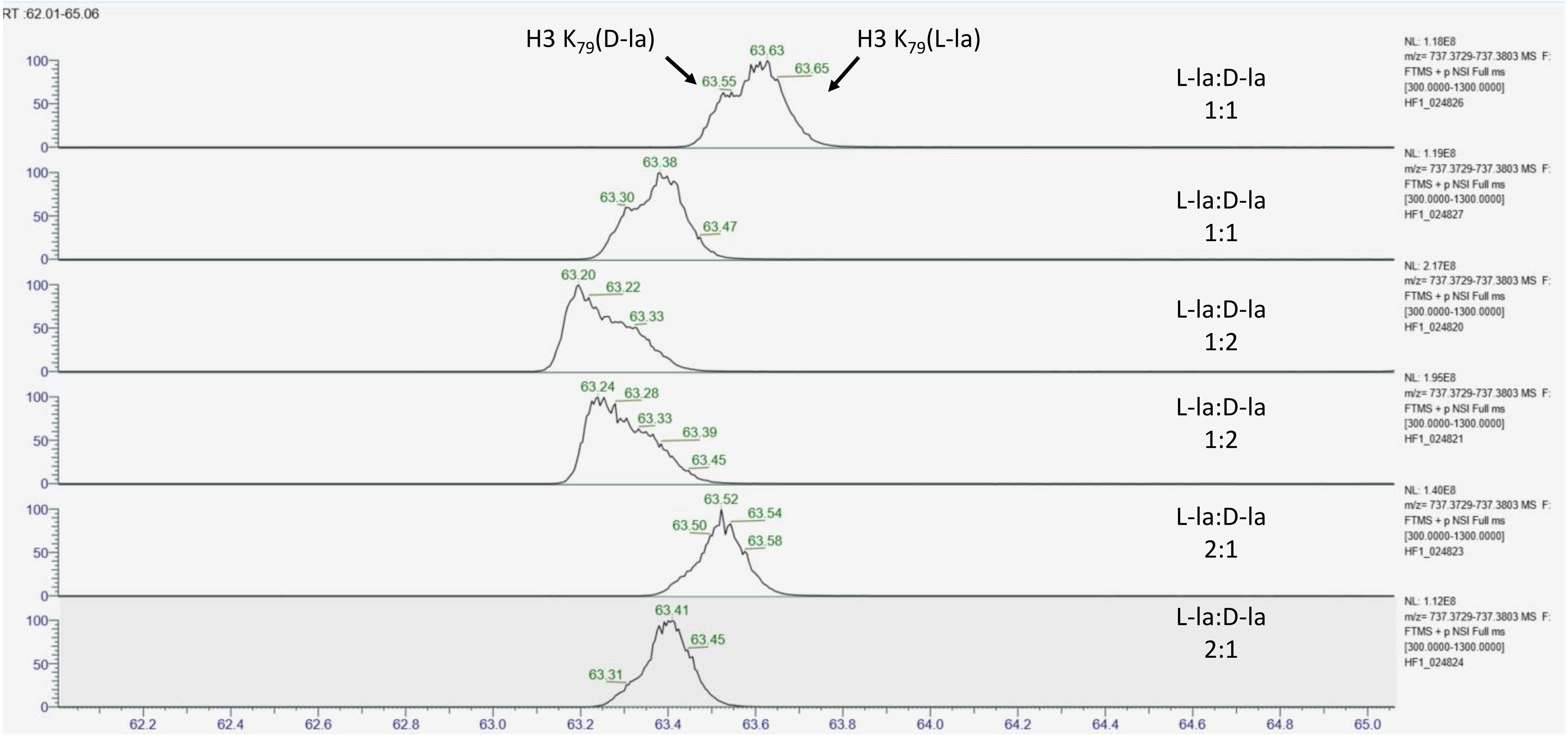

### MS/MS spectra acquired at apex of each chromatographic peak for L- and D-lactylated H3 EIAQDFK$_{79}$TDLR

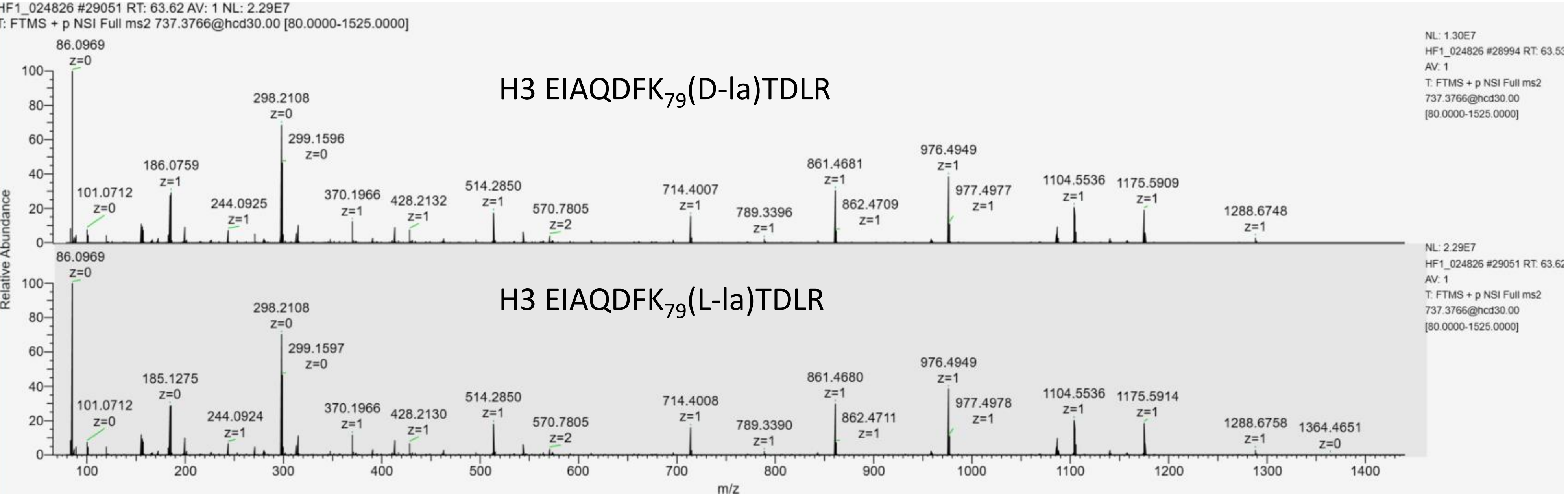


L-la:D-la
1:1

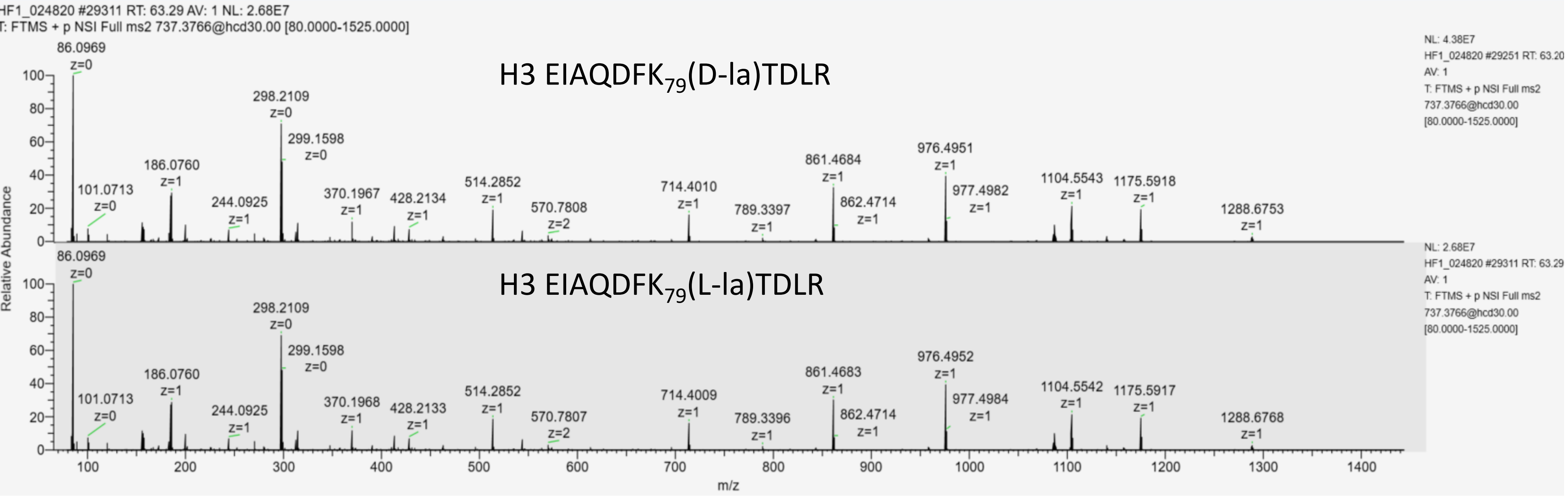


L-la:D-la
1:2

### MS/MS spectra acquired at apex of each chromatographic peak for L- and D-lactylated H3 EIAQDFK$_{79}$TDLR

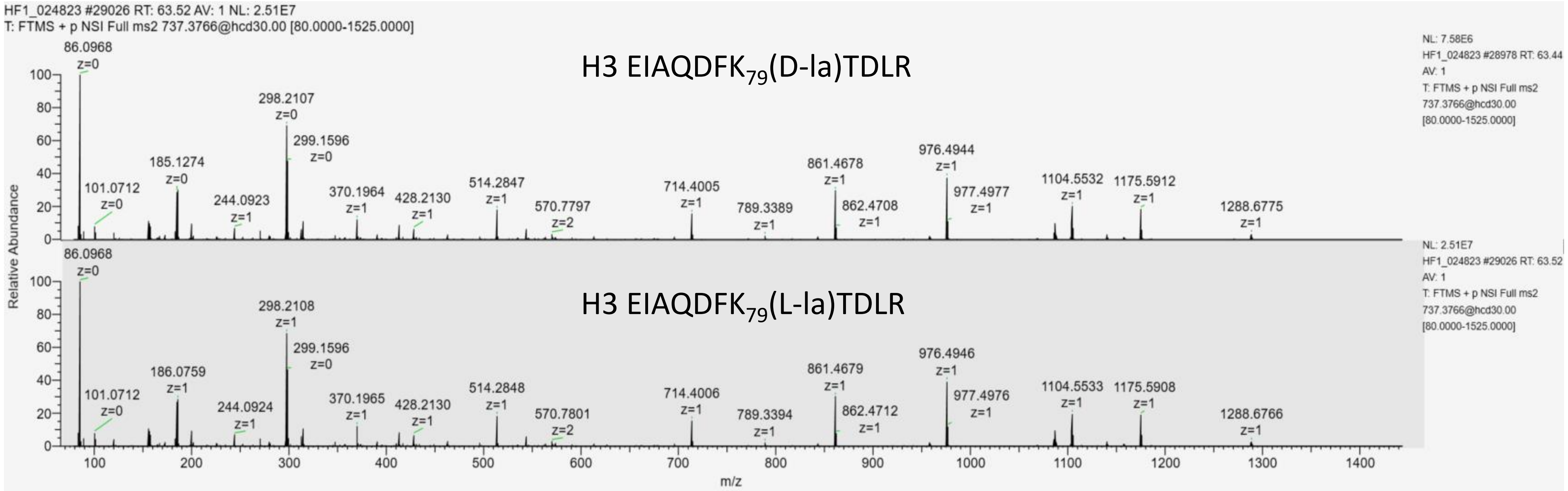


L-la:D-la
1:2

### Extracted ion chromatogram of H3 VTIMPK$_{122}$(la)DIQLAR (761.9305,2+)

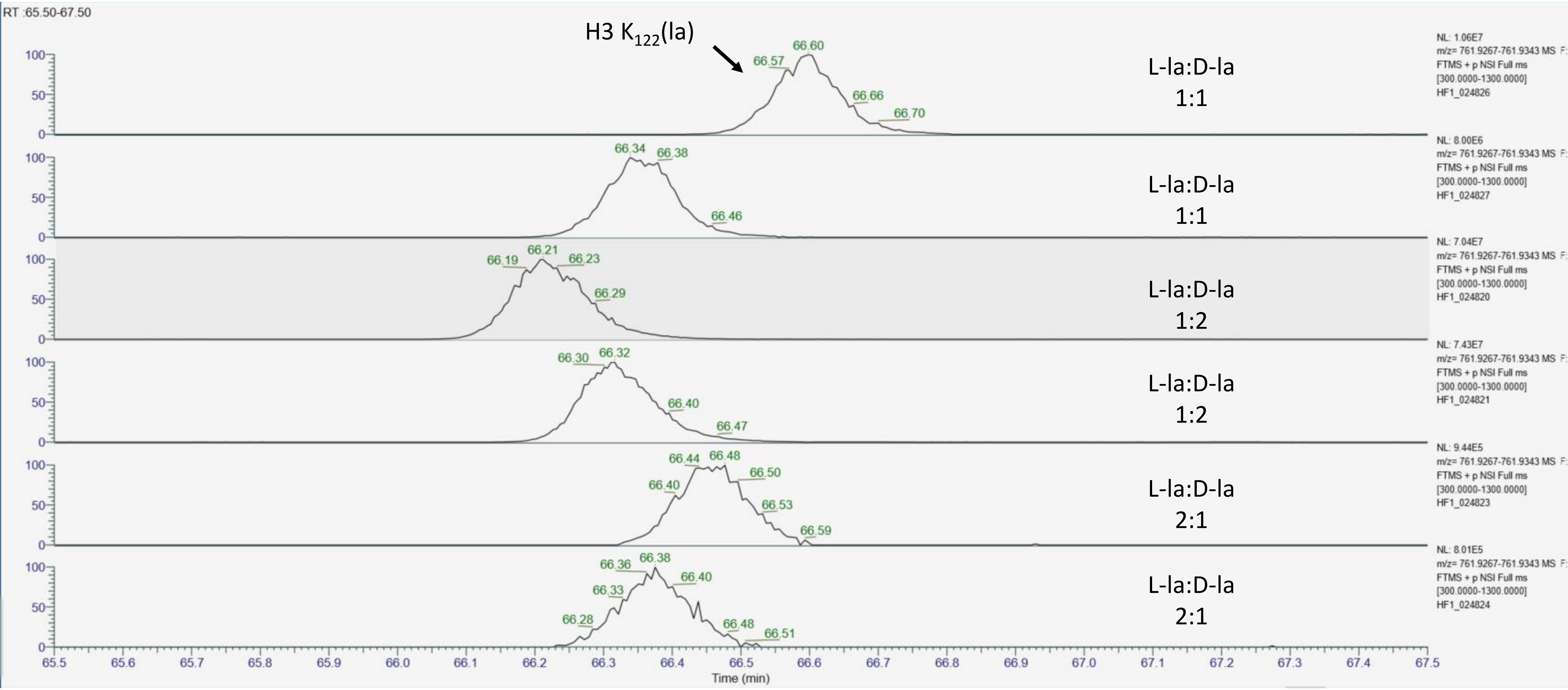

### MS/MS spectra acquired at apex of each chromatographic peak for H3 VTIMPK$_{122}$(la)DIQLAR in three injections

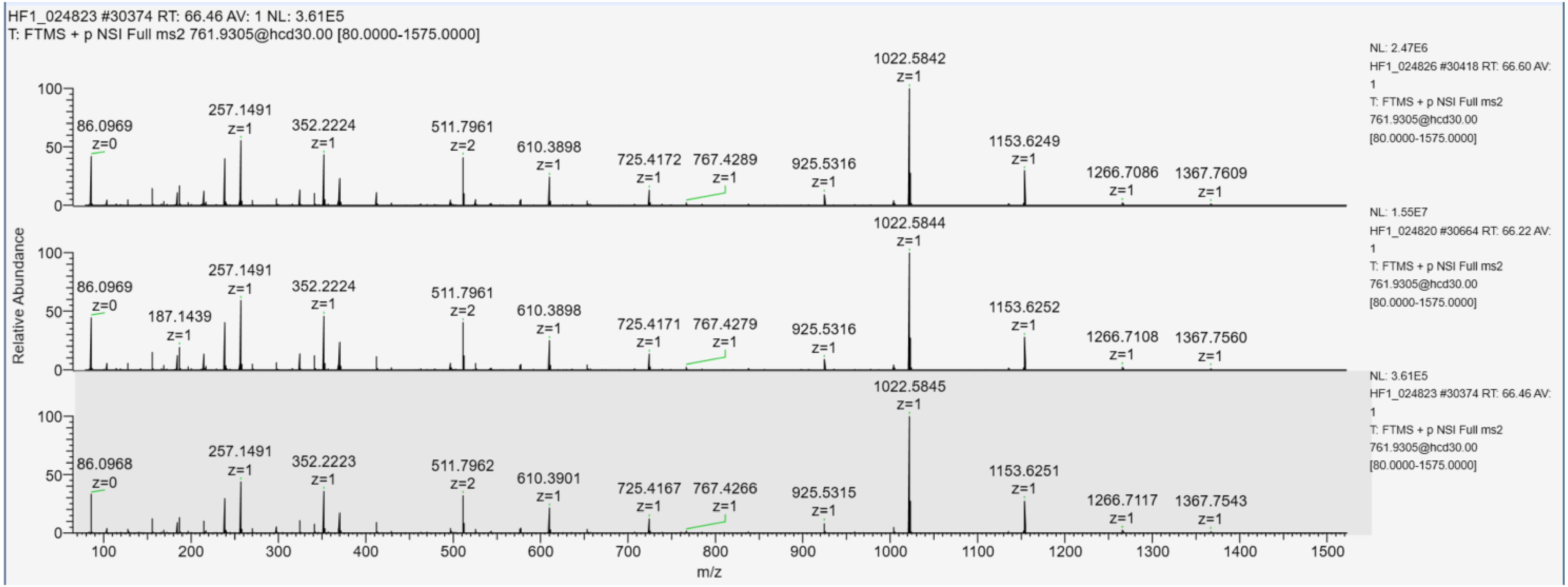


L-la:D-la 1:1

L-la:D-la 1:2

L-la:D-la 2:1

### Extracted ion chromatogram of H3 VTIM(ox)PK$_{122}$(la)DIQLAR (m/z=769.9279,2+)

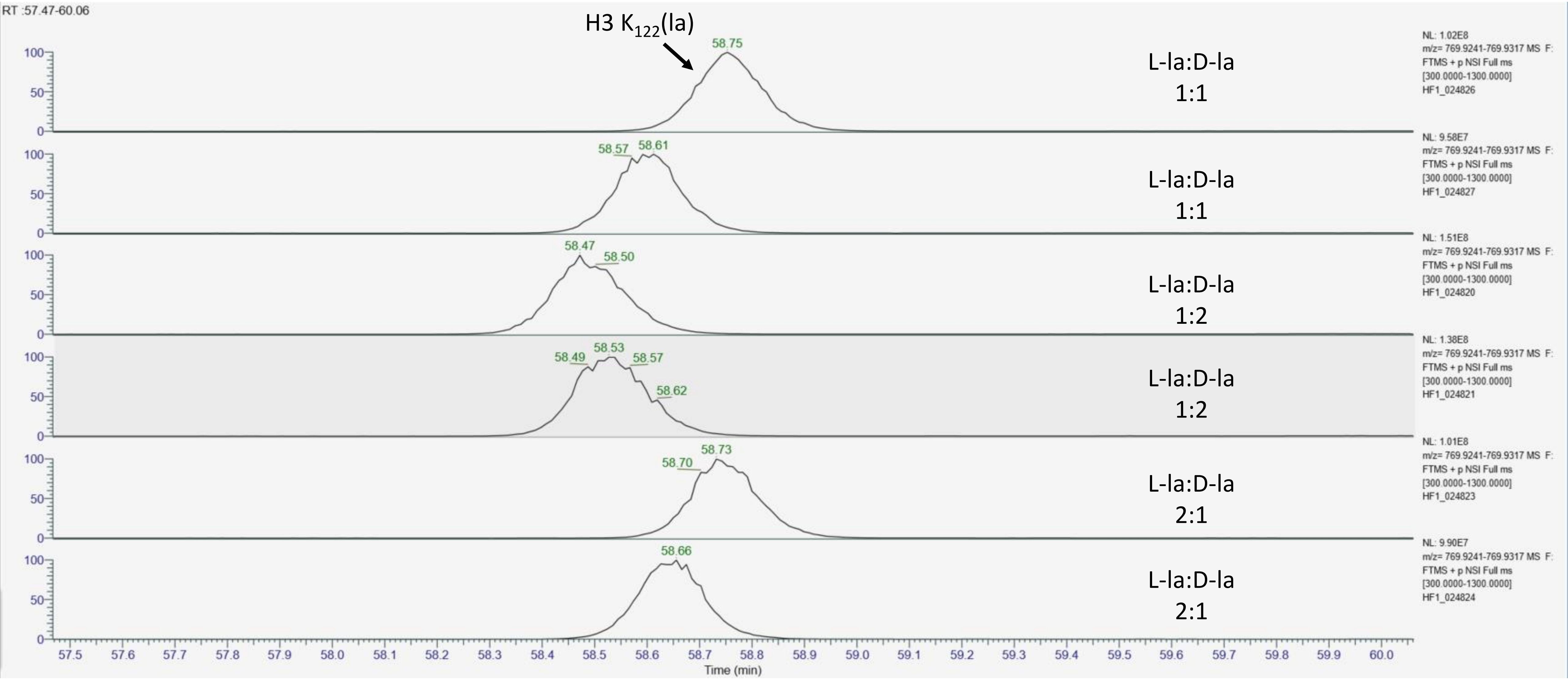

### MS/MS spectra acquired at apex of each chromatographic peak for lactylated H3 VTIM(ox)$P_{122}$DIQLAR in three injections

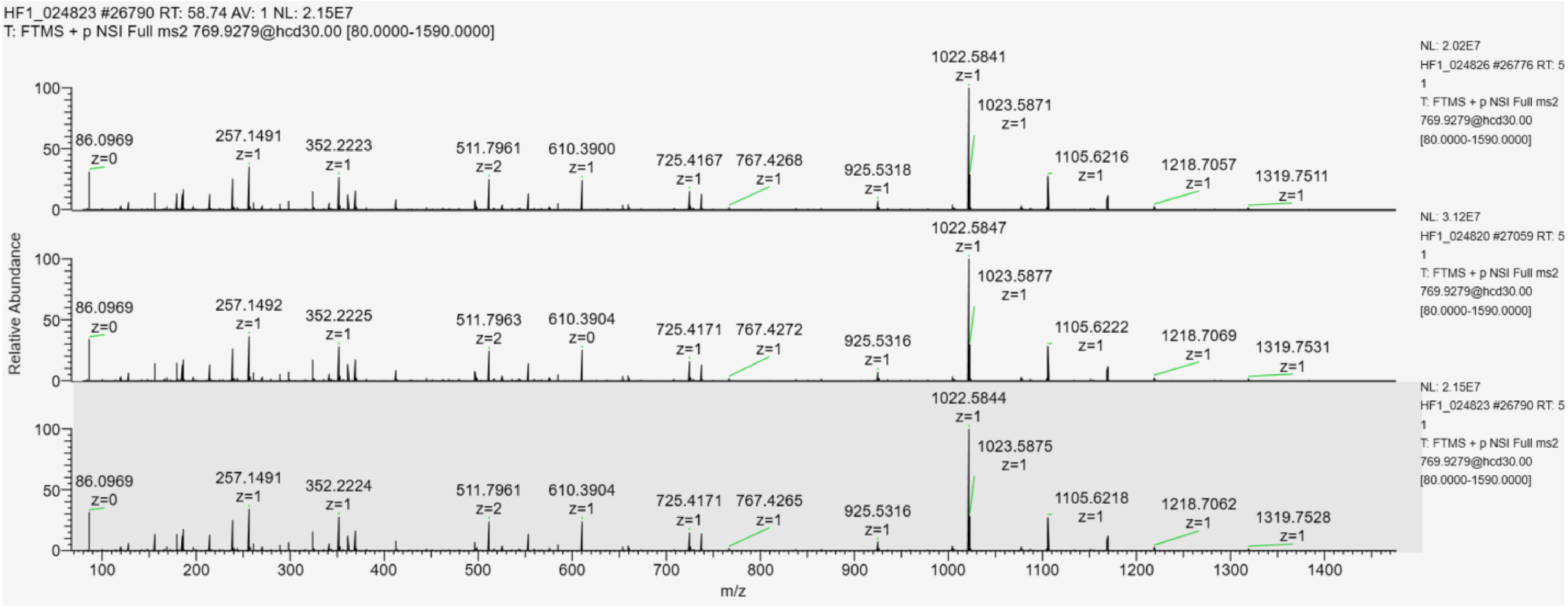


L-la:D-la 1:1

L-la:D-la 1:2

L-la:D-la 2:1

### Extracted ion chromatogram of lactylated H4 $GK_5GGK_8GLGK_{12}GGAK_{16}R$ (m/z= 788.9559,2+)

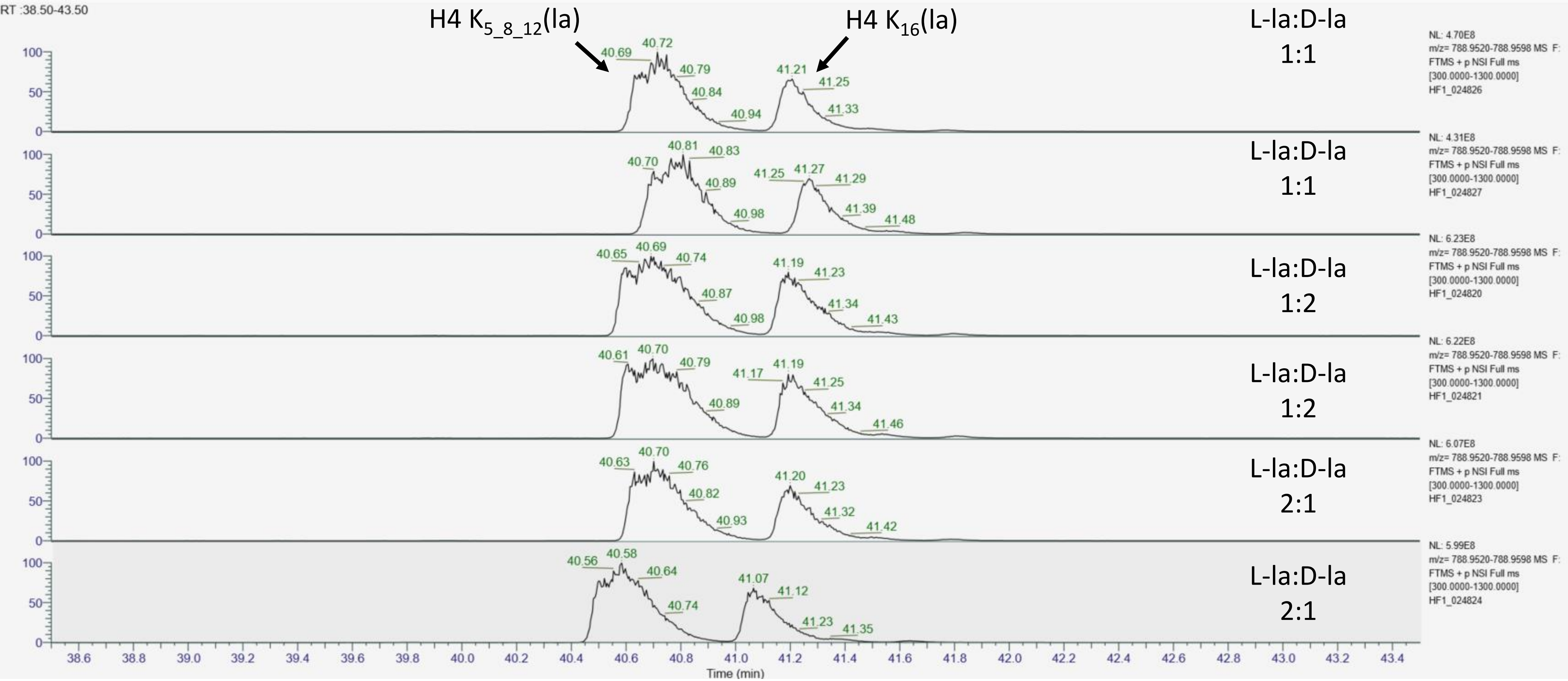

### MS/MS spectra acquired at the apex of each chromatographic peak for H4 GK$_5$(pr)GGK$_8$(pr)GLGK$_{12}$(la)GGAK$_{16}$(pr)R in three injections

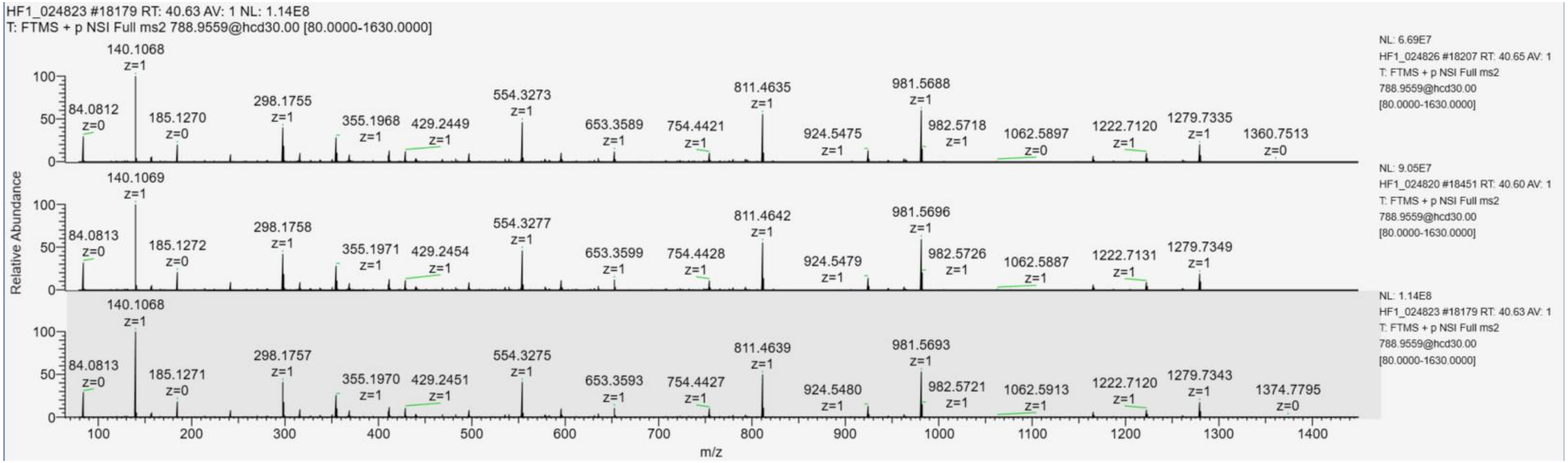


L-la:D-la 1:1

L-la:D-la 1:2

L-la:D-la 2:1

### MS/MS spectra acquired at the apex of each chromatographic peak for H4 $GK_5$(la)$GGK_8$(pr)$GLGK_{12}$(pr)$GGAK_{16}$(pr)R and $GK_5$(pr)$GGK_8$(la)$GLGK_{12}$(pr)$GGAK_{16}$(pr)R

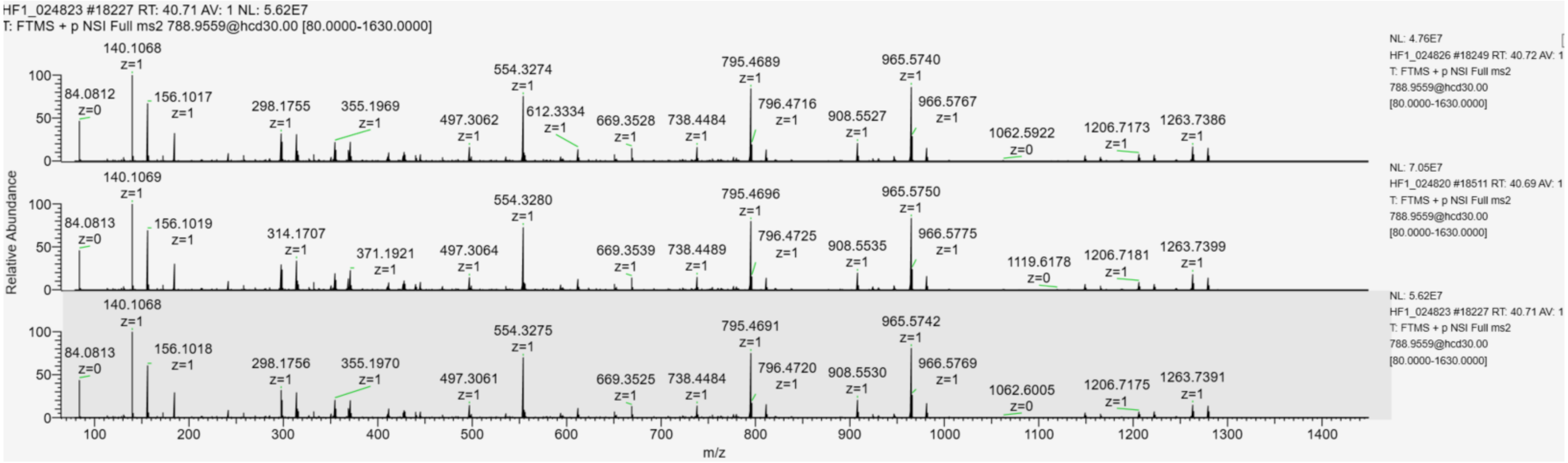


L-la:D-la
1:1

L-la:D-la
1:2

L-la:D-la
2:1

### MS/MS spectra acquired at the apex of each chromatographic peak for H4 GK$_5$(pr)GGK$_8$(pr)GLGK$_{12}$(pr)GGAK$_{16}$(la)R

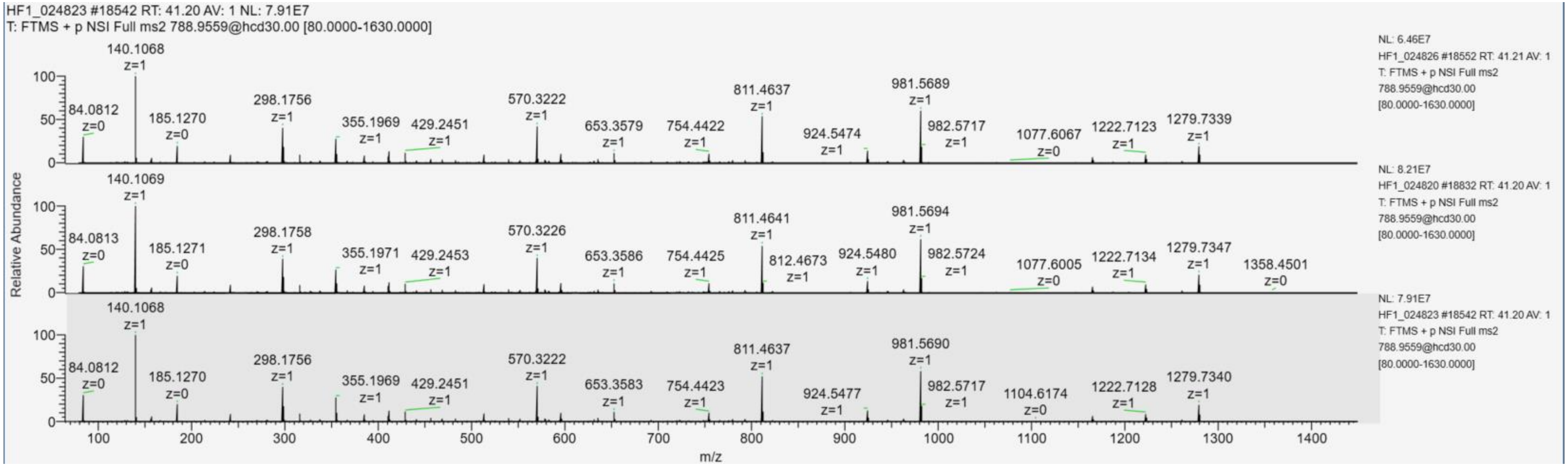


L-la:D-la
1:1

L-la:D-la
1:2

L-la:D-la
2:1

**Extracted ion chromatogram of lactylated H4 DNIQGITK$_{31}$PAIR (m/z= 732.4082,2+)**

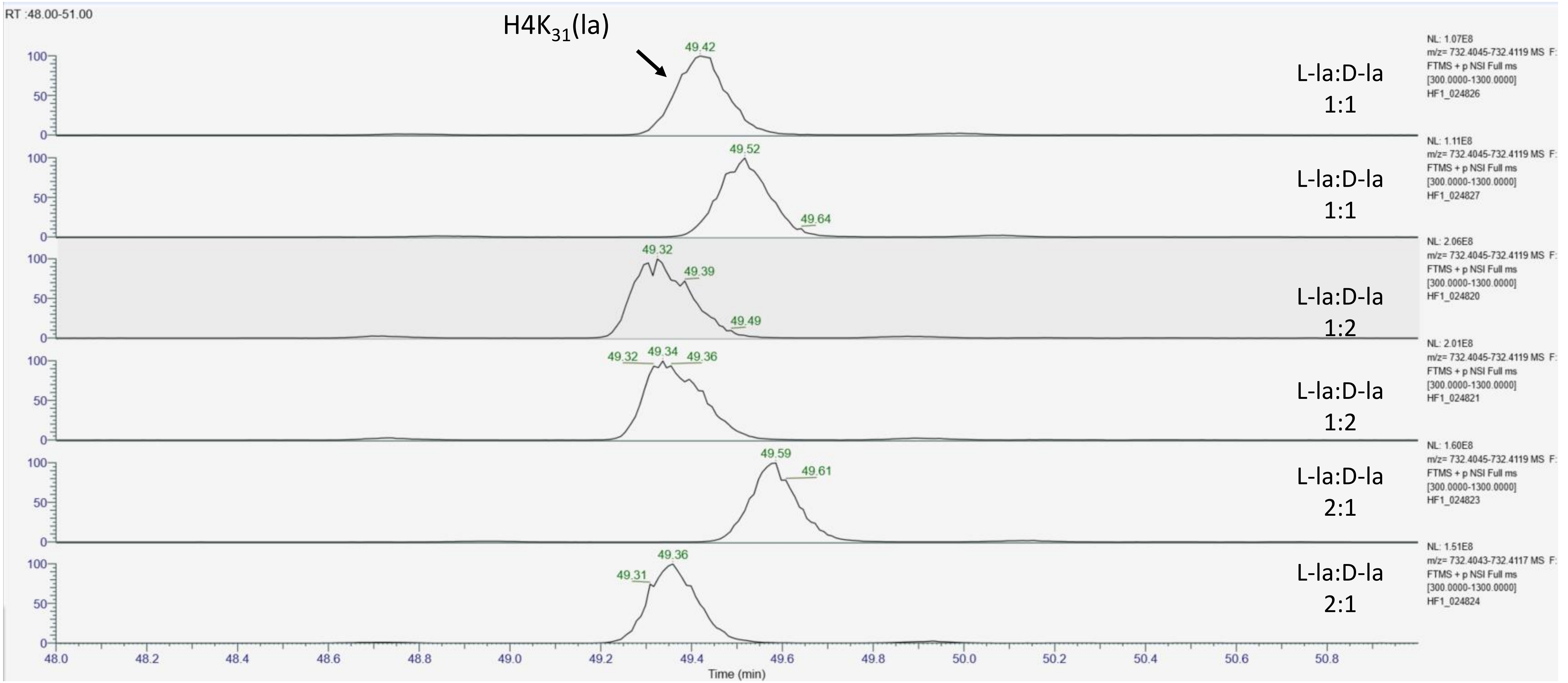

### MS/MS spectra acquired at apex of each chromatographic peak for H4 DNIQGITK$_{31}$(la)R in three injections

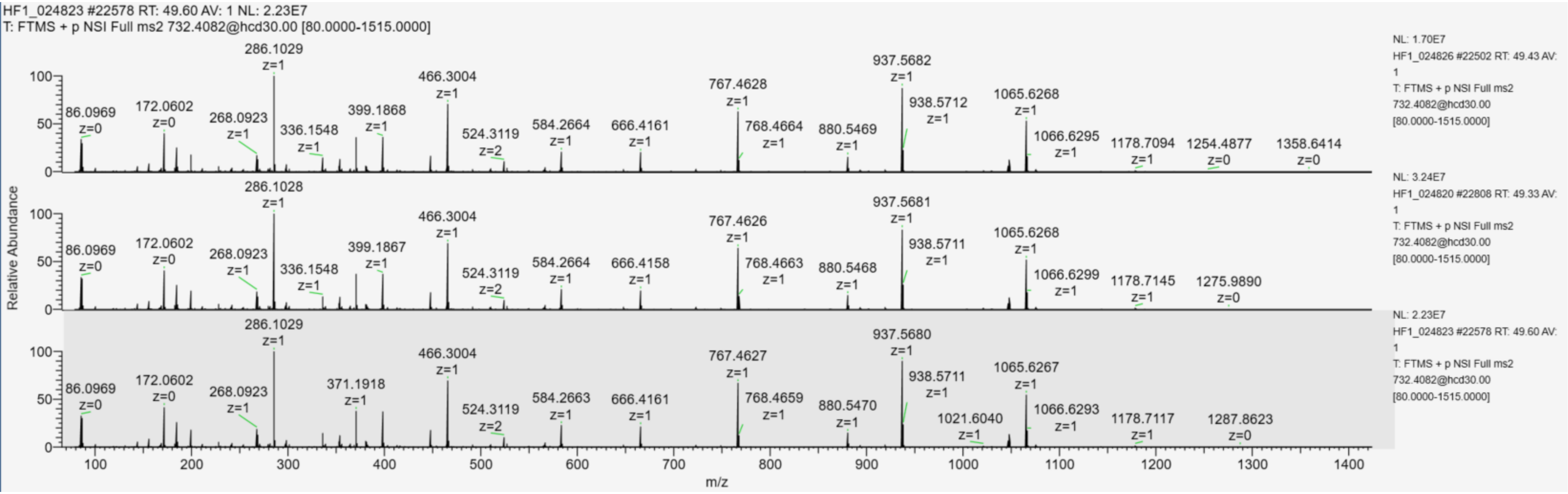


L-la:D-la 1:1

L-la:D-la 1:2

L-la:D-la 2:1

**Extracted ion chromatogram of L- and D-lactylated H4 DAVTYTEHAK$_{77}$R (m/z=714.8533,2+)**

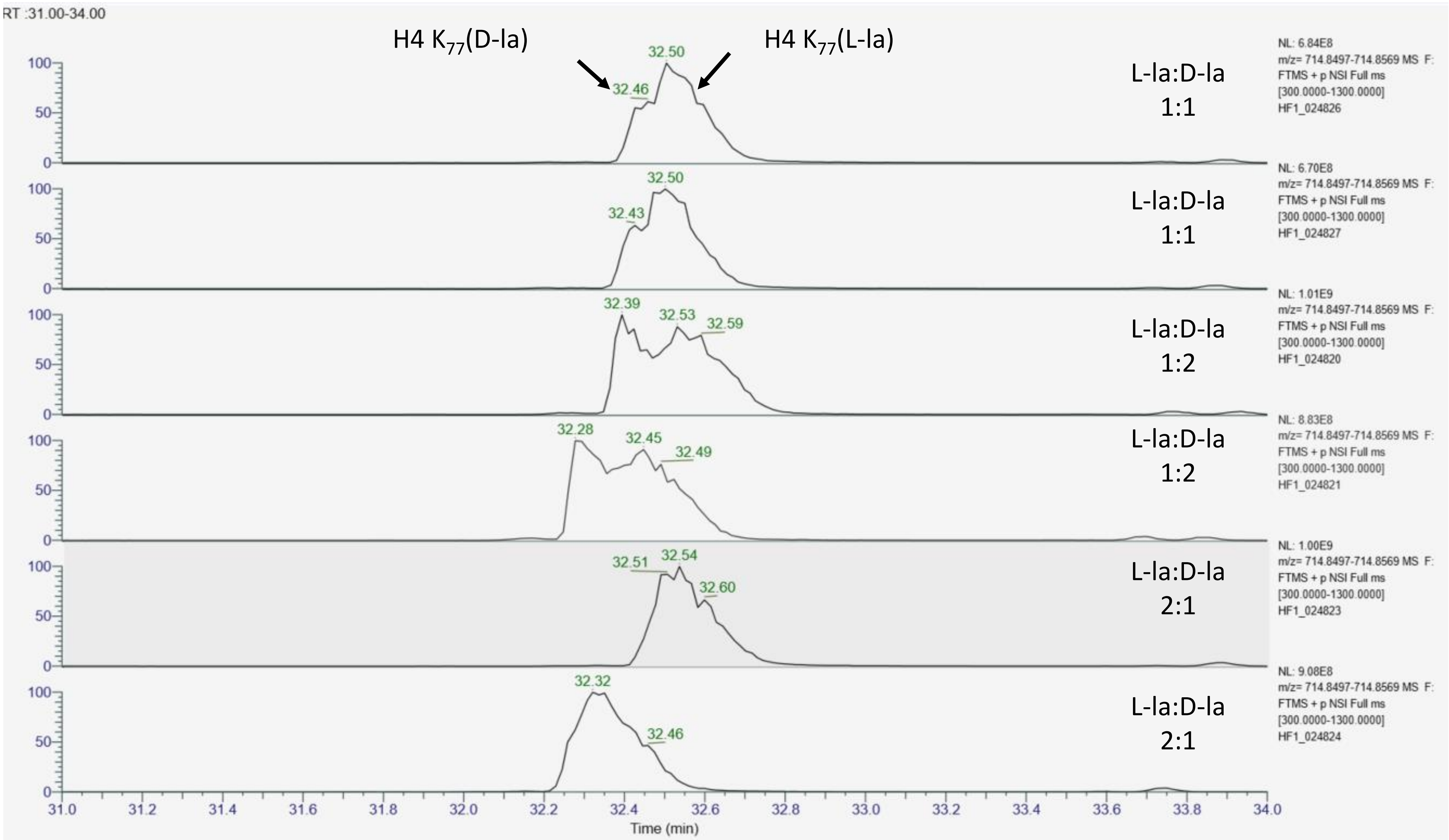

#### MS/MS spectra acquired at the apex of each chromatographic peak for L- and D-lactylated H4 DAVTYTEHAK$_{77}$R

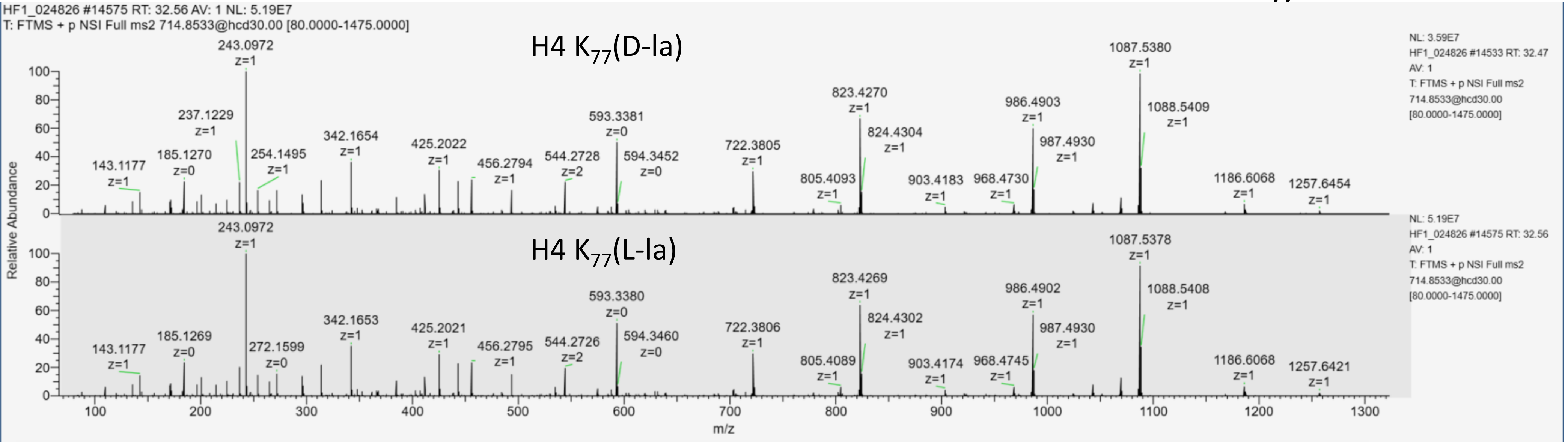


L-la:D-la
1:1

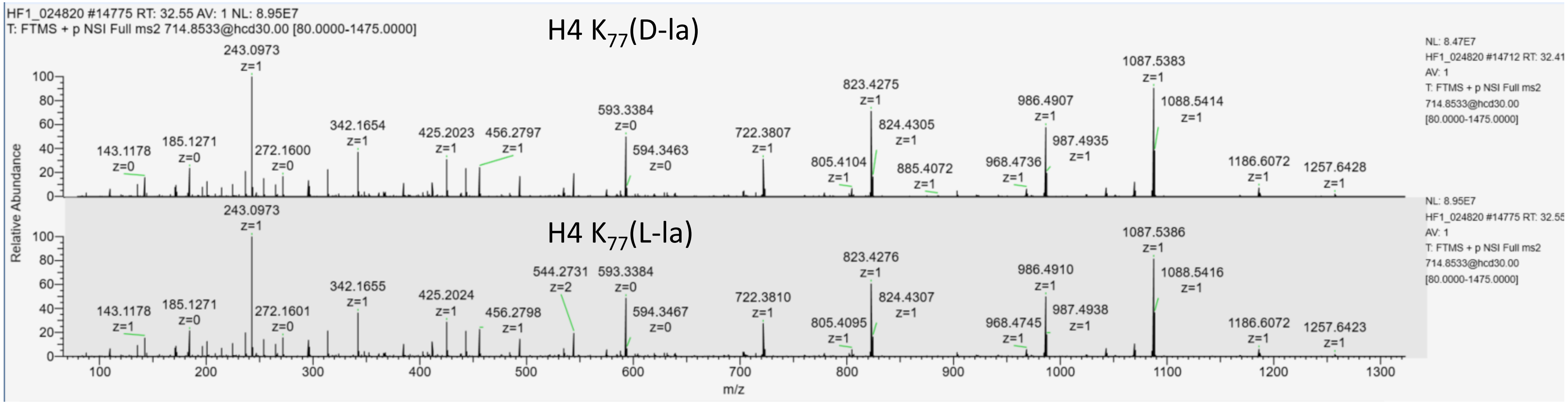


L-la:D-la
1:2

### MS/MS spectra acquired at apex of each chromatographic peak for L- and D-lactylated H4 DAVTYTEHAK$_{77}$R

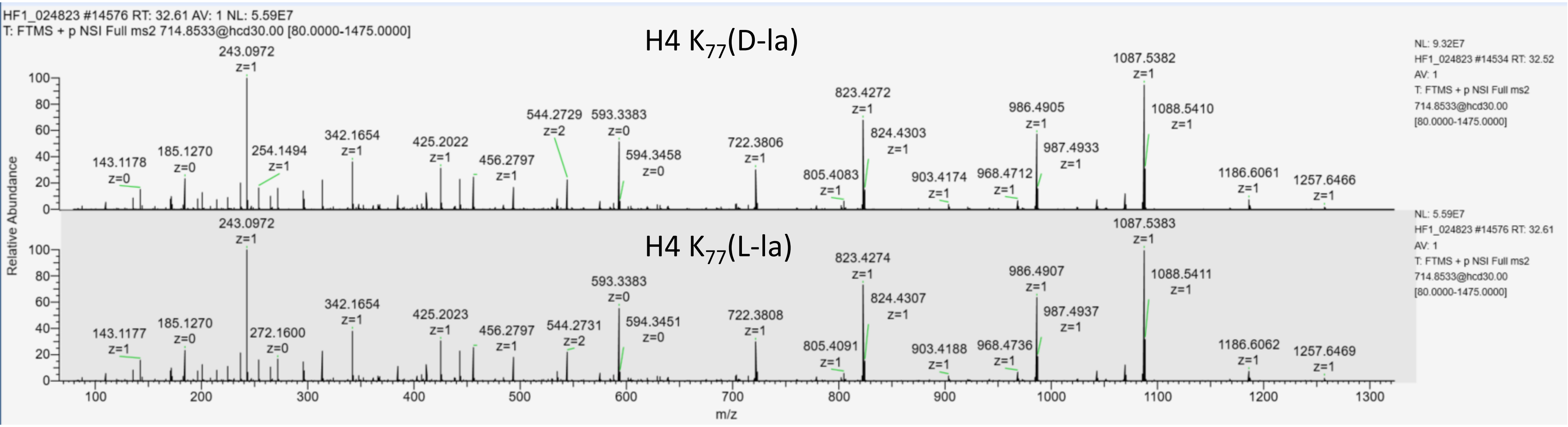


L-la:D-la
2:1

#### Extracted ion chromatogram of lactylated H4 $K_{79}TVTAMDVVYALK_{91}R$ (m/z= 894.9938 ,2+)

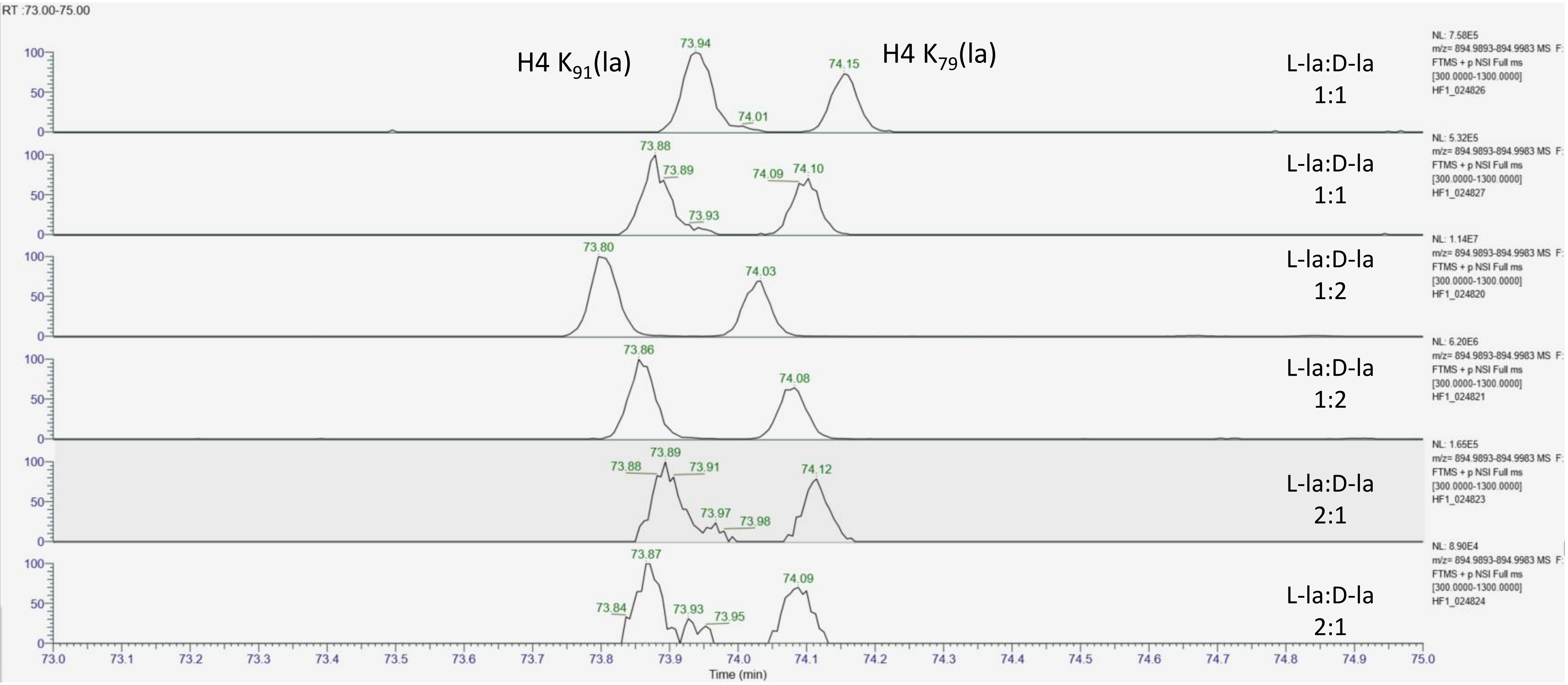

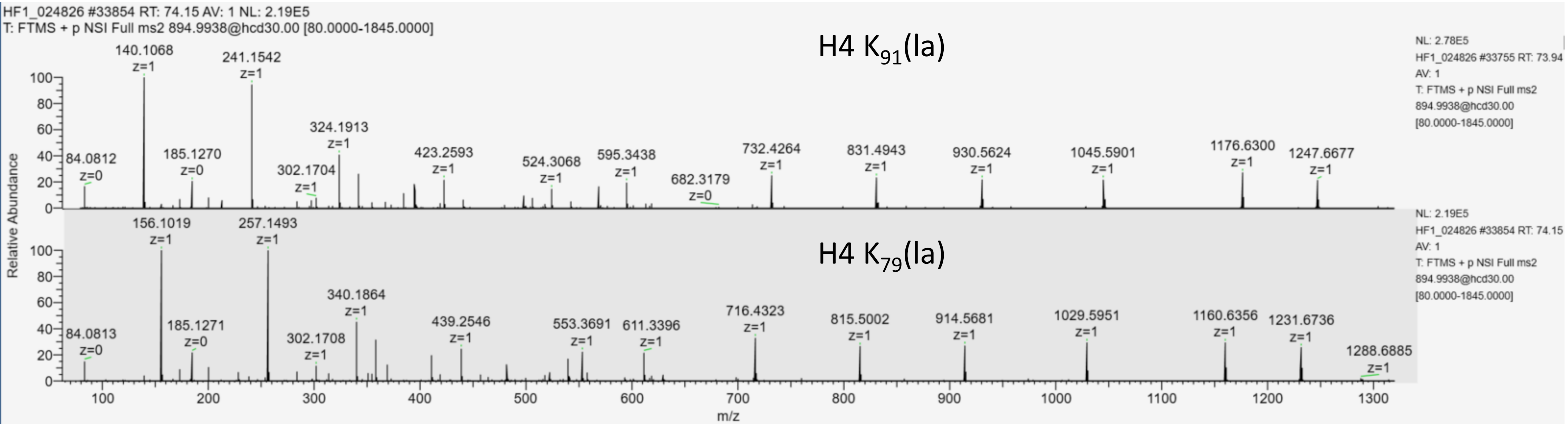


L-la:D-la
1:1

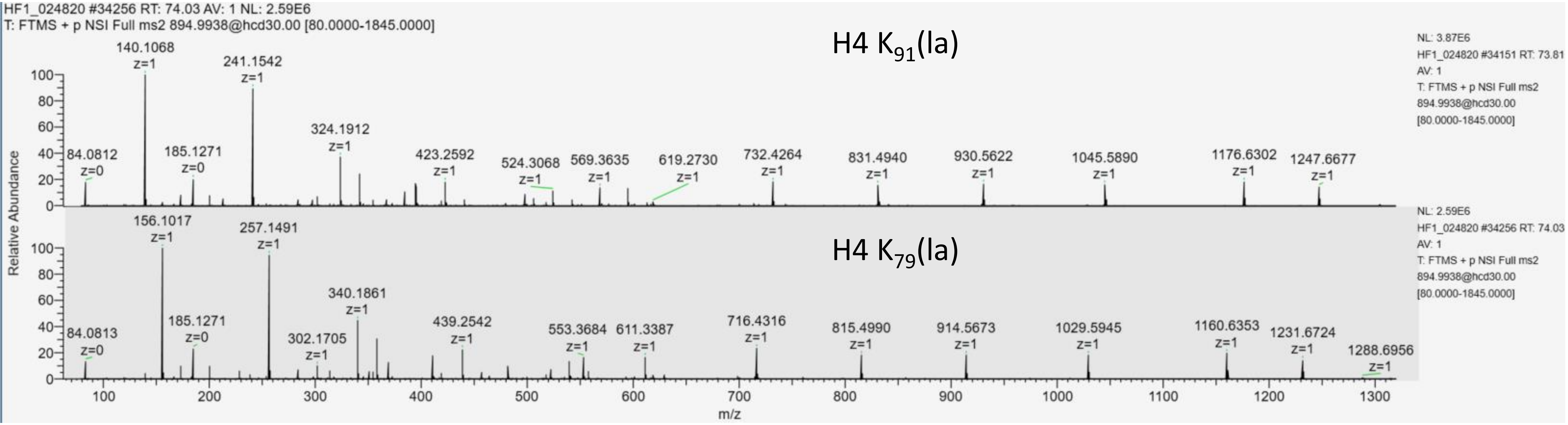


L-la:D-la
1:2

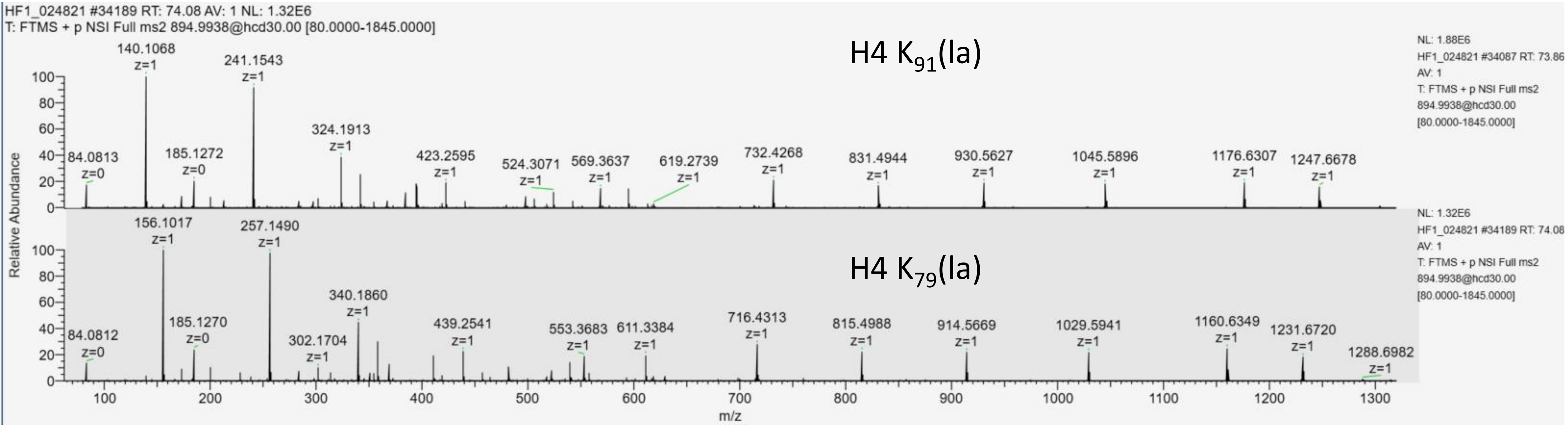
HF1_024821 #34189 RT: 74.08 AV: 1 NL: 1.32E6
T: FTMS + p NSI Full ms2 894.9938@hcd30.00 [80.0000-1845.0000]
H4 $K_{91}$(la)
H4 $K_{79}$(la)
NL: 1.88E6
HF1_024821 #34087 RT: 73.86
AV: 1
T: FTMS + p NSI Full ms2
894.9938@hcd30.00
[80.0000-1845.0000]
NL: 1.32E6
HF1_024821 #34189 RT: 74.08
AV: 1
T: FTMS + p NSI Full ms2
894.9938@hcd30.00
[80.0000-1845.0000]
Relative Abundance
m/z

L-la:D-la
2:1

### Extracted ion chromatogram of lactylated H4 $K_{79}$TVTAM(ox)DVVYALK$_{91}$R (902.9918,2+)

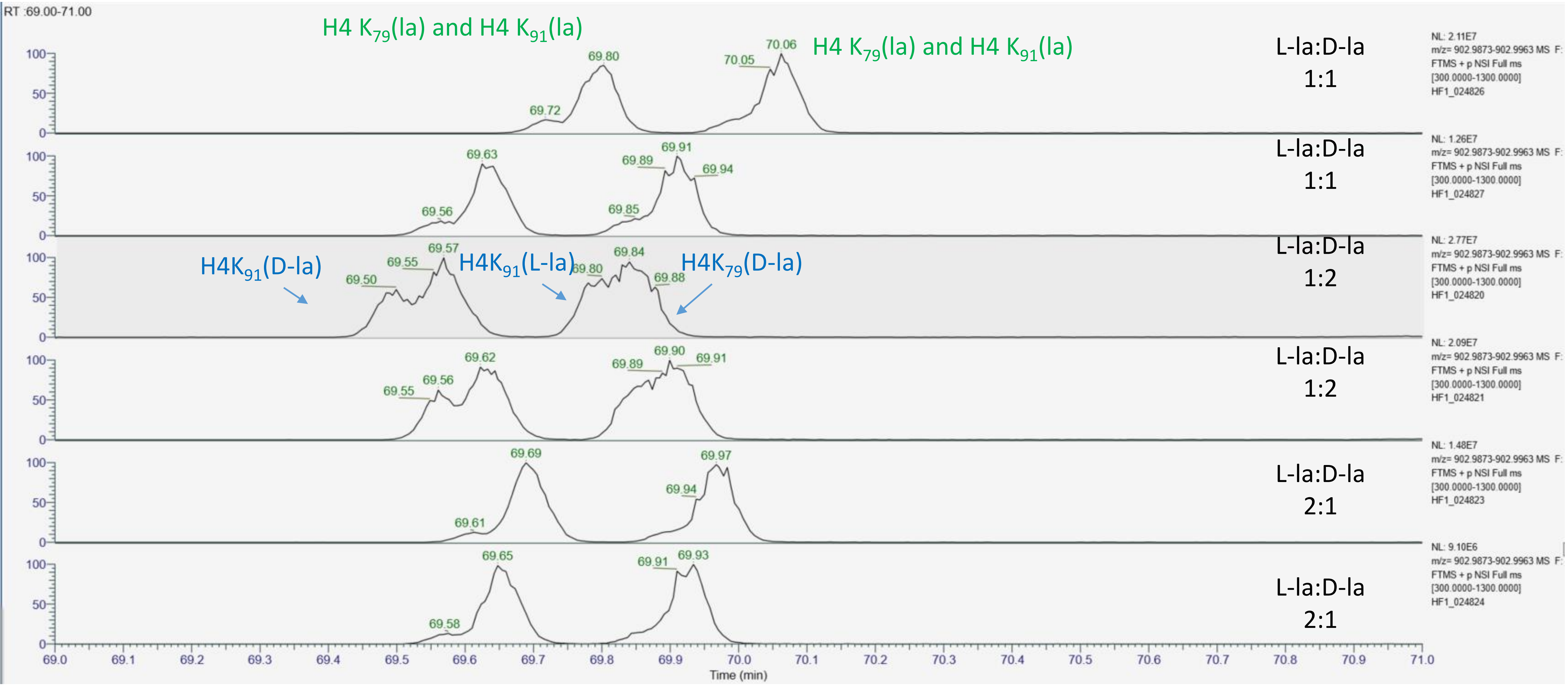

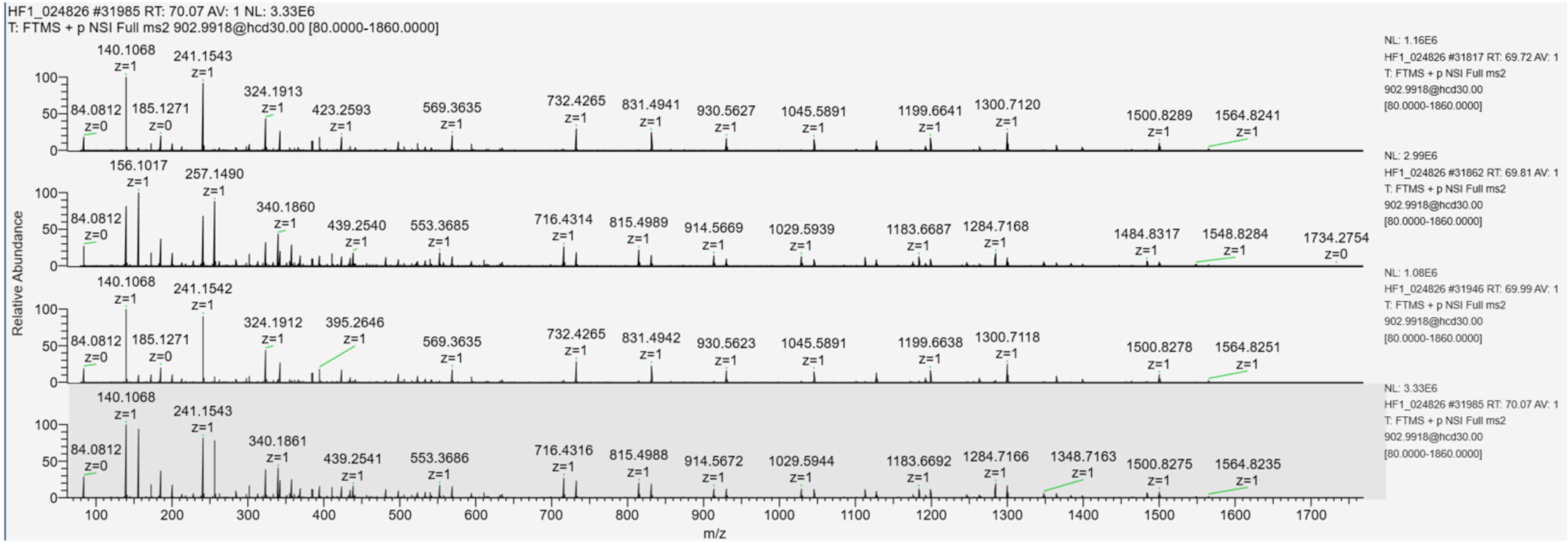


L-la:D-la
1:1

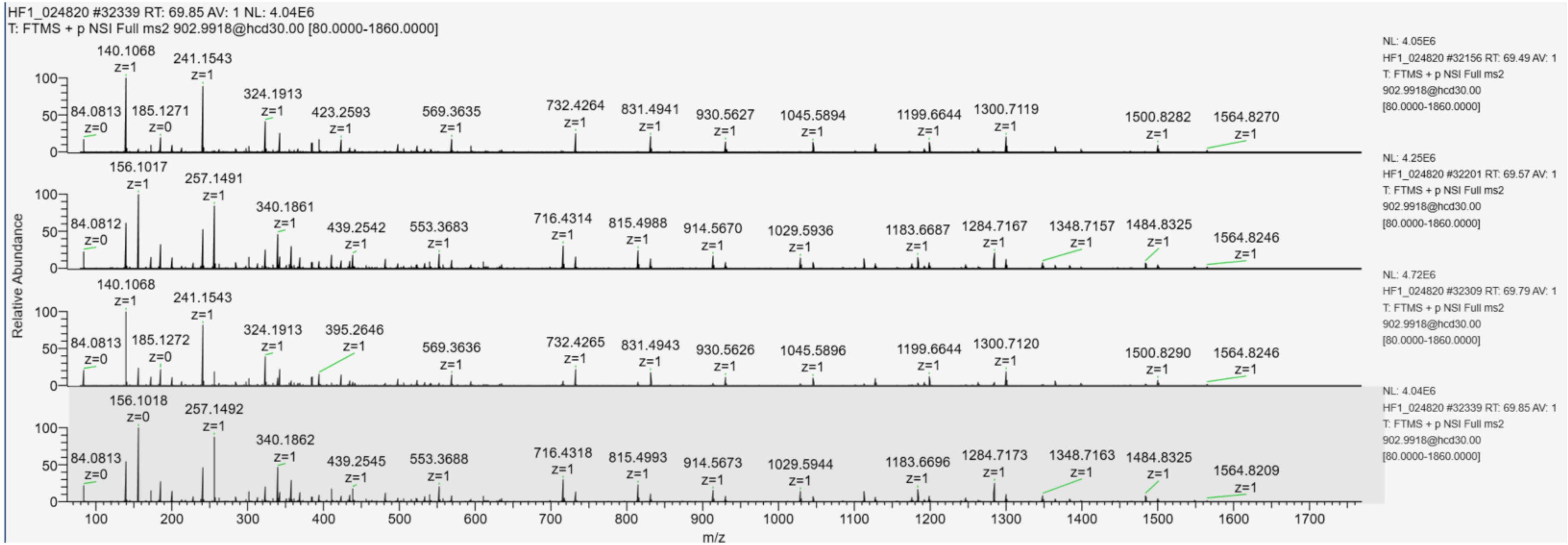


L-la:D-la
1:2

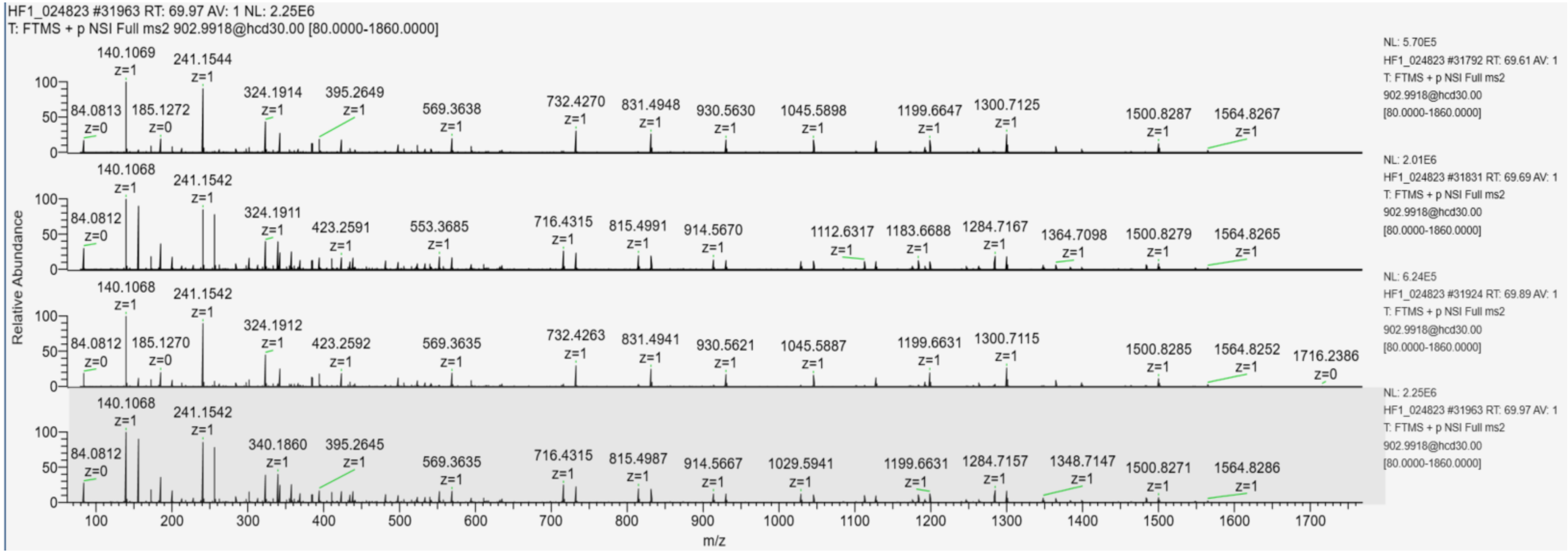
HF1_024823 #31963 RT: 69.97 AV: 1 NL: 2.25E6
T: FTMS + p NSI Full ms2 902.9918@hcd30.00 [80.0000-1860.0000]
Relative Abundance
m/z
NL: 5.70E5
HF1_024823 #31792 RT: 69.61 AV: 1
T: FTMS + p NSI Full ms2
902.9918@hcd30.00
[80.0000-1860.0000]
NL: 2.01E6
HF1_024823 #31831 RT: 69.69 AV: 1
T: FTMS + p NSI Full ms2
902.9918@hcd30.00
[80.0000-1860.0000]
NL: 6.24E5
HF1_024823 #31924 RT: 69.89 AV: 1
T: FTMS + p NSI Full ms2
902.9918@hcd30.00
[80.0000-1860.0000]
NL: 2.25E6
HF1_024823 #31963 RT: 69.97 AV: 1
T: FTMS + p NSI Full ms2
902.9918@hcd30.00
[80.0000-1860.0000]

L-la:D-la
2:1

**Section 4:** Statistical assessment of the abundance difference between L- and D-lactylated peptides that are chromatographically separated.

Two approaches were considered to normalize the MS signals detected for lactylated peptides in the analyses acquired on the four biological replicates of testis histones (Suppl. table 8).

For each variant and each modified lysine site, we compared:

- The abundance of peptides carrying L-La and D-La blocked by biological replicate (results on the left);
- The abundance of peptides normalized to the sum of abundances of lactylated peptides in each run (results on the right).

The results on the left take into account the effect of biological replicate (assuming that all variability comes from biology). The column on the right normalizes by the injected quantity (assuming that all variability comes from analysis). Reality probably lies in-between.

The results of statistical tests assessing whether L- and D-lactyl are of different abundance on the considered lysine residues are indicated at the bottom of the columns; they appear in red when below 0.05.

# H3K18

#### Oneway Analysis of Data By Modif Variant=H3, Site 2=K18

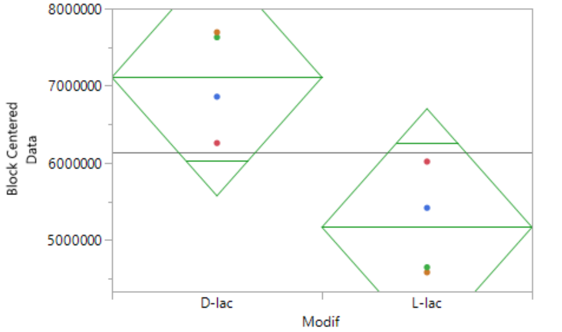


##### Oneway Anova

###### Summary of Fit

| | |
|---|---|
| Rsquare | 0.955218 |
| Adj Rsquare | 0.895509 |
| Root Mean Square Error | 965269.8 |
| Mean of Response | 6138750 |
| Observations (or Sum Wgts) | 8 |

###### Pooled t Test

L-lac-D-lac

Assuming equal variances

| | | | |
|---|---|---|---|
| Difference | -1942500 | t Ratio | -2.84595 |
| Std Err Dif | 682549 | DF | 3 |
| Upper CL Dif | 229675 | Prob > \|t\| | 0.0653 |
| Lower CL Dif | -4114675 | Prob > t | 0.9673 |
| Confidence | 0.95 | Prob < t | 0.0327* |

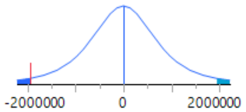


###### Analysis of Variance

| Source | DF | Sum of Squares | Mean Square | F Ratio | Prob > F |
|---|---|---|---|---|---|
| Modif | 1 | 7.5466e+12 | 7.547e+12 | 8.0994 | 0.0653 |
| Label | 3 | 5.2077e+13 | 1.736e+13 | 18.6307 | 0.0192* |
| Error | 3 | 2.7952e+12 | 9.317e+11 | | |
| C. Total | 7 | 6.2419e+13 | | | |

###### Means for Oneway Anova

| Level | Number | Mean | Std Error | Lower 95% | Upper 95% |
|---|---|---|---|---|---|
| D-lac | 4 | 7110000 | 482635 | 5574040.3 | 8645959.7 |
| L-lac | 4 | 5167500 | 482635 | 3631540.3 | 6703459.7 |

Std Error uses a pooled estimate of error variance

###### Block Means

| Label | Mean | Number |
|---|---|---|
| Abundance 1 | 2180000 | 2 |
| Abundance 2 | 9310000 | 2 |
| Abundance 3 | 6530000 | 2 |
| Abundance 4 | 6535000 | 2 |

##### Friedman Rank Test

| Level | Count | Score Sum | Expected Score | Score Mean | (Mean-Mean0)/Std0 |
|---|---|---|---|---|---|
| D-lac | 4 | 8.000 | 6.000 | 2.00000 | 1.871 |
| L-lac | 4 | 4.000 | 6.000 | 1.00000 | -1.871 |

###### 2 - Sample Test, Normal Approximation

| S | Z | Prob>\|Z\| |
|---|---|---|
| 4 | -1.87083 | 0.0614 |

###### 1-Way Test, ChiSquare Approximation

| ChiSquare | DF | Prob>ChiSq |
|---|---|---|
| 4.0000 | 1 | 0.0455* |

Small sample sizes. Refer to statistical tables for tests, rather than large-sample approximations.

Block Label

#### Oneway Analysis of Data By Modif Variant=H3, Lys=K18

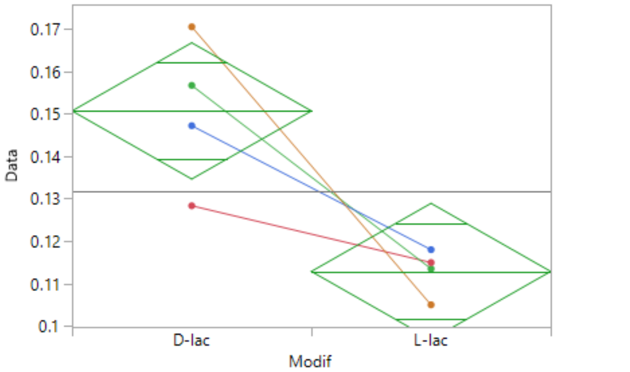


##### Oneway Anova

###### Summary of Fit

| | |
|---|---|
| Rsquare | 0.734724 |
| Adj Rsquare | 0.690512 |
| Root Mean Square Error | 0.013133 |
| Mean of Response | 0.131771 |
| Observations (or Sum Wgts) | 8 |

###### Pooled t Test

L-lac-D-lac

Assuming equal variances

| | | | |
|---|---|---|---|
| Difference | -0.03786 | t Ratio | -4.07652 |
| Std Err Dif | 0.00929 | DF | 6 |
| Upper CL Dif | -0.01513 | Prob > \|t\| | 0.0065* |
| Lower CL Dif | -0.06058 | Prob > t | 0.9967 |
| Confidence | 0.95 | Prob < t | 0.0033* |

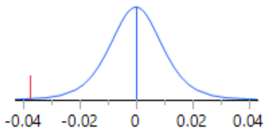


###### Analysis of Variance

| Source | DF | Sum of Squares | Mean Square | F Ratio | Prob > F |
|---|---|---|---|---|---|
| Modif | 1 | 0.00286640 | 0.002866 | 16.6180 | 0.0065* |
| Error | 6 | 0.00103493 | 0.000172 | | |
| C. Total | 7 | 0.00390133 | | | |

###### Means for Oneway Anova

| Level | Number | Mean | Std Error | Lower 95% | Upper 95% |
|---|---|---|---|---|---|
| D-lac | 4 | 0.150700 | 0.00657 | 0.13463 | 0.16677 |
| L-lac | 4 | 0.112842 | 0.00657 | 0.09677 | 0.12891 |

Std Error uses a pooled estimate of error variance

##### Wilcoxon / Kruskal-Wallis Tests (Rank Sums)

| Level | Count | Score Sum | Expected Score | Score Mean | (Mean-Mean0)/Std0 |
|---|---|---|---|---|---|
| D-lac | 4 | 26.000 | 18.000 | 6.50000 | 2.165 |
| L-lac | 4 | 10.000 | 18.000 | 2.50000 | -2.165 |

###### Wilcoxon Two-Sample Test, Normal Approximation

| S | Z | Prob>\|Z\| |
|---|---|---|
| 10 | -2.16506 | 0.0304* |

###### Kruskal-Wallis Test, ChiSquare Approximation

| ChiSquare | DF | Prob>ChiSq |
|---|---|---|
| 5.3333 | 1 | 0.0209* |

Small sample sizes. Refer to statistical tables for tests, rather than large-sample approximations.

H3K23

#### Oneway Analysis of Data By Modif Variant=H3, Site 2=K23

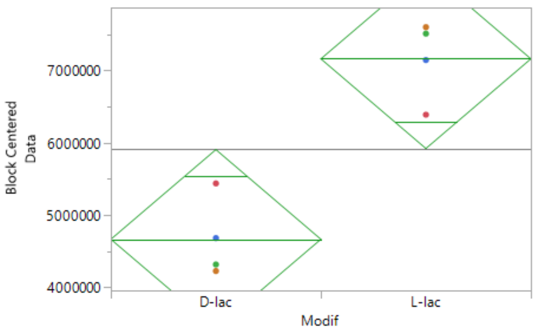


##### Oneway Anova

###### Summary of Fit

| | |
|---|---|
| Rsquare | 0.970035 |
| Adj Rsquare | 0.930082 |
| Root Mean Square Error | 778553.7 |
| Mean of Response | 5913750 |
| Observations (or Sum Wgts) | 8 |

###### Pooled t Test

L-lac-D-lac

Assuming equal variances

| | | | |
|---|---|---|---|
| Difference | 2492500 | t Ratio | 4.527533 |
| Std Err Dif | 550521 | DF | 3 |
| Upper CL Dif | 4244502 | Prob > \|t\| | 0.0202* |
| Lower CL Dif | 740498 | Prob > t | 0.0101* |
| Confidence | 0.95 | Prob < t | 0.9899 |

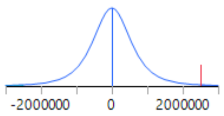


###### Analysis of Variance

| Source | DF | Sum of Squares | Mean Square | F Ratio | Prob > F |
|---|---|---|---|---|---|
| Modif | 1 | 1.2425e+13 | 1.243e+13 | 20.4986 | 0.0202* |
| Label | 3 | 4.6442e+13 | 1.548e+13 | 25.5394 | 0.0123* |
| Error | 3 | 1.8184e+12 | 6.061e+11 | | |
| C. Total | 7 | 6.0685e+13 | | | |

###### Means for Oneway Anova

| Level | Number | Mean | Std Error | Lower 95% | Upper 95% |
|---|---|---|---|---|---|
| D-lac | 4 | 4667500 | 389277 | 3428647.4 | 5906352.6 |
| L-lac | 4 | 7160000 | 389277 | 5921147.4 | 8398852.6 |

Std Error uses a pooled estimate of error variance

###### Block Means

| Label | Mean | Number |
|---|---|---|
| Abundance 1 | 2315000 | 2 |
| Abundance 2 | 9105000 | 2 |
| Abundance 3 | 6080000 | 2 |
| Abundance 4 | 6155000 | 2 |

##### Friedman Rank Test

| Level | Count | Score Sum | Expected Score | Score Mean | (Mean-Mean0)/Std0 |
|---|---|---|---|---|---|
| D-lac | 4 | 4.000 | 6.000 | 1.00000 | -1.871 |
| L-lac | 4 | 8.000 | 6.000 | 2.00000 | 1.871 |

###### 2 - Sample Test, Normal Approximation

| S | Z | Prob>\|Z\| |
|---|---|---|
| 8 | 1.87083 | 0.0614 |

###### 1-Way Test, ChiSquare Approximation

| ChiSquare | DF | Prob>ChiSq |
|---|---|---|
| 4.0000 | 1 | 0.0455* |

Small sample sizes. Refer to statistical tables for tests, rather than large-sample approximations.

Block Label

#### Oneway Analysis of Data By Modif Variant=H3, Lys=K23

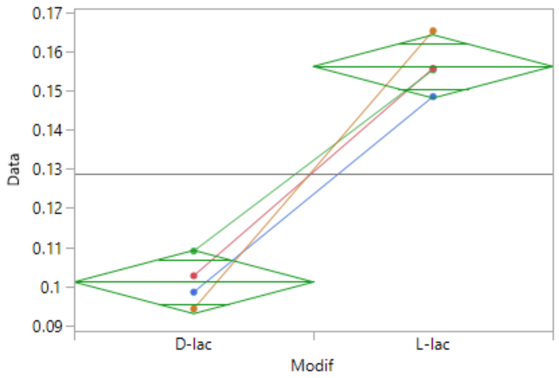


##### Oneway Anova

###### Summary of Fit

| | |
|---|---|
| Rsquare | 0.958572 |
| Adj Rsquare | 0.951667 |
| Root Mean Square Error | 0.00661 |
| Mean of Response | 0.128624 |
| Observations (or Sum Wgts) | 8 |

###### Pooled t Test

L-lac-D-lac

Assuming equal variances

| | | | |
|---|---|---|---|
| Difference | 0.055072 | t Ratio | 11.78254 |
| Std Err Dif | 0.004674 | DF | 6 |
| Upper CL Dif | 0.066508 | Prob > \|t\| | <.0001* |
| Lower CL Dif | 0.043635 | Prob > t | <.0001* |
| Confidence | 0.95 | Prob < t | 1.0000 |

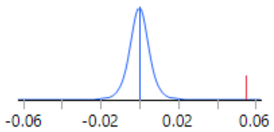


###### Analysis of Variance

| Source | DF | Sum of Squares | Mean Square | F Ratio | Prob > F |
|---|---|---|---|---|---|
| Modif | 1 | 0.00606576 | 0.006066 | 138.8284 | <.0001* |
| Error | 6 | 0.00026216 | 0.000044 | | |
| C. Total | 7 | 0.00632792 | | | |

###### Means for Oneway Anova

| Level | Number | Mean | Std Error | Lower 95% | Upper 95% |
|---|---|---|---|---|---|
| D-lac | 4 | 0.101088 | 0.00331 | 0.09300 | 0.10917 |
| L-lac | 4 | 0.156159 | 0.00331 | 0.14807 | 0.16425 |

Std Error uses a pooled estimate of error variance

##### Wilcoxon / Kruskal-Wallis Tests (Rank Sums)

| Level | Count | Score Sum | Expected Score | Score Mean | (Mean-Mean0)/Std0 |
|---|---|---|---|---|---|
| D-lac | 4 | 10.000 | 18.000 | 2.50000 | -2.165 |
| L-lac | 4 | 26.000 | 18.000 | 6.50000 | 2.165 |

###### Wilcoxon Two-Sample Test, Normal Approximation

| S | Z | Prob>\|Z\| |
|---|---|---|
| 26 | 2.16506 | 0.0304* |

###### Kruskal-Wallis Test, ChiSquare Approximation

| ChiSquare | DF | Prob>ChiSq |
|---|---|---|
| 5.3333 | 1 | 0.0209* |

Small sample sizes. Refer to statistical tables for tests, rather than large-sample approximations.

# H3.tK18

#### Oneway Analysis of Data By Modif Variant=H3.t, Site 2=K18

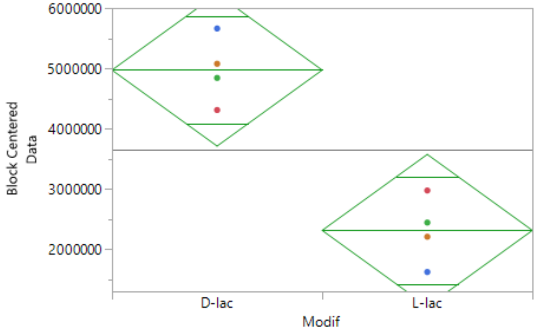


##### Oneway Anova

###### Summary of Fit

| | |
|---|---|
| Rsquare | 0.943242 |
| Adj Rsquare | 0.867565 |
| Root Mean Square Error | 792855.5 |
| Mean of Response | 3641875 |
| Observations (or Sum Wgts) | 8 |

###### Pooled t Test

L-lac-D-lac

Assuming equal variances

| | | | |
|---|---|---|---|
| Difference | -2661250 | t Ratio | -4.74686 |
| Std Err Dif | 560633 | DF | 3 |
| Upper CL Dif | -877064 | Prob > \|t\| | 0.0177* |
| Lower CL Dif | -4445436 | Prob > t | 0.9911 |
| Confidence | 0.95 | Prob < t | 0.0089* |

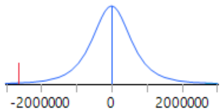


###### Analysis of Variance

| Source | DF | Sum of Squares | Mean Square | F Ratio | Prob > F |
|---|---|---|---|---|---|
| Modif | 1 | 1.4165e+13 | 1.416e+13 | 22.5327 | 0.0177* |
| Label | 3 | 1.7176e+13 | 5.725e+12 | 9.1079 | 0.0512 |
| Error | 3 | 1.8859e+12 | 6.286e+11 | | |
| C. Total | 7 | 3.3226e+13 | | | |

###### Means for Oneway Anova

| Level | Number | Mean | Std Error | Lower 95% | Upper 95% |
|---|---|---|---|---|---|
| D-lac | 4 | 4972500 | 396428 | 3710890 | 6234110 |
| L-lac | 4 | 2311250 | 396428 | 1049640 | 3572860 |

Std Error uses a pooled estimate of error variance

###### Block Means

| Label | Mean | Number |
|---|---|---|
| Abundance 1 | 1492500 | 2 |
| Abundance 2 | 5620000 | 2 |
| Abundance 3 | 3870000 | 2 |
| Abundance 4 | 3585000 | 2 |

##### Friedman Rank Test

| Level | Count | Score Sum | Expected Score | Score Mean | (Mean-Mean0)/Std0 |
|---|---|---|---|---|---|
| D-lac | 4 | 8.000 | 6.000 | 2.00000 | 1.871 |
| L-lac | 4 | 4.000 | 6.000 | 1.00000 | -1.871 |

###### 2 - Sample Test, Normal Approximation

| S | Z | Prob>\|Z\| |
|---|---|---|
| 4 | -1.87083 | 0.0614 |

###### 1-Way Test, ChiSquare Approximation

| ChiSquare | DF | Prob>ChiSq |
|---|---|---|
| 4.0000 | 1 | 0.0455* |

Small sample sizes. Refer to statistical tables for tests, rather than large-sample approximations.

Block Label

#### Oneway Analysis of Data By Modif Variant=H3.t, Lys=K18

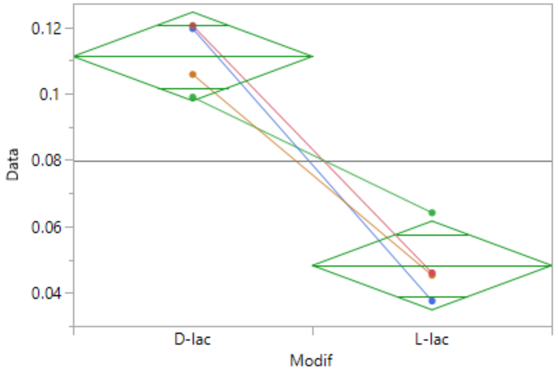


##### Oneway Anova

###### Summary of Fit

| | |
|---|---|
| Rsquare | 0.917146 |
| Adj Rsquare | 0.903337 |
| Root Mean Square Error | 0.010926 |
| Mean of Response | 0.079745 |
| Observations (or Sum Wgts) | 8 |

###### Pooled t Test

L-lac-D-lac

Assuming equal variances

| | | | |
|---|---|---|---|
| Difference | -0.06296 | t Ratio | -8.14964 |
| Std Err Dif | 0.00773 | DF | 6 |
| Upper CL Dif | -0.04406 | Prob > \|t\| | 0.0002* |
| Lower CL Dif | -0.08187 | Prob > t | 0.9999 |
| Confidence | 0.95 | Prob < t | <.0001* |

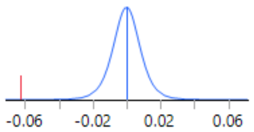


###### Analysis of Variance

| Source | DF | Sum of Squares | Mean Square | F Ratio | Prob > F |
|---|---|---|---|---|---|
| Modif | 1 | 0.00792845 | 0.007928 | 66.4166 | 0.0002* |
| Error | 6 | 0.00071625 | 0.000119 | | |
| C. Total | 7 | 0.00864470 | | | |

###### Means for Oneway Anova

| Level | Number | Mean | Std Error | Lower 95% | Upper 95% |
|---|---|---|---|---|---|
| D-lac | 4 | 0.111226 | 0.00546 | 0.09786 | 0.12459 |
| L-lac | 4 | 0.048264 | 0.00546 | 0.03490 | 0.06163 |

Std Error uses a pooled estimate of error variance

##### Wilcoxon / Kruskal-Wallis Tests (Rank Sums)

| Level | Count | Score Sum | Expected Score | Score Mean | (Mean-Mean0)/Std0 |
|---|---|---|---|---|---|
| D-lac | 4 | 26.000 | 18.000 | 6.50000 | 2.165 |
| L-lac | 4 | 10.000 | 18.000 | 2.50000 | -2.165 |

###### Wilcoxon Two-Sample Test, Normal Approximation

| S | Z | Prob>\|Z\| |
|---|---|---|
| 10 | -2.16506 | 0.0304* |

###### Kruskal-Wallis Test, ChiSquare Approximation

| ChiSquare | DF | Prob>ChiSq |
|---|---|---|
| 5.3333 | 1 | 0.0209* |

Small sample sizes. Refer to statistical tables for tests, rather than large-sample approximations.

H3.tK23

#### Oneway Analysis of Data By Modif Variant=H3.t, Site 2=K23

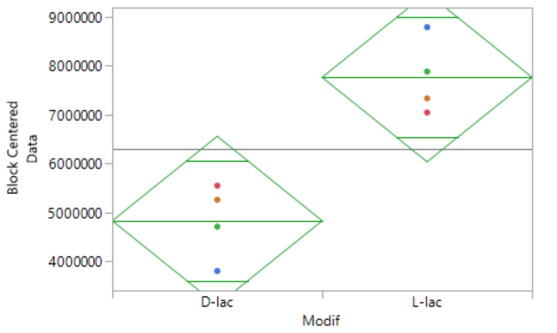


##### Oneway Anova

###### Summary of Fit

| | |
|---|---|
| Rsquare | 0.945756 |
| Adj Rsquare | 0.873431 |
| Root Mean Square Error | 1088914 |
| Mean of Response | 6295000 |
| Observations (or Sum Wgts) | 8 |

###### Pooled t Test

L-lac-D-lac

Assuming equal variances

| | | | |
|---|---|---|---|
| Difference | 2940000 | t Ratio | 3.818289 |
| Std Err Dif | 769978 | DF | 3 |
| Upper CL Dif | 5390415 | Prob > \|t\| | 0.0316* |
| Lower CL Dif | 489585 | Prob > t | 0.0158* |
| Confidence | 0.95 | Prob < t | 0.9842 |

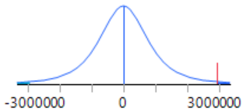


###### Analysis of Variance

| Source | DF | Sum of Squares | Mean Square | F Ratio | Prob > F |
|---|---|---|---|---|---|
| Modif | 1 | 1.7287e+13 | 1.729e+13 | 14.5793 | 0.0316* |
| Label | 3 | 4.4733e+13 | 1.491e+13 | 12.5755 | 0.0332* |
| Error | 3 | 3.5572e+12 | 1.186e+12 | | |
| C. Total | 7 | 6.5578e+13 | | | |

###### Means for Oneway Anova

| Level | Number | Mean | Std Error | Lower 95% | Upper 95% |
|---|---|---|---|---|---|
| D-lac | 4 | 4825000 | 544457 | 3092295.1 | 6557704.9 |
| L-lac | 4 | 7765000 | 544457 | 6032295.1 | 9497704.9 |

Std Error uses a pooled estimate of error variance

###### Block Means

| Label | Mean | Number |
|---|---|---|
| Abundance 1 | 2510000 | 2 |
| Abundance 2 | 9010000 | 2 |
| Abundance 3 | 7050000 | 2 |
| Abundance 4 | 6610000 | 2 |

##### Friedman Rank Test

| Level | Count | Score Sum | Expected Score | Score Mean | (Mean-Mean0)/Std0 |
|---|---|---|---|---|---|
| D-lac | 4 | 4.000 | 6.000 | 1.00000 | -1.871 |
| L-lac | 4 | 8.000 | 6.000 | 2.00000 | 1.871 |

###### 2 - Sample Test, Normal Approximation

| S | Z | Prob>\|Z\| |
|---|---|---|
| 8 | 1.87083 | 0.0614 |

###### 1-Way Test, ChiSquare Approximation

| ChiSquare | DF | Prob>ChiSq |
|---|---|---|
| 4.0000 | 1 | 0.0455* |

Small sample sizes. Refer to statistical tables for tests, rather than large-sample approximations.

Block Label

#### Oneway Analysis of Data By Modif Variant=H3.t, Lys=K23

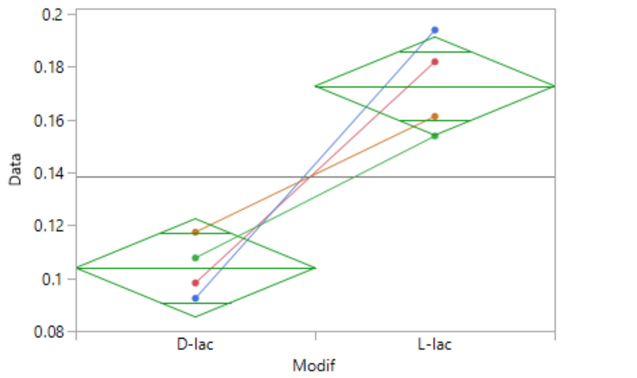


##### Oneway Anova

###### Summary of Fit

| | |
|---|---|
| Rsquare | 0.872447 |
| Adj Rsquare | 0.851189 |
| Root Mean Square Error | 0.015189 |
| Mean of Response | 0.138321 |
| Observations (or Sum Wgts) | 8 |

###### Pooled t Test

L-lac-D-lac

Assuming equal variances

| | | | |
|---|---|---|---|
| Difference | 0.068804 | t Ratio | 6.4062 |
| Std Err Dif | 0.010740 | DF | 6 |
| Upper CL Dif | 0.095084 | Prob > \|t\| | 0.0007* |
| Lower CL Dif | 0.042523 | Prob > t | 0.0003* |
| Confidence | 0.95 | Prob < t | 0.9997 |

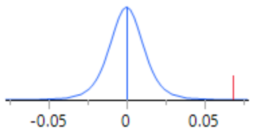


###### Analysis of Variance

| Source | DF | Sum of Squares | Mean Square | F Ratio | Prob > F |
|---|---|---|---|---|---|
| Modif | 1 | 0.00946787 | 0.009468 | 41.0394 | 0.0007* |
| Error | 6 | 0.00138421 | 0.000231 | | |
| C. Total | 7 | 0.01085208 | | | |

###### Means for Oneway Anova

| Level | Number | Mean | Std Error | Lower 95% | Upper 95% |
|---|---|---|---|---|---|
| D-lac | 4 | 0.103919 | 0.00759 | 0.08534 | 0.12250 |
| L-lac | 4 | 0.172723 | 0.00759 | 0.15414 | 0.19131 |

Std Error uses a pooled estimate of error variance

##### Wilcoxon / Kruskal-Wallis Tests (Rank Sums)

| Level | Count | Score Sum | Expected Score | Score Mean | (Mean-Mean0)/Std0 |
|---|---|---|---|---|---|
| D-lac | 4 | 10.000 | 18.000 | 2.50000 | -2.165 |
| L-lac | 4 | 26.000 | 18.000 | 6.50000 | 2.165 |

###### Wilcoxon Two-Sample Test, Normal Approximation

| S | Z | Prob>\|Z\| |
|---|---|---|
| 26 | 2.16506 | 0.0304* |

###### Kruskal-Wallis Test, ChiSquare Approximation

| ChiSquare | DF | Prob>ChiSq |
|---|---|---|
| 5.3333 | 1 | 0.0209* |

Small sample sizes. Refer to statistical tables for tests, rather than large-sample approximations.

H3K56

#### Oneway Analysis of Data By Modif Variant=H3, Site 2=K56

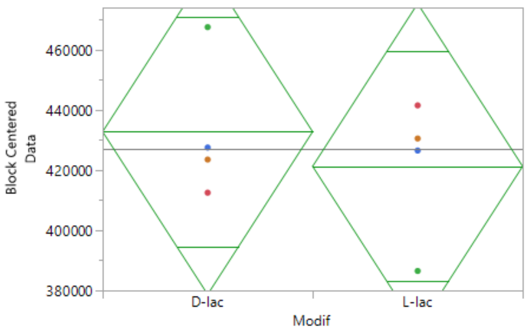


##### Oneway Anova

###### Summary of Fit

| | |
|---|---|
| Rsquare | 0.986129 |
| Adj Rsquare | 0.967634 |
| Root Mean Square Error | 33968.12 |
| Mean of Response | 427000 |
| Observations (or Sum Wgts) | 8 |

###### Pooled t Test

L-lac-D-lac

Assuming equal variances

| | | | |
|---|---|---|---|
| Difference | -11500 | t Ratio | -0.47879 |
| Std Err Dif | 24019 | DF | 3 |
| Upper CL Dif | 64939 | Prob > \|t\| | 0.6648 |
| Lower CL Dif | -87939 | Prob > t | 0.6676 |
| Confidence | 0.95 | Prob < t | 0.3324 |

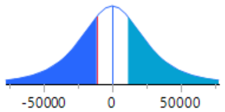


###### Analysis of Variance

| Source | DF | Sum of Squares | Mean Square | F Ratio | Prob > F |
|---|---|---|---|---|---|
| Modif | 1 | 264500000 | 264500000 | 0.2292 | 0.6648 |
| Label | 3 | 2.4582e+11 | 8.194e+10 | 71.0149 | 0.0028* |
| Error | 3 | 3461500000 | 1.1538e+9 | | |
| C. Total | 7 | 2.4954e+11 | | | |

###### Means for Oneway Anova

| Level | Number | Mean | Std Error | Lower 95% | Upper 95% |
|---|---|---|---|---|---|
| D-lac | 4 | 432750 | 16984 | 378699 | 486801 |
| L-lac | 4 | 421250 | 16984 | 367199 | 475301 |

Std Error uses a pooled estimate of error variance

###### Block Means

| Label | Mean | Number |
|---|---|---|
| Abundance 1 | 187500 | 2 |
| Abundance 2 | 677500 | 2 |
| Abundance 3 | 458500 | 2 |
| Abundance 4 | 384500 | 2 |

##### Friedman Rank Test

| Level | Count | Score Sum | Expected Score | Score Mean | (Mean-Mean0)/Std0 |
|---|---|---|---|---|---|
| D-lac | 4 | 6.000 | 6.000 | 1.50000 | 0.000 |
| L-lac | 4 | 6.000 | 6.000 | 1.50000 | 0.000 |

###### 2 - Sample Test, Normal Approximation

| S | Z | Prob>\|Z\| |
|---|---|---|
| 6 | 0.00000 | 1.0000 |

###### 1-Way Test, ChiSquare Approximation

| ChiSquare | DF | Prob>ChiSq |
|---|---|---|
| 0.0000 | 1 | 1.0000 |

Small sample sizes. Refer to statistical tables for tests, rather than large-sample approximations.

Block Label

#### Oneway Analysis of Data By Modif Variant=H3, Lys=K56

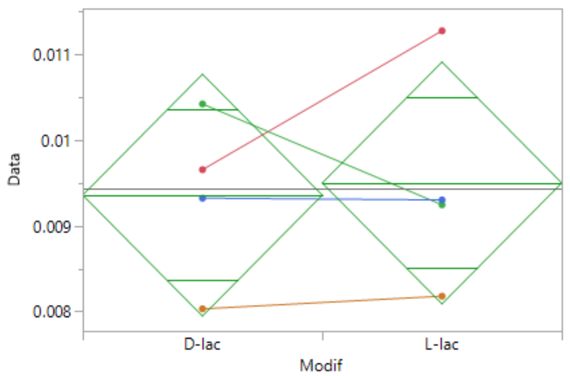


##### Oneway Anova

###### Summary of Fit

| | |
|---|---|
| Rsquare | 0.005075 |
| Adj Rsquare | -0.16075 |
| Root Mean Square Error | 0.001152 |
| Mean of Response | 0.009427 |
| Observations (or Sum Wgts) | 8 |

###### Pooled t Test

L-lac-D-lac

Assuming equal variances

| | | | |
|---|---|---|---|
| Difference | 0.00014 | t Ratio | 0.174938 |
| Std Err Dif | 0.00081 | DF | 6 |
| Upper CL Dif | 0.00214 | Prob > \|t\| | 0.8669 |
| Lower CL Dif | -0.00185 | Prob > t | 0.4334 |
| Confidence | 0.95 | Prob < t | 0.5666 |

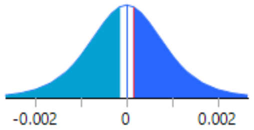


###### Analysis of Variance

| Source | DF | Sum of Squares | Mean Square | F Ratio | Prob > F |
|---|---|---|---|---|---|
| Modif | 1 | 4.06116e-8 | 4.0612e-8 | 0.0306 | 0.8669 |
| Error | 6 | 7.96222e-6 | 1.327e-6 | | |
| C. Total | 7 | 8.00283e-6 | | | |

###### Means for Oneway Anova

| Level | Number | Mean | Std Error | Lower 95% | Upper 95% |
|---|---|---|---|---|---|
| D-lac | 4 | 0.009356 | 0.00058 | 0.00795 | 0.01077 |
| L-lac | 4 | 0.009498 | 0.00058 | 0.00809 | 0.01091 |

Std Error uses a pooled estimate of error variance

##### Wilcoxon / Kruskal-Wallis Tests (Rank Sums)

| Level | Count | Score Sum | Expected Score | Score Mean | (Mean-Mean0)/Std0 |
|---|---|---|---|---|---|
| D-lac | 4 | 19.000 | 18.000 | 4.75000 | 0.144 |
| L-lac | 4 | 17.000 | 18.000 | 4.25000 | -0.144 |

###### Wilcoxon Two-Sample Test, Normal Approximation

| S | Z | Prob>\|Z\| |
|---|---|---|
| 17 | -0.14434 | 0.8852 |

###### Kruskal-Wallis Test, ChiSquare Approximation

| ChiSquare | DF | Prob>ChiSq |
|---|---|---|
| 0.0833 | 1 | 0.7728 |

Small sample sizes. Refer to statistical tables for tests, rather than large-sample approximations.

# H4K77

#### Oneway Analysis of Data By Modif Variant=H4, Site 2=K77

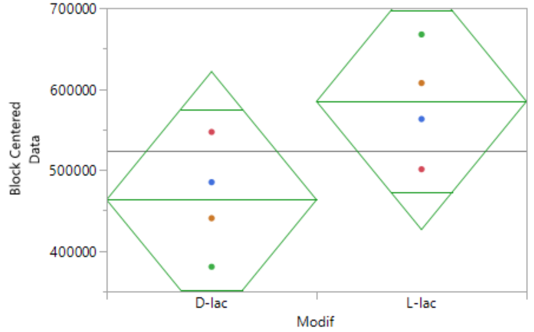


##### Oneway Anova

###### Summary of Fit

| | |
|---|---|
| Rsquare | 0.903023 |
| Adj Rsquare | 0.77372 |
| Root Mean Square Error | 99506.28 |
| Mean of Response | 524000 |
| Observations (or Sum Wgts) | 8 |

###### Pooled t Test

L-lac-D-lac

Assuming equal variances

| | | | |
|---|---|---|---|
| Difference | 121500 | t Ratio | 1.726795 |
| Std Err Dif | 70362 | DF | 3 |
| Upper CL Dif | 345422 | Prob > \|t\| | 0.1827 |
| Lower CL Dif | -102422 | Prob > t | 0.0913 |
| Confidence | 0.95 | Prob < t | 0.9087 |

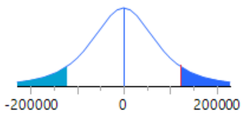


###### Analysis of Variance

| Source | DF | Sum of Squares | Mean Square | F Ratio | Prob > F |
|---|---|---|---|---|---|
| Modif | 1 | 2.9524e+10 | 2.952e+10 | 2.9818 | 0.1827 |
| Label | 3 | 2.4708e+11 | 8.236e+10 | 8.3178 | 0.0577 |
| Error | 3 | 2.9704e+10 | 9.9015e+9 | | |
| C. Total | 7 | 3.063e+11 | | | |

###### Means for Oneway Anova

| Level | Number | Mean | Std Error | Lower 95% | Upper 95% |
|---|---|---|---|---|---|
| D-lac | 4 | 463250 | 49753 | 304913 | 621587 |
| L-lac | 4 | 584750 | 49753 | 426413 | 743087 |

Std Error uses a pooled estimate of error variance

###### Block Means

| Label | Mean | Number |
|---|---|---|
| Abundance 1 | 276000 | 2 |
| Abundance 2 | 734500 | 2 |
| Abundance 3 | 635000 | 2 |
| Abundance 4 | 450500 | 2 |

##### Friedman Rank Test

| Level | Count | Score Sum | Expected Score | Score Mean | (Mean-Mean0)/Std0 |
|---|---|---|---|---|---|
| D-lac | 4 | 5.000 | 6.000 | 1.25000 | -0.935 |
| L-lac | 4 | 7.000 | 6.000 | 1.75000 | 0.935 |

###### 2 - Sample Test, Normal Approximation

| S | Z | Prob>\|Z\| |
|---|---|---|
| 7 | 0.93541 | 0.3496 |

###### 1-Way Test, ChiSquare Approximation

| ChiSquare | DF | Prob>ChiSq |
|---|---|---|
| 1.0000 | 1 | 0.3173 |

Small sample sizes. Refer to statistical tables for tests, rather than large-sample approximations.

Block Label

#### Oneway Analysis of Data By Modif Variant=H4, Lys=K77

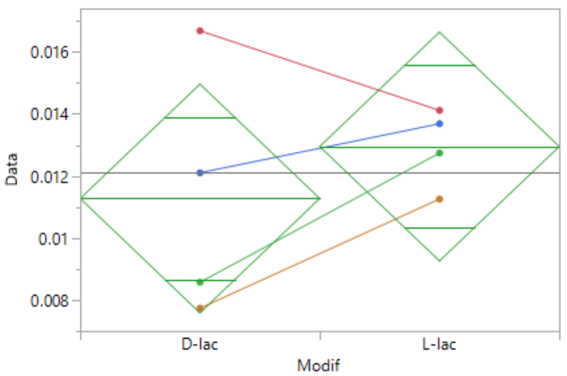


##### Oneway Anova

###### Summary of Fit

| | |
|---|---|
| Rsquare | 0.093306 |
| Adj Rsquare | -0.05781 |
| Root Mean Square Error | 0.003016 |
| Mean of Response | 0.012112 |
| Observations (or Sum Wgts) | 8 |

###### Pooled t Test

L-lac-D-lac

Assuming equal variances

| | | | |
|---|---|---|---|
| Difference | 0.00168 | t Ratio | 0.785779 |
| Std Err Dif | 0.00213 | DF | 6 |
| Upper CL Dif | 0.00689 | Prob > \|t\| | 0.4619 |
| Lower CL Dif | -0.00354 | Prob > t | 0.2309 |
| Confidence | 0.95 | Prob < t | 0.7691 |

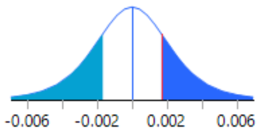


###### Analysis of Variance

| Source | DF | Sum of Squares | Mean Square | F Ratio | Prob > F |
|---|---|---|---|---|---|
| Modif | 1 | 0.00000561 | 5.6146e-6 | 0.6174 | 0.4619 |
| Error | 6 | 0.00005456 | 9.0933e-6 | | |
| C. Total | 7 | 0.00006017 | | | |

###### Means for Oneway Anova

| Level | Number | Mean | Std Error | Lower 95% | Upper 95% |
|---|---|---|---|---|---|
| D-lac | 4 | 0.011274 | 0.00151 | 0.00759 | 0.01496 |
| L-lac | 4 | 0.012950 | 0.00151 | 0.00926 | 0.01664 |

Std Error uses a pooled estimate of error variance

##### Wilcoxon / Kruskal-Wallis Tests (Rank Sums)

| Level | Count | Score Sum | Expected Score | Score Mean | (Mean-Mean0)/Std0 |
|---|---|---|---|---|---|
| D-lac | 4 | 15.000 | 18.000 | 3.75000 | -0.722 |
| L-lac | 4 | 21.000 | 18.000 | 5.25000 | 0.722 |

###### Wilcoxon Two-Sample Test, Normal Approximation

| S | Z | Prob>\|Z\| |
|---|---|---|
| 21 | 0.72169 | 0.4705 |

###### Kruskal-Wallis Test, ChiSquare Approximation

| ChiSquare | DF | Prob>ChiSq |
|---|---|---|
| 0.7500 | 1 | 0.3865 |

Small sample sizes. Refer to statistical tables for tests, rather than large-sample approximations.

**Section 5:** Reason why H4K31la could not be quantified in the four biological replicates of mouse testis histones

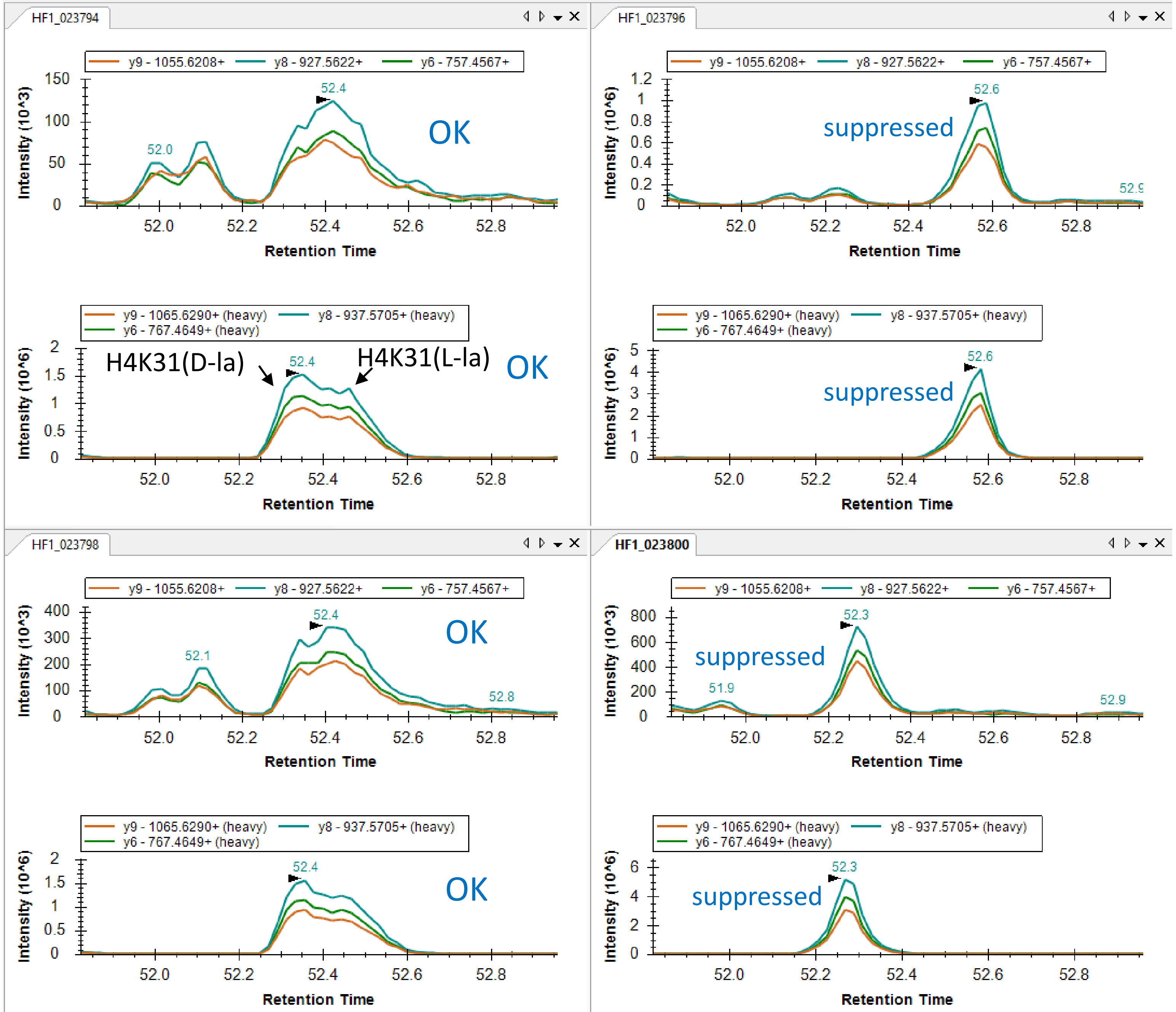


Detection of MS/MS signals for heavy and endogenous DNIQGITK31(la)PAIR is hampered by another abundant species in two analyses out of the four.

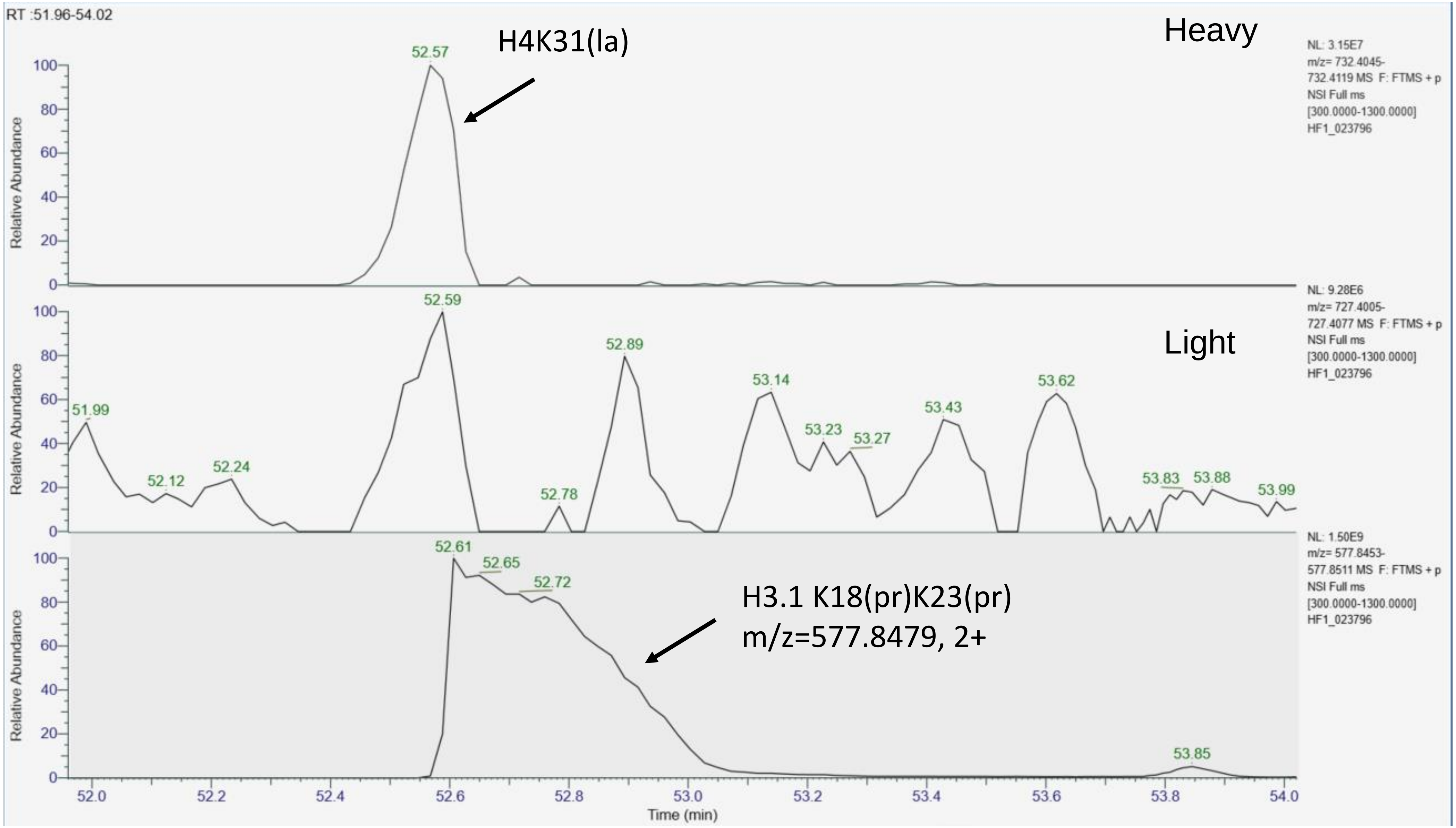


The interfering peptide is probably the highly intense (and abundant) peptide H3 K18prQLATK23prAAR.

#### Section 6: MS2 signals for endogenous and heavy H3K79la-containing peptides show that the endogenous peptide essentially matches the D-lactylated synthetic peptide

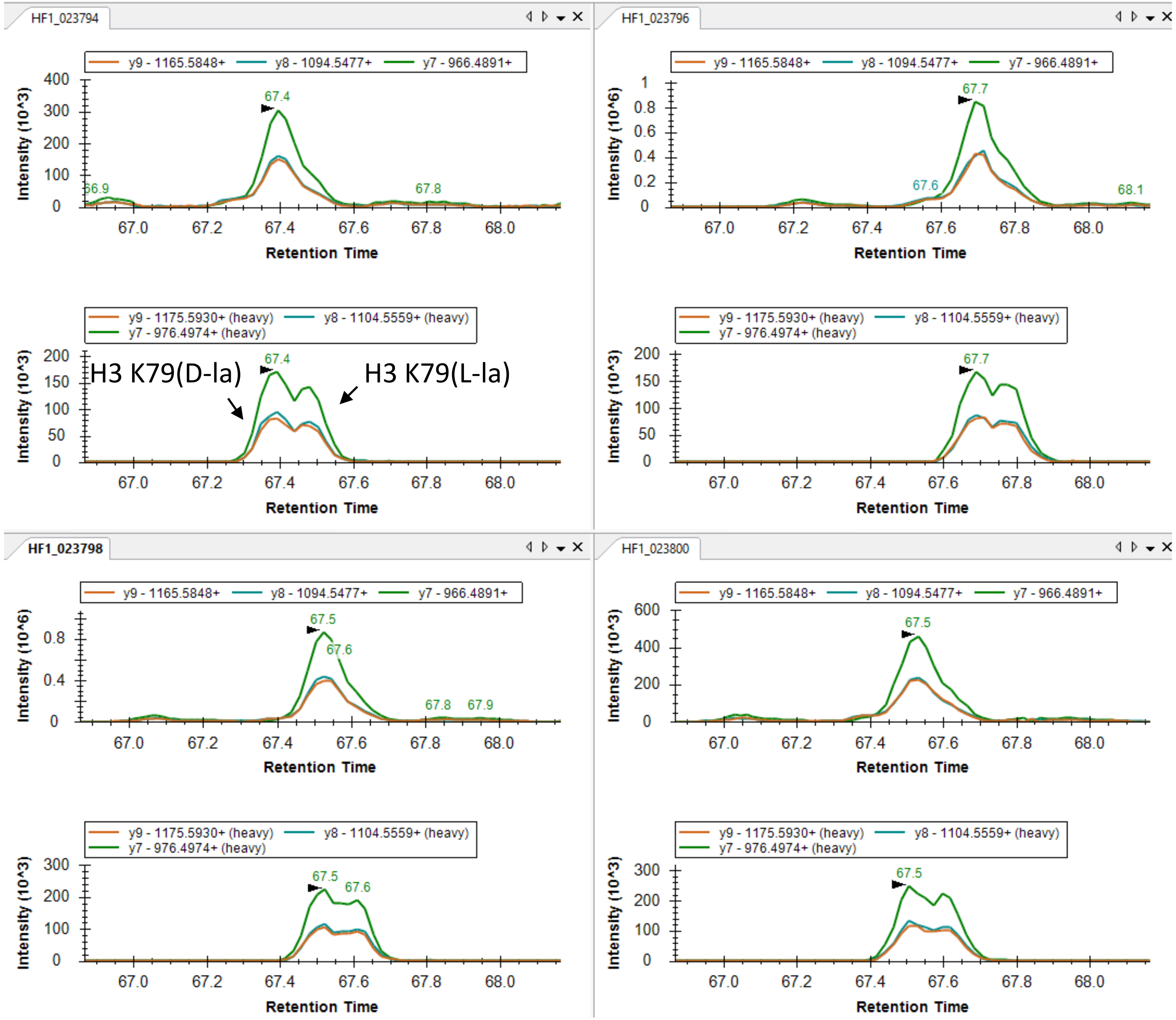

**Section 7:** Comparison of the retention times of acetylated peptides and their lactylated counterparts

#### Retention time of synthetic acetylated peptides versus D-lactylated peptides of H3 and H4

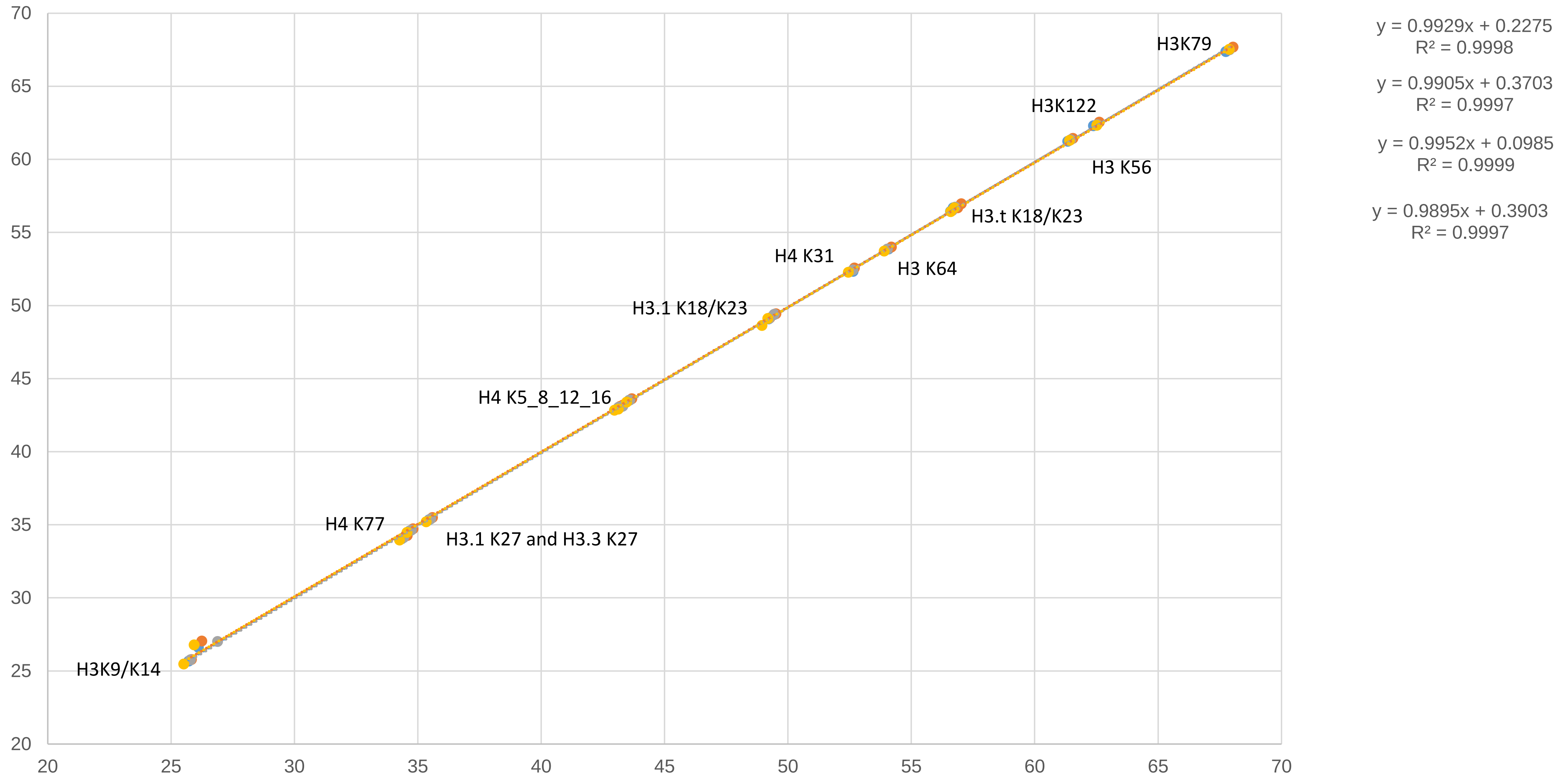


Four technical replicate injections were performed.

#### Retention time of synthetic acetylated peptides versus L-lactylated peptides of H3 and H4

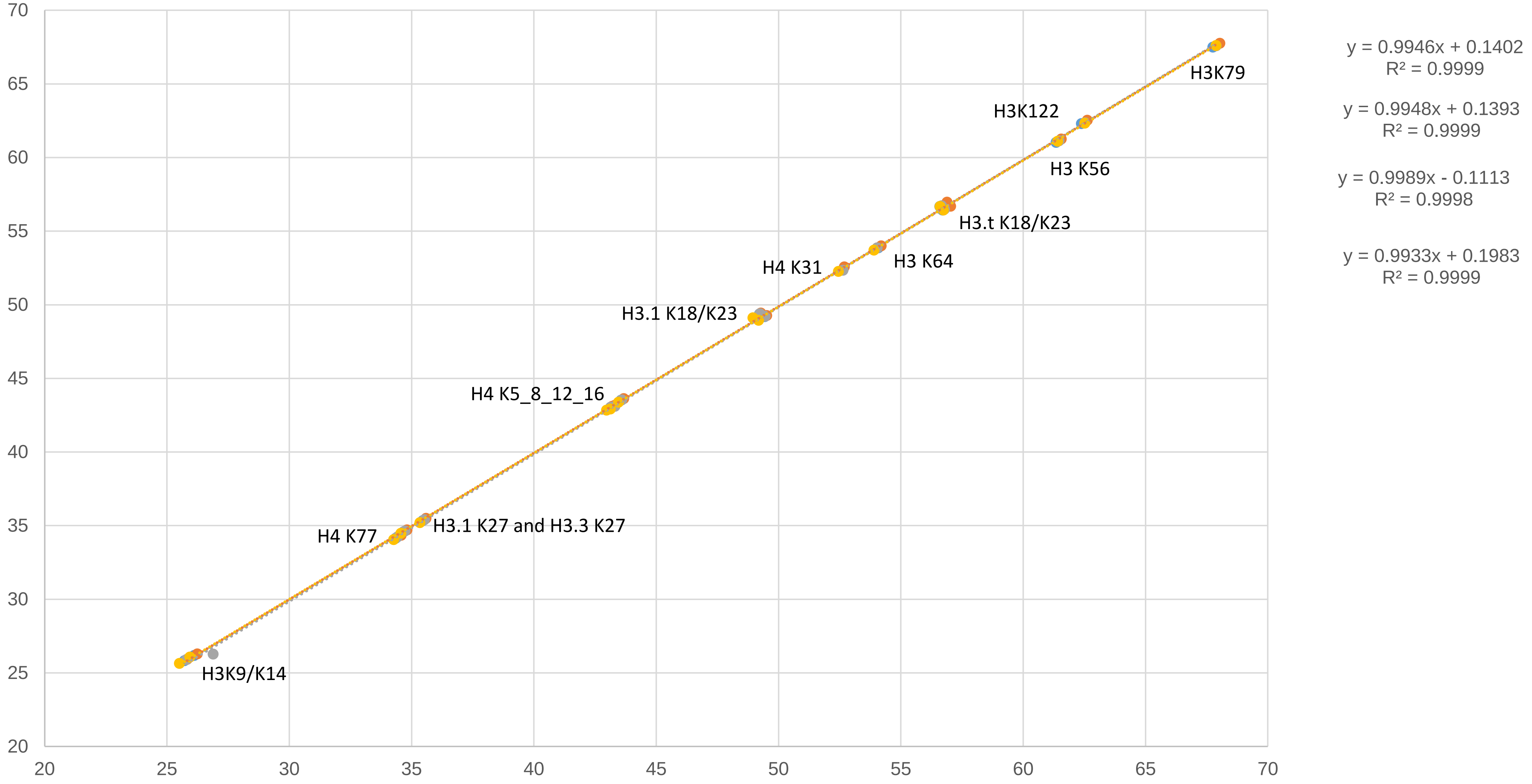


Four technical replicate injections were performed.

**Section 8:** Comparison of abundance of acetylated peptides and their lactylated counterparts on H4

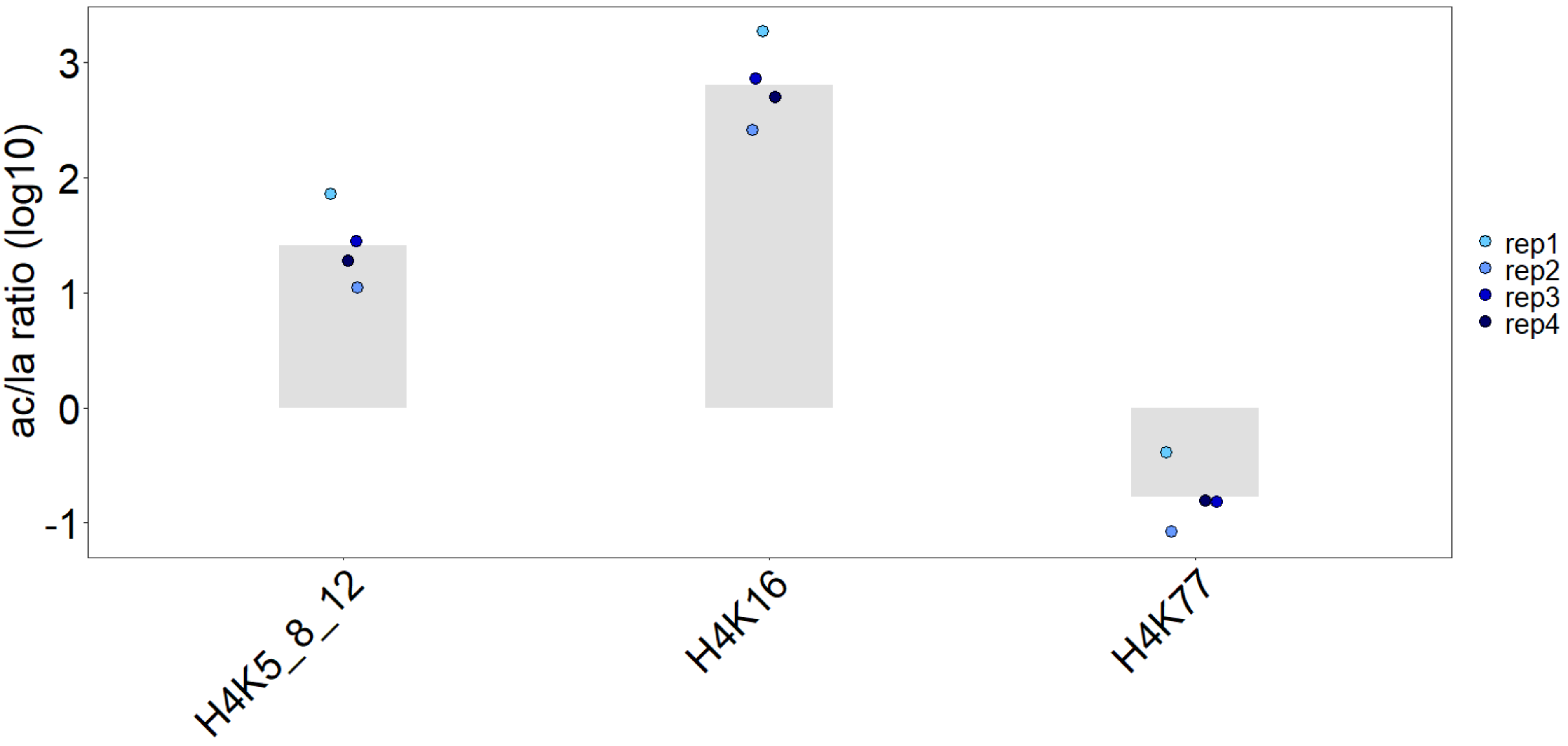

**Section 9:** Verification of the absence of the isomeric structure with lactyl, N-ε-(carboxyethyl)

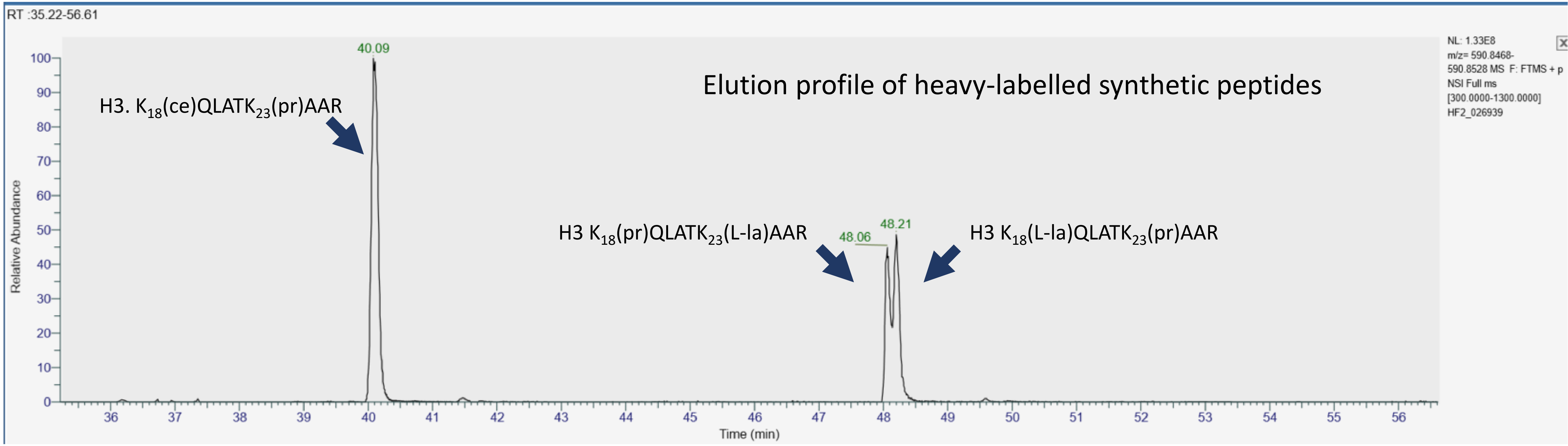
RT :35.22-56.61
Elution profile of heavy-labelled synthetic peptides
H3. $K_{18}$(ce)QLATK$_{23}$(pr)AAR
H3 $K_{18}$(pr)QLATK$_{23}$(L-la)AAR
H3 $K_{18}$(L-la)QLATK$_{23}$(pr)AAR
40.09
48.06
48.21
NL: 1.33E8
m/z= 590.8468-590.8528 MS F: FTMS + p NSI Full ms [300.0000-1300.0000]
HF2_026939
Relative Abundance
Time (min)

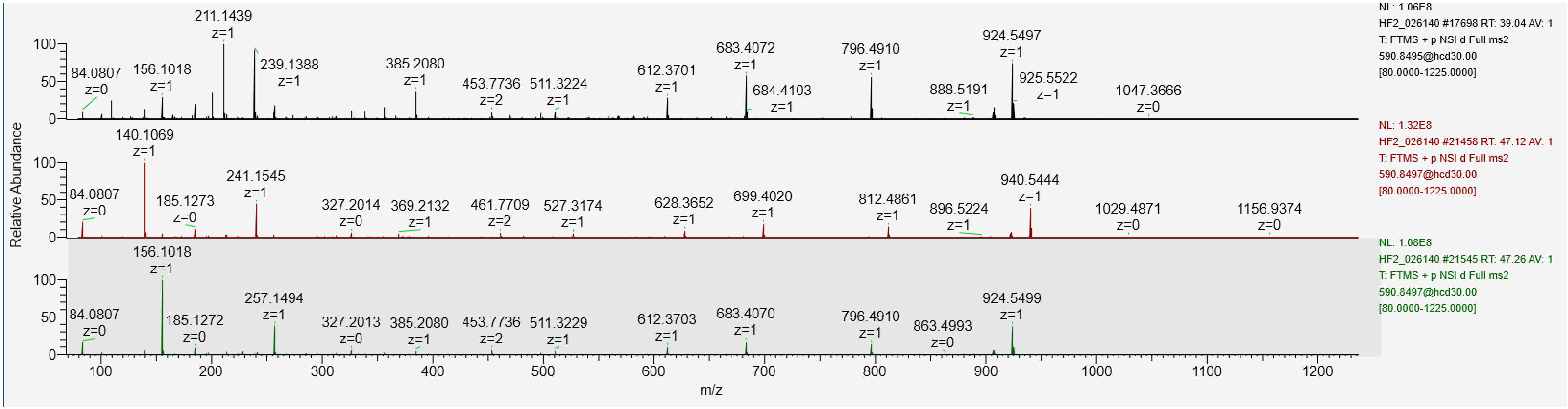
NL: 1.06E8
HF2_026140 #17698 RT: 39.04 AV: 1
T: FTMS + p NSI d Full ms2 590.8495@hcd30.00 [80.0000-1225.0000]
NL: 1.32E8
HF2_026140 #21458 RT: 47.12 AV: 1
T: FTMS + p NSI d Full ms2 590.8497@hcd30.00 [80.0000-1225.0000]
NL: 1.08E8
HF2_026140 #21545 RT: 47.26 AV: 1
T: FTMS + p NSI d Full ms2 590.8497@hcd30.00 [80.0000-1225.0000]
Relative Abundance
m/z

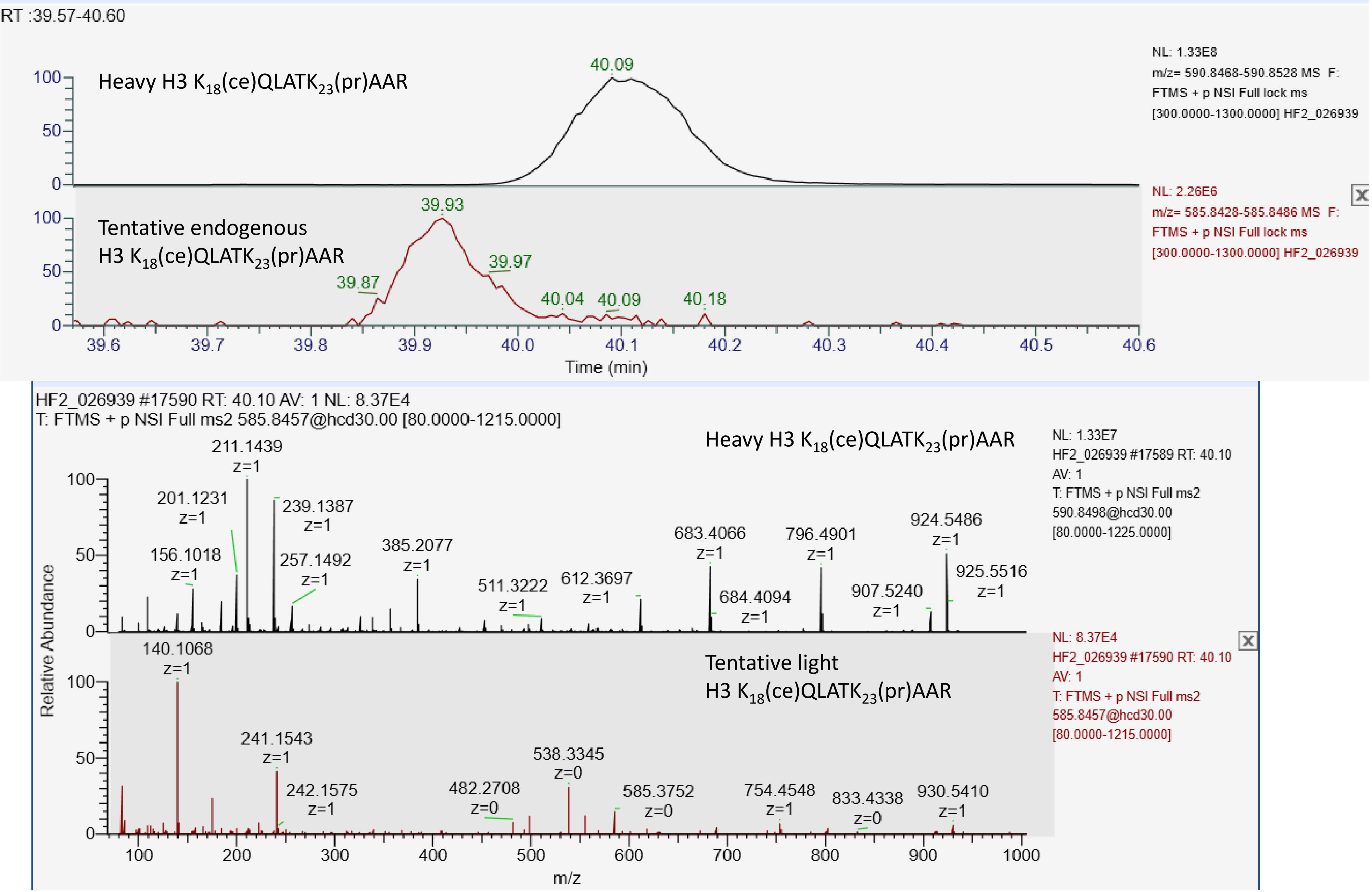
RT :39.57-40.60
Heavy H3 K18(ce)QLATK23(pr)AAR
NL: 1.33E8
m/z= 590.8468-590.8528 MS F: FTMS + p NSI Full lock ms [300.0000-1300.0000] HF2_026939
Tentative endogenous H3 K18(ce)QLATK23(pr)AAR
NL: 2.26E6
m/z= 585.8428-585.8486 MS F: FTMS + p NSI Full lock ms [300.0000-1300.0000] HF2_026939
Time (min)
HF2_026939 #17590 RT: 40.10 AV: 1 NL: 8.37E4
T: FTMS + p NSI Full ms2 585.8457@hcd30.00 [80.0000-1215.0000]
Heavy H3 K18(ce)QLATK23(pr)AAR
NL: 1.33E7
HF2_026939 #17589 RT: 40.10 AV: 1
T: FTMS + p NSI Full ms2 590.8498@hcd30.00 [80.0000-1225.0000]
Tentative light H3 K18(ce)QLATK23(pr)AAR
NL: 8.37E4
HF2_026939 #17590 RT: 40.10 AV: 1
T: FTMS + p NSI Full ms2 585.8457@hcd30.00 [80.0000-1215.0000]
Relative Abundance
m/z
